# Single-Cell Resolution of Cellular Damage Illuminates Disease Progression

**DOI:** 10.1101/2025.02.20.639269

**Authors:** Tsimafei Padvitski, Paula Unger Avila, He Chen, Cem Özel, Fabian Braun, Marie-Kristin Kroll, Christian Vossen, Peter Wienand, Linus Butt, David Unnersjö-Jess, Heike Göbel, Christoph Kuppe, Roman-Ulrich Müller, Pål O. Westermark, Paul T. Brinkkötter, Bernhard Schermer, Thomas Benzing, F. Thomas Wunderlich, Martin Kann, Andreas Beyer

## Abstract

Degenerative diseases feature progressive accumulation of cellular damage, leading to impaired organ function. Capturing such gradual decline of individual cells *in vivo* remains challenging. Here, we present a universal framework for quantifying damage at single-cell resolution, to uncover molecular trajectories of degeneration. This method enables the detection of progressive damage within cell populations under physiological and pathological conditions. We developed the Podocyte Damage Score (PDS) and Hepatocyte Damage Score (HDS) to monitor deterioration in kidney glomerulosclerosis and liver steatosis. The application of these scores to murine and human datasets quantified cellular damage across disease models and distinguished varying degrees of damage even in unperturbed samples. The PDS revealed circadian gene expression dysregulation as a consequence of podocyte injury, while the HDS identified a threshold of hepatocyte damage leading to senescence and metabolic dysfunction. The approach provides a scalable tool for decoding disease progression and identifying therapeutic targets across degenerative disorders.

## Introduction

Degenerative diseases, such as chronic kidney disease (CKD) and non-alcoholic fatty liver disease (NAFLD), represent a growing global health burden. The global prevalence of CKD in 2010 was estimated at ∼11%,^1^ and non-alcoholic fatty liver disease (NAFLD) was considered to affect ∼25% of the global population in 2016.^2^ These conditions are characterized by the gradual accumulation of cellular damage, leading to progressive loss of tissue function and contributing to severe complications, including organ failure and cancer. Despite their prevalence, the molecular mechanisms that drive the onset and progression of degenerative diseases remain poorly understood. This knowledge gap hampers the development of targeted diagnostics and effective therapies. A central challenge in studying degenerative diseases lies in capturing the slow and heterogeneous progression of cellular dysfunction. Disease onset often involves subtle, progressive changes within specific cell populations that are difficult to detect using conventional bulk analyses or histopathology. Comparing molecular differences between cases and controls cannot map out the continuum of cellular damage accumulation that is characteristic of degenerative diseases. Moreover, existing animal models frequently mimic rare, monogenic forms of disease and fail to reflect the complexity of sporadic, multifactorial conditions in humans.

In CKD, the progressive decline of kidney function almost invariably involves damage to podocytes, terminally differentiated epithelial cells within the kidney glomeruli, the filtration units of the kidney. However, recent large-scale transcriptomic approaches in humans have shown that changes in glomerular transcriptional programs, while depicting individual aspects of CKD secondary changes, cannot explain the gradual progress of podocyte degeneration.^3–7^ Similarly, NAFLD involves a complex interplay between liver cell types, while its defining criterion is damage to the predominant hepatic epithelial cell type, the hepatocyte.^2^ Hepatocytes are key to metabolic functions of the liver and are tightly connected to inflammatory and immune responses. However, pathways of gradual degeneration and ultimately carcinogenesis are still elusive.^8,9^

Advances in single-cell RNA sequencing (scRNA-seq) and spatial transcriptomics have revolutionized our ability to profile gene expression at cellular resolution, offering new opportunities to dissect disease processes at unprecedented detail and study the function of disease-associated genes,^10^ and regulatory events.^11^ However, current analytical approaches are limited in their capacity to quantify gradual cellular deterioration. The primary challenge in interpreting scRNA-seq data stems from the absence of information regarding the physiological status of individual cells. Whereas cell types can faithfully be recovered from scRNA-seq data, deriving quantitative information about cellular damage remains an unsolved problem. Traditional clustering and differential expression analyses primarily capture discrete cell states, overlooking continuous damage progression. Pseudo-time trajectory methods, while useful for modeling developmental processes, often struggle to distinguish between physiological variability and pathological decline. Hence, it remains elusive how to map out the entire continuum of disease-associated cellular state changes from healthy to severely damaged.

To address this gap, we introduce a novel computational framework for quantifying cellular damage at single-cell resolution. By developing cell-type-specific damage scores - namely, the Podocyte Damage Score (PDS) and Hepatocyte Damage Score (HDS) - we systematically track molecular deterioration in kidney and liver disease models. These scores, for the first time, represent *in vivo* states of cells in chronic disease over extended time periods and enable the identification of molecular pathways associated with progressive damage, even within rare cell-types or early-stage diseased cells. Applying this framework across diverse murine models and human datasets, we uncover conserved mechanisms of cellular injury, including disrupted circadian rhythm in podocytes and senescence-driven metabolic dysfunction in hepatocytes.

Our approach is transferrable to other chronic diseases and offers a powerful tool for mapping cellular damage trajectories, providing critical insights into disease progression as well as potential therapeutic targets across degenerative conditions.

## Results

### Damage scores

Cellular damage can be defined in different ways. Here, we operationally define ‘cellular damage’ as a pathological cellular state that is not specific to any disease-causing mechanism. Instead, damage is a common, pathological cellular state resulting from a broader spectrum of disease entities, disease-causing mechanisms, and/or a loss of homeostasis. Such cell-type specific cellular damage states are well established in histopathology. Here, we develop an approach for the molecular characterization of such states. To demonstrate the general applicability of our concept, we developed two single-cell damage scores: one for podocytes (Podocyte Damage Score, PDS), to track damage during the progression of CKD, and a second for hepatocytes (Hepatocyte Damage Score, HDS), to explore progressive damage in chronic liver pathologies such as NAFLD. In the following, we first motivate the selection criteria for damage marker genes and subsequently validate both scores using mouse and human single-cell and spatial omics data. Finally, we present applications of first the PDS and then the HDS to demonstrate the added value and novel mechanistic insight gained from these scores.

The key notion of our approach is to identify marker genes whose expression is associated with increasing damage across a wide range of disease models (Figure 1A). By combining diverse disease models in the selection process, we ensure that the marker genes are not just markers for one specific intervention, but report on damage of the target cell type in general. The damage scores combine information from dozens of marker genes, which increases robustness and reduces the impact of single (potentially noisy) outlier genes. Further, our approach ensures that selected marker genes are expressed in the target cell type, thereby excluding for example genes specific to tissue-resident immune cells that might be indicative of inflammation. Using gene expression data from 37 datasets covering 9 podocytopathy models and ∼250 samples covering 13 liver disease models (Table S1), we first ranked genes by their p-values for differential expression in each of these models (always comparing the diseased/experimental condition versus the respective control). Those ranks were subsequently averaged across all disease models per cell type to obtain one global gene ranking for podocytes and one for hepatocytes. Next, we applied two additional filters: first, we removed genes that were not expressed in the target cell type (i.e. podocytes or hepatocytes; Methods). Second, we only kept genes with a consistent direction of change (same sign) in at least 75% of all studies. Finally, we chose the top *n* genes as damage marker genes for each of the two cell types. To quantify the damage of an individual cell (or bulk sample), we tested how extreme the expression of PDS/HDS marker genes was in a given cell using AUCell.^12^ Genes with positive and negative log-fold changes were combined considering the direction of effects (Methods), resulting in an algorithm capable of scoring cellular damage at single cell resolution (Figure 1B,E).

**Figure 1:**
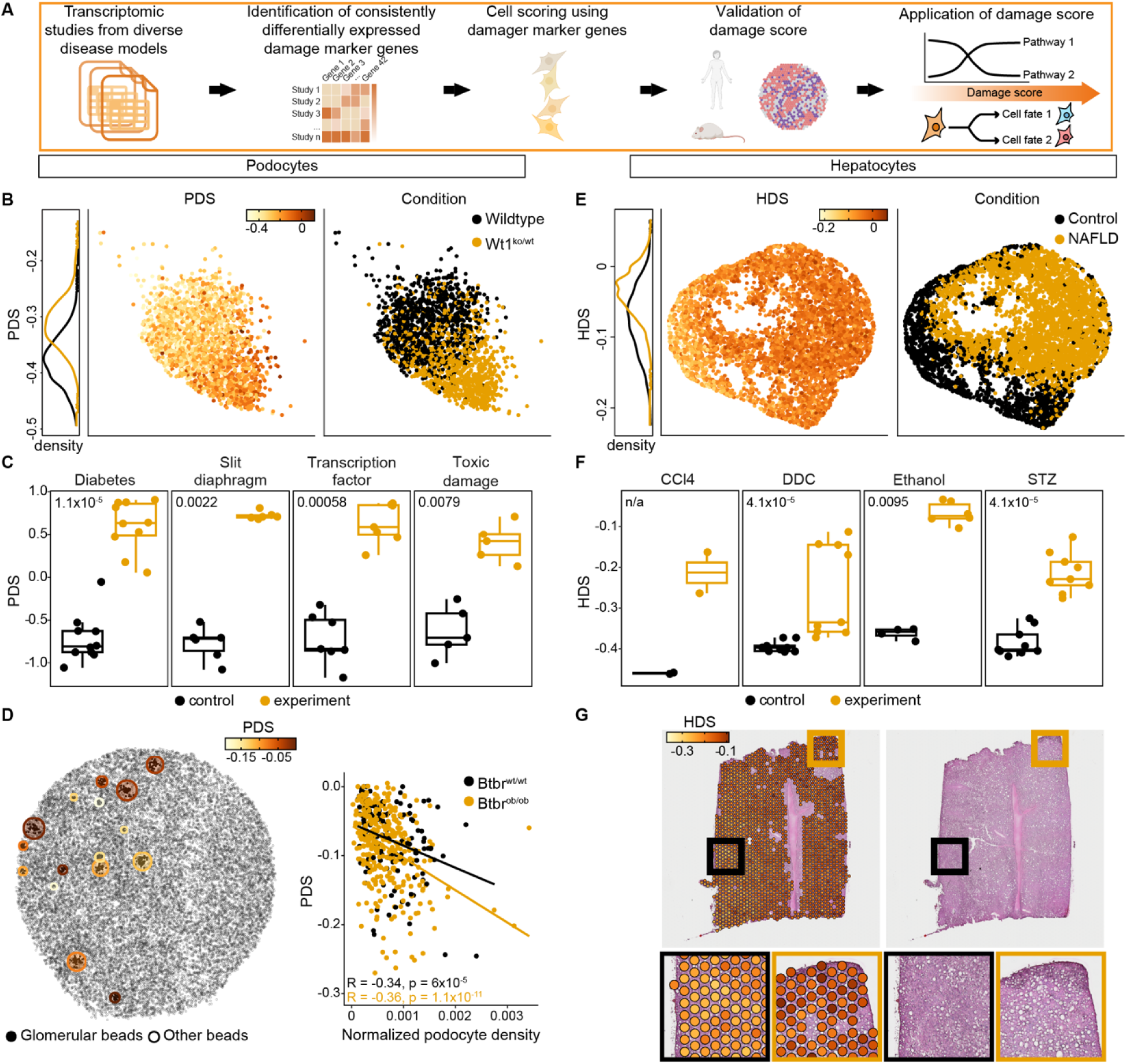
A strategy to quantify damage at the single cell level based on transcriptomic data. A. Robust damage marker genes are identified by compiling transcriptomic data from a great diversity of disease models. Top genes that change consistently across models are selected as damage markers. Damage of individual cells is quantified using a rank-based approach. After validation, the damage scores are used to analyze the molecular progression of chronic degenerative diseases *in vivo*. B. Application of the podocyte damage score (PDS) to podocytes from snRNA-seq data of the Wt1^ko/wt^ mouse model (a podocyte damage model). Left UMAP is colored by PDS (see color scale), right UMAP is colored by condition. The PDS density plot (left) shows the distribution of PDS values separated by experimental condition (same colors as UMAP on the right). C. Leave-one-model-out cross validation of the PDS. All data of the indicated disease models were left out during the creation of a variant damage signature and used afterwards for testing PDS in the indicated disease models (see Table S1 for grouping). Each dot represents the average damage score per sample. Scores are scaled and centered per study. P-values of Wilcoxon rank test (experimental versus control conditions) are shown. The boxplots show first (Q1), second (Q2), and third quartiles (Q3) of distribution. Whiskers represent Q1 - 1.5 x interquartile range (IQR) and Q3 + 1.5 x IQR. D. Spatial validation of the PDS using kidney Slide-seqV2 spatial transcriptomics data on diabetic Btbr^ob/ob^ and control kidneys (GSE190094^45^). Representative section of a diabetic mouse kidney is shown. Circles represent glomerular areas and are colored by the mean PDS calculated for podocyte beads (as determined in Marshall et al.^45^) within one glomerulus. The PDS correlated with glomerular size (higher PDS for larger glomeruli), albeit without correction for tissue sectioning-related artifacts. Right plot shows PDS measurements normalized for podocyte density after correcting for area variance caused by tissue sectioning and reveals increased podocyte damage corresponding to hyperfiltration in low podocyte density settings such as diabetes or glomerular hypertrophy (see Figure S3A for density calculation). E. Application of the hepatocyte damage score (HDS) to hepatocytes of snRNA-seq data from 24-week-old mice fed with either a standard diet (control) or a NAFLD-inducing diet (NAFLD).^9,48^ Left UMAP is colored by HDS (see color scale), right UMAP is colored by diet. HDS density plot (left) shows the distribution of HDS values separated by experimental condition (same colors as UMAP on the right). F. Leave-one-model-out cross validation of the HDS. Procedure is the same as in C. Each dot represents the HDS of one sample. CCl4: hepatic toxic injury by CCl4 injection, DDC: hepatotoxin-induced cholestatic liver injury via 3,5-diethoxycarbonyl-1,4-dihydrocollidine, Ethanol: alcoholic liver injury via ethanol administration, STZ: streptozotocin induced liver injury. Boxplots show first (Q1), second, and third quartiles (Q) of distribution. Whiskers represent Q1 - 1.5 × IQR and Q3 + 1.5 × IQR. P-values of Wilcoxon rank test (experimental versus control conditions) are shown. N/a: statistics not applicable due to n = 2. G. Spatial validation of HDS using 10X Visium mouse spatial transcriptomic data^9,48^. Liver tissue section of a mouse with NAFLD. Left: HDS computed for each spot that passed filtering in spatial transcriptomics data. Right: Corresponding H&E staining. Indicated areas are enlarged as insets, in which HDS correlates with the degree of lipid accumulation. Black boxes indicate area with little steatosis and low HDS, yellow boxes indicate area with marked steatosis and high HDS. For additional data see Figure S4B.

#### Damage scores robustly characterize cellular states

To establish the robustness of our approach, we first confirmed that varying the number of top genes used for the damage scores had little influence on the outcome. Varying the number of maker genes between 30 to 200 genes only minimally affected the scores’ ability to distinguish (diseased) experimental samples from (healthy) control samples (Figure S1A-D). Likewise, randomizing parts of the input data had little influence on the ability of the damage scores to distinguish healthy from diseased samples. Even when randomizing (shuffling) up to 70% of all input studies, the resulting damage scores were still able to distinguish experimental from control samples in disease models that were not used for developing the damage scores (Figure S1E,F). The choice of the top *n* genes used for computing the damage scores is, to an extent, arbitrary. We decided to continue all subsequent work with the top 42 marker genes for each, PDS and HDS (Table S2), based on assessing the statistical power to separate diseased from healthy samples (Figure S1B,D). Next, we confirmed that the PDS and HDS were indeed cell-type specific. By analyzing snRNA-seq data obtained from glomeruli and liver, we confirmed that only the targeted cell-type showed significant variation with damage (Figure S1G,H). Furthermore, available glomerular proteomic data^13^ confirmed that PDS marker proteins were more abundant in podocytes compared to mesangial and glomerular endothelial cells (Figure S2A).

A central aim of our study was to develop scores that measure cellular damage across disease conditions instead of reflecting molecular adaptations in specific disease models. We tested this notion using a ‘leave one model out’ cross validation: first we removed all samples of one similar group of disease models from the training data, then performed the procedure described above to determine damage marker genes on the remaining samples and, finally, tested the resulting damage scores on the left-out disease model samples. In each of the cross-validation rounds, both the PDS and HDS were able to distinguish experimental samples from control samples in the unseen disease models (Figure 1C, 1F and Figure S2B-S2E). Altogether, these results demonstrated that the damage scores are remarkably robust against diverse variations in the data and that they capture generic molecular changes operating across disease models in a cell-type specific fashion.

#### Damage coincides with declining cellular identity

Most marker genes for PDS and HDS were downregulated with increasing damage (PDS: 39/42, HDS 30/42, Table S2), suggesting a loss of cell type-specific gene expression. Downregulated PDS markers included genes with established key roles in podocyte biology, such as genes coding for the slit diaphragm-associated regulators Magi2, Thsd7a, Robo2, and Ddn.^14–16^ Further, mutations in the PDS genes Plce1, Wt1, Magi2, and Ptpro are known to cause podocytopathies in humans.^14,17–19^ Other PDS genes with established function in podocytes include the transcription factors (TF) Mafb and Dach1, as well as Sulf1, Cldn5, Optn, Pard3b, Enpep, and Sema3g.^20–25^ Overall, PDS genes were enriched for pathways with high relevance to podocyte biology (Table S2, Figure S2F).

Likewise, many HDS marker genes downregulated in the disease condition are involved in hepatocyte-specific functions such as bile acid metabolism (e.g. Cyp7b1^26^), detoxification (e.g. Selenbp2^27^) and factors secreted into the blood (e.g. Serpinc1 i.e., antithrombin and C8b).^28^ Further, downregulated HDS marker genes are involved in the maintenance of cellular homeostasis, for example, maintaining genomic stability (e.g. Mcm10^29^), response to oxidative stress (e.g., Mpv17l^30^), and immunoregulation (e.g. Ddt^31^). Other downregulated HDS genes have previously been associated with the progression of liver disease (e.g. Nudt7^32^, Hes6^33^), as well as with impaired liver regeneration (e.g. Lifr^34^, and Tm7sf2^35^, Table S2).

Upregulated HDS genes were active in fatty acid uptake (e.g. Cd36^36^), suppression of gluconeogenesis (e.g. Nrg1^37^), and lipid transport (e.g. Apoa4^38^), and were significantly enriched for pathways relevant to cell-adhesion and wound healing processes, indicating responses to tissue damage (Tables S2-S3). Examples are Rtn4, which activates TGFβ-mediated fibrosis,^39^ as well as Iqgap1, which has already been shown to positively correlate with damage. In line with this evidence, silencing Iqgap1 ameliorates hepatic fibrogenesis,^40^ and suppresses liver cancer.^41^ Taken together, these results suggest that cellular damage in podocytes and hepatocytes proceeds in conjunction with cell identity loss.^42,43^

Lastly, we checked podocyte and liver damage markers for associations with known GWAS variants (see Methods). Eight PDS and ten HDS marker genes were associated with kidney and liver disease-related GWAS traits, respectively (Table S2). A significant enrichment for GWAS variants was detected in the gene bodies of podocyte damage marker genes (p = 0.031). This enrichment was enhanced upon inclusion of flanking, non-coding genomic regions (p = 0.002, Fisher’s exact test). While HDS marker genes were not significantly enriched for GWAS variants (Fisher’s exact test p = 0.1 for both tests as described above), they showed a trend towards preferentially containing genes that were associated with liver disease.

### PDS and HDS report on pathologic cellular states

#### Morphological validation

To validate the association of the PDS with histologically detectable damage, we used two morphological features: first, we used glomerular size, because glomerular hyperfiltration such as in diabetic kidney disease results in glomerular hypertrophy and consecutive podocyte stress, as podocytes are unable to proliferate for compensation.^44,45^ Thus, larger glomeruli with lower podocyte densities are indicative of greater podocyte damage. We used spatial transcriptomic data from diabetic and control mouse kidneys to correlate glomerular podocyte density and glomerular PDS.^45^ We observed a significant correlation (p < 1x10^-4^ in controls, p < 1x10^-10^ in diabetic mice) between the two measures (Figure 1D, Figure S3A), showing that the PDS captures within sample variability of cellular damage. Importantly, PDS indicates that glomerular size in wildtype mice is associated with podocyte stress presumably attributable to increased filtration load. Second, we used the slit diaphragm (SD) length, which is a proxy for damage to the podocyte filtration barrier, with shorter length indicating greater damage.^46^ We used STED imaging of SDs to assess the protein expression of three PDS signature members localizing to the SD, namely THSD7A, CLDN5, and EPHB1. This confirmed decreased (EPHB1, THSD7A) or redistributed signals (CLDN5) with shortened SDs in podocytopathy models as compared to controls, which is consistent with greater PDS values indicating greater podocyte damage (Figure S3B-S3C). Further, SD length quantification in three mouse models revealed a significant negative correlation of SD length with the PDS, i.e. models with larger PDS signals exhibited elevated morphological signs of damage (Figure S3D).

In chronic liver disease, a unifying, histologically detectable and locally heterogeneous feature of many disease conditions including viral hepatitis, AFLD, NAFLD, and NASH, is the ectopic formation and accumulation of lipid droplets in hepatocytes (also called liver steatosis).^47^ Therefore, we assessed the local accumulation of lipid droplets in hepatocytes as a proxy for accumulated hepatocyte damage. Using spatial transcriptomic data from a cohort of NAFLD mice and healthy controls^9,48^, we confirmed that the HDS captured within sample variation of liver tissue damage. First, we observed significantly lower average HDS in healthy control liver sections when compared to liver sections from NAFLD mice (Figure S4A). Second, within the same tissue section from NAFLD mice, areas with more abundant fat droplets had greater HDS than less lipid-laden areas (Figure 1G, Figure S4B). Taken together, these results confirm that the PDS and HDS are molecular measures of histologically detectable damage.

#### Functional validation

To test if the PDS also reflects functional decline of podocytes, we correlated the PDS with albuminuria measurements in mice. The albumin/creatinine ratio (ACR) quantifies excess albumin in the urine and is a quantitative and established measure of podocyte damage.^46,49^ We analyzed 12 mouse studies with combined bulk glomerular transcriptomic and ACR measurements and found a significant positive correlation (Spearman = 0.45, p = 0.017) between PDS and albuminuria (Figure S3E). The correlation was even stronger (Spearman = 0.78, p = 1.8x10^-6^) for novel snRNA-seq data generated within this study (Figure 2A, Figure S7). Here, we aggregated PDS across podocytes within each sample and then correlated these pseudo-bulk PDS with albuminuria (Figure 2A). Taken together, these results indicate that the PDS quantitatively reflects functional decline of podocytes. In some contexts, the PDS may even complement albuminuria as a surrogate measure of podocyte damage, because unlike albuminuria it is measurable outside of a functioning organism and even in single cells.

**Figure 2:**
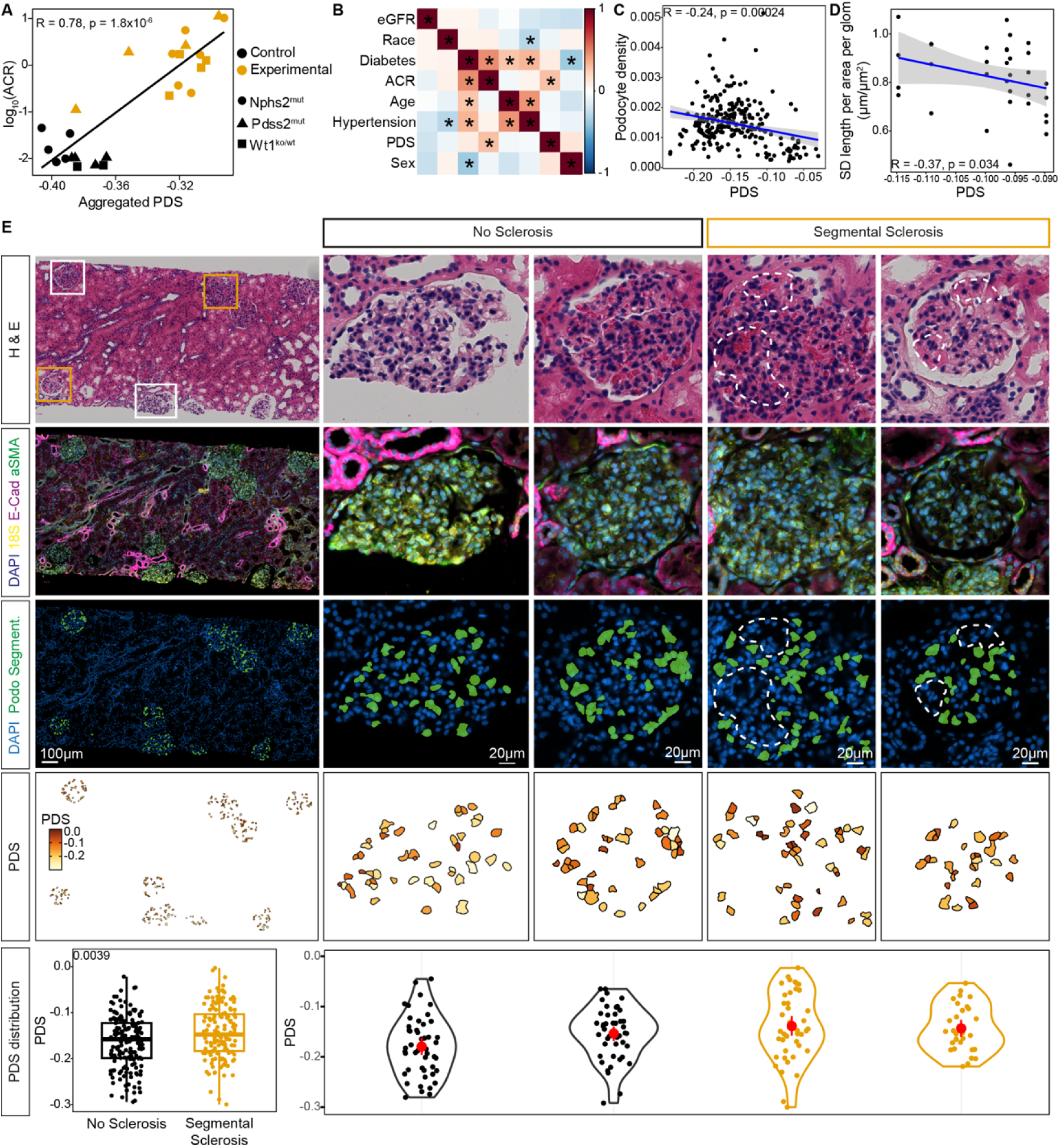
Functional validation of the PDS on mouse and human datasets A. Correlation between albuminuria (logarithm of the Albumin/Creatinine ratio in mg/mg (ACR)), an established clinical measure of podocyte damage, and the PDS for three different murine disease models (Nphs2^mut^, Pdss2^mut^, Wt1^ko/wt^). Cells of each snRNA-seq sample were aggregated (‘pseudo-bulk’) and represented as one symbol. Shape and color of symbols represent the study and experimental status of the sample, respectively. Spearman correlation and p-value are provided. B. Spearman rank correlation (color-scale) between the PDS and clinical attributes of acute kidney injury (AKI) and chronic kidney disease (CKD) patients as provided in the KPMP kidney tissue atlas (https://atlas.kpmp.org/). Asterisks denote correlations with p-value < 0.05. scRNA-seq (n = 17) and snRNA-seq (n = 37) data from patients with available albuminuria measurements were used. (eGFR: estimated glomerular filtration rate; ACR: urinary albumin/creatinine ratio). C. Significant negative correlation between PDS and podocyte density - calculated as number of podocytes per glomerular area - in human Xenium spatial transcriptomic data. PDS is calculated per individual podocyte and aggregated per glomerulus. Spearman correlation coefficient and p-value is shown in the top of the panel. Blue line indicates linear regression; gray area shows 0.95 confidence interval around the regression line. 238 gloms across 8 patients (each in 2 replicates) were analyzed. D. Significant negative correlation between slit diaphragm (SD) length as measured by confocal microscopy and PDS measured on identical glomeruli in human Xenium spatial transcriptomics (n = 7). Blue line provides regression; gray area indicates 0.95 confidence interval around the regression line. Spearman correlation coefficient and p-value are provided. E. Human Xenium spatial transcriptomic data from a 71 y/o female suffering from FSGS. Top row: H&E histology including overview and high magnification images of select glomeruli (see boxes in overview). Dashed white lines in high-magnification images indicate areas of potential segmental sclerosis. Second row: immunofluorescence staining signal used for cell segmentation including nuclei (DAPI, blue), 18S-RNA (yellow), E-Cadherin/CD45/ATP1A1 (all magenta), and Vimentin/alpha-smooth muscle actin (SMA, all green). Third row: location of segmented podocytes together with nuclei. Dashed lines indicate glomerular areas void of podocytes indicative of segmental sclerosis. Images in second and third row were generated with Xenium Explorer 3. Fourth row: PDS values (color scale) per single segmented podocyte. Podocytes with insufficient transcriptomic signal depth to calculate PDS were omitted. Bottom row: Boxplot shows PDS of individual podocytes, grouped by glomerular morphology (no sclerosis vs segmental sclerosis), p-value of the Wilcoxon test is shown in the top left. Violin plots show distribution of single podocyte PDS values (dots) within the glomerulus shown in high-magnification images. Red dot: median, red line: interquartile range.

As the validation of both, PDS and HDS showed remarkably robust results on the transcriptomic level, we further tested whether the damage reporting capacity of our markers could be expanded to the protein level. To this end, we calculated PDS and HDS on proteomic data from murine models of podocyte and liver disease, respectively.^13,46,50–52^ Without exception, PDS and HDS separated diseased from control samples (Figure S3F, Figure S4C). Future application of our scores may hence be extended to proteomics studies.

#### PDS and HDS are predictive in human data

PDS and HDS were developed using exclusively murine transcriptomic data and revealed very robust gene signatures. Hence, we asked next whether the signature was conserved in and therefore applicable to human data as well. To this end, where available, we identified human orthologues of all murine genes constituting the signatures (see Methods). For the PDS, we re-analyzed publicly available human data from the Kidney Precision Medicine Project (KPMP),^3^ including patient clinical characteristics, histology, sc/snRNA-seq data, and spatial transcriptomics. To test the clinical predictive value of the PDS in humans, we correlated human ACR with the PDS and other established clinical predictors of albuminuria such as age, sex, diabetic status across the KPMP cohort. Only diabetic status and the PDS were significantly correlated with ACR (Figure 2B, Figure S5A), thereby replicating the findings obtained in mice. Within the KPMP Visium spatial transcriptomic dataset, we identified a single biopsy core clearly showing an FSGS phenotype. Within this sample, PDS correlated with histologic features of FSGS distinguishing healthy from sclerosed glomeruli and even resolving segmental glomerulosclerosis within a single glomerulus (Figure S5B-S5C).

Motivated by the success on this existing human data, we generated new spatial human transcriptomics data using the Xenium Prime 5K platform, which provides substantially improved spatial resolution compared to the Visium system (spots 55 µm in diameter spaced at 100 µm vs < 50 nm precision for single transcript localization). Using a customized gene panel that included PDS marker genes, we generated such data for eight patients with minimal change disease or FSGS,^53^ and combined clinical data with histology, spatial transcriptomics, and confocal microscopy (to measure SD length per glomerulus). First, this analysis reproduced the correlation between podocyte density as a measure of hyperfiltration stress and PDS in humans (Figure 2C). Further, using SD length data and PDS obtained in identical single glomeruli, we confirmed a significant negative correlation between SD length and PDS (Figure 2D). Finally, leveraging the subcellular resolution and built-in image segmentation information in Xenium spatial transcriptomic data, we analyzed the distribution of single podocyte PDS values in glomeruli featuring segmental sclerosis. This analysis revealed that PDS values were elevated in sclerosed glomeruli compared to glomeruli without sclerosis (Figure 2E).

In addition, when we computed the PDS on scRNA-seq data of podocytes that were recovered from human urine within the NEPTUNE study,^54^ PDS correlated with the urine protein/creatinine ratios (UPCR) at the time of urine collection (Figure S5D). Finally, in microarray, bulk RNA-seq, and snRNA-seq data of human kidney biopsies, the PDS distinguished healthy control subjects from patients suffering from primary FSGS, IgA nephropathy, and chronic kidney disease, respectively (Figure S5E). ^3,7,55^ To conclude, a human ortholog-based PDS was readily capable of discriminating functional and morphological features of human podocytopathies.

In a parallel approach for the HDS, a total of 34 HDS marker genes had orthologs in humans. Using these orthologs, we calculated the HDS on hepatocytes of snRNA-seq human liver samples from two data sets including NASH and liver fibrosis phenotypes^56,57^. The HDS readily differentiated three healthy control subjects from three patients suffering from NASH (Figure 3A), as well as three healthy controls from three patients suffering from liver fibrosis (Figure 3B).^57^ Interestingly, the healthy controls represented a wide range of ages, which was reflected by higher HDS levels in older healthy patients (Figure 3C). To test if an increase of HDS was also associated with morphological tissue changes in humans, we analyzed spatial transcriptomic data (10X Visium) from liver sections of two patients with steatosis.^9^ Based on histology, spots in the spatial data were manually categorized as areas with or without steatosis (Figure S6A). Upon calculating the HDS, values in tissue-areas with steatosis were significantly higher than in tissue-areas without steatosis (Figure 3D-3E, Figure S6A). Taken together, our findings indicate that conserved cell-type specific damage scores can be developed from murine transcriptomic data that are subsequently amenable to translational application in humans.

**Figure 3:**
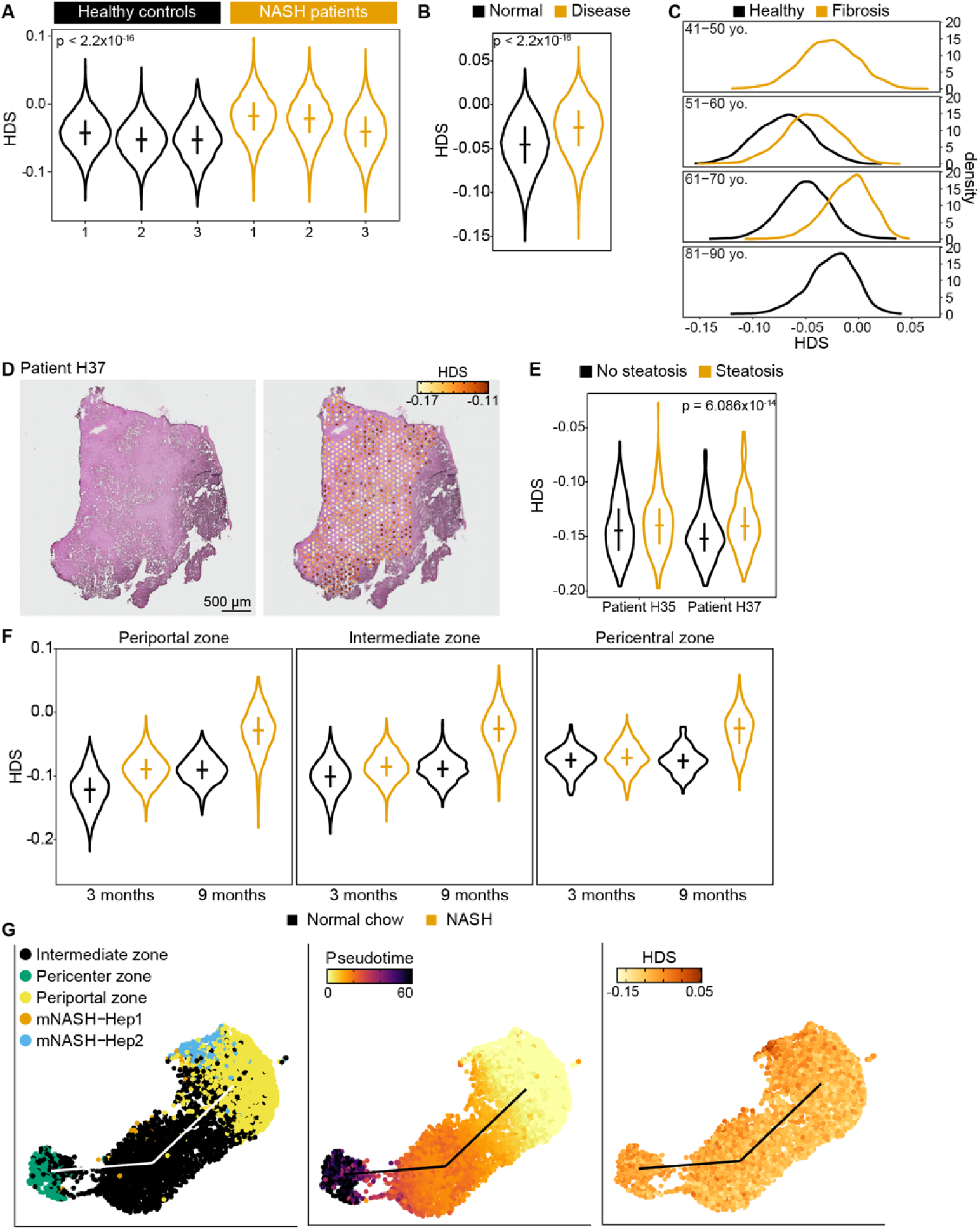
Functional validation of the HDS on mouse and human datasets A. Violin plots for HDS distribution in hepatocyte snRNA-seq data of human healthy subjects and NASH patients (GSE189600^56^). HDS marker genes were mapped to human ortholog genes. Age and sex of healthy donors in the same order as in figure: 47 y/o female, 50 y/o male and 50 y/o male. Age and sex of NASH patients in the same order as in figure: 49 y/o male, 64 y/o female and 31 y/o male. Horizontal lines show median of the distribution and vertical lines represent the IQR. P-value of one-sided Wilcoxon rank test (NASH versus healthy) is shown. B. Distribution of HDS in hepatocyte snRNA-seq data from three healthy subjects and three patients suffering from liver fibrosis (data from GSE210077^57^). Violin plots for HDS measurements pooled by health status (n = 3). HDS distribution is shifted to higher values in fibrotic individuals. One-sided Wilcoxon rank sum test statistic is shown. Horizontal lines show median of the distribution and vertical lines represent the IQR. C. Same data as in B. HDS increases with age in healthy human subjects and depending on disease status. Individuals are shown by increasing age range (see top of plots) from top to bottom. Each curve represents data from one individual colored by disease state (see legend). D. HDS reflects within sample variance of human liver steatosis. HDS calculation on 10x Visium human spatial transcriptomic data from the Liver Cell Atlas (GSE192742^9,48^). Only subjects with liver steatosis were analyzed. Left panel: histology, right panel: HDS per Visium spot overlay. Spots representing hepatocytes were manually annotated. Color scale represents the 10%- and 90%-quantiles of the HDS distribution across all samples. E. HDS separates steatotic from non-steatotic areas within the same human liver. 10x Visium spots of human steatotic livers as shown in D were manually sub-classified into steatotic and non-steatotic regions based on the H&E staining. Violin plots show HDS distribution in regions classified as non-steatotic (black) vs. steatotic (orange), including the sample from the patient shown in d and additional data shown in Figure S6A. Horizontal lines show median of the distribution and vertical lines represent the IQR. P-Value of one-sided Wilcoxon rank sum test on pooled HDS values for steatotic vs non-steatotic regions shown in top of plot. F. HDS in relationship to liver zonation in mice; hepatocyte snRNA-seq data taken from GSE189600^56^. Male mice were fed with standard or NASH promoting diet at 4 weeks of age for 3 or 9 months (n = 1). HDS distributions for cells from control (black) and NASH inducing diet (orange) are shown. Horizontal lines show median of the distribution and vertical lines represent the IQR. Statistical comparison not appropriate due to n=1 per group. Hepatocyte zone annotation provided by authors of data set (GSE189600^56^). G. HDS versus pseudo-time trajectories (PT), same data as in F; UMAPs show hepatocytes of a 3-month-old mouse fed NASH inducing diet. Each dot represents a hepatocyte colored by hepatocyte zone (left), PT (middle) and HDS (right). Hepatocyte zone annotation from original publication^56^. mNASH-Hep1/2: classification from original publication corresponding to damaged hepatocytes whose zone of origin could not be determined according to the authors. PT correlates with zonation, while the HDS is independent of liver zonation. See Figure S6B for additional data.

#### Comparison of HDS and trajectory inference

Pseudo-time trajectories have been proposed as an alternative approach for the computational alignment of single-cell and single-nucleus data along developmental, spatial and temporal gradients.^58^ Compared to the single-cell damage scores presented here, pseudo-time trajectories have potentially two disadvantages: first, they are not readily applicable across different studies, disease models and species as pseudo-time trajectories are commonly inferred per dataset or per study after appropriate data integration. Second, pseudo-time trajectories may have a disadvantage if gradients in single-cell data are dominated by signals other than cellular damage. An example is liver zonation: gene expression in hepatocytes is strongly determined by their relative position with respect to periportal and central veins, a signal that potentially ‘shades’ expression changes caused by cellular damage. To test this notion, we compared two established methods for pseudo-time trajectory inference (TSCAN^59^ and Monocle3^60^) to the HDS. Applying both algorithms to hepatocyte snRNA-seq data from mice suffering from NASH and controls, for which zonation information was available at individual hepatocyte level^56^, we confirmed that the HDS separated undamaged from damaged hepatocytes independent of their zone of origin (Figure 3F, Figure S6B), i.e. the HDS is independent of liver zonation. In contrast, the pseudo-time trajectories were dominated by gene expression patterns driven by hepatocyte zonation (Figure 3G, Figure S6C-S6D). Thus, the damage scores presented here provide information that is distinct from and complementary to pseudo-time trajectories.

### Characterizing cellular changes during disease progression

#### Pathway signatures reveal altered metabolism in damaged podocytes

Having validated our scores, we next sought to leverage their potential in addressing unanswered questions in kidney and liver biology. A key feature of the single-cell damage scores developed here is that they facilitate the analysis of progressive molecular changes associated with increasing cellular damage in disease models and even under unperturbed control conditions. To implement this notion, we developed a scheme that first sorts cells *in silico* along a damage trajectory from low to high damage and next identifies regulatory pathways and cellular processes (subsequently collectively called ‘pathways’) whose activity changes progressively along this axis. Cells with similar damage scores were binned (grouped) to reduce noise, and pathway activities per bin were determined using AUCell (see Methods) for all KEGG and Reactome pathways. We note that this approach does not compare gene expression variation between samples, which runs the risk of being affected by batch effects and other inter-individual variation. Instead, this scheme enables the analysis of disease progression within a single sample. Importantly, cells from control samples also displayed a spectrum of damage. Even though the average damage was lower in control samples compared to the experimental conditions, such samples always contained a small fraction of cells with relatively high damage scores (Figure 1B, 1E, 4A). This observation motivated us to sort cells by damage within the control samples and identify pathway activity changes along this gradient of otherwise unperturbed cells. Thereby, it became possible to identify pathway activity changes that were consistently changing in the same direction between experimental and control conditions, which can be considered general components of cellular damage and less likely attributable to a specific disease model or trigger. This approach allowed for addressing a key conundrum in the diagnostic approach to glomerular kidney disease, i.e., the contrast between a uniform histologic phenotype of podocyte damage – FSGS – and the multiple triggers, diseases and molecular pathways resulting in this phenotype. First, this poses a clinical challenge, as histology is often insufficient to provide a conclusive diagnosis. Second, it remained challenging to differentiate common from trigger-specific cellular events leading to the FSGS phenotype.

To address this challenge, we obtained published glomerular scRNA-seq data for four podocytopathy models,^61^ and generated snRNA-seq data for another three models (Figure S7) to reflect a wide range of podocyte damage triggers relevant to important aspects of podocyte biology as well as clinical settings. Together, these models include mutations in slit diaphragm proteins (Nphs2^mut^, Cd2ap^pko^), metabolic and oxidative stress (Pdss2^mut^, doxorubicin), antibody-mediated glomerular inflammation (NTN), diabetes (Btbr^ob/ob^), and mutation of a key podocyte transcription factor (Wt1^ko/wt^, Table S4).

To test if damage scores were quantitatively comparable across disease models and studies, we compared damage scores between control samples from different studies. If the scores are consistent, we would expect: (1) control samples across studies to show similar damage distributions, and (2) experimental samples to differ according to the degree of induced damage. This was indeed the case (Figure S8A). Pairwise Kolmogorov-Smirnov distances between PDS distributions were on average lower between controls than between experimental PDS distributions (two-sample Wilcoxon test, p = 2.5x10^-11^), arguing in favor of a common PDS baseline in control samples. A similar behavior was noted for the HDS (two-sample Wilcoxon text, p = 2,7x10^-4^, Figure S8B), supporting our notion that damage scores are robust and quantitatively comparable across studies. Interestingly, this approach allowed for identifying a small shift in average PDS baseline between scRNA-seq and snRNA-seq studies, suggesting that the additional damage exerted by tissue digestion for scRNA-seq was captured by the PDS (Figure S8A).

Next, we quantified pathway activity changes along the PDS axis as outlined above and focused on pathways that were either already known to be relevant to podocyte biology or changed highly dynamically with increasing damage. This way, we established ‘pathway fingerprints’ that characterize the regulatory, cell structural and metabolic changes happening during progressively increasing damage in a specific disease model (Figure 4A-4B, Figure S8C). These pathway fingerprints provide a rich source of information, allowing us to discern common and unique patterns of pathway changes across disease models. First, pathways that were expected to show model-specific behavior, due to the nature of various damage triggers, distinguished fingerprints from each other. For example, mutating the mitochondrial regulator Pdss2 was reflected by specific activity changes of the closely associated mitochondrial pathway ‘thermogenesis’. Similarly, in a complex with NPHS2 (podocin), tight junction proteins are part of the slit diaphragm protein complex,^62,63^ and the tight junction pathway within the fingerprint of the podocin mutation model (Nphs2^mut^) expectedly distinguished this model from others (Figure 4A). In addition to highlighting model-specific pathways, fingerprints also established pathways commonly modulated with podocyte injury and independent of the trigger.

**Figure 4:**
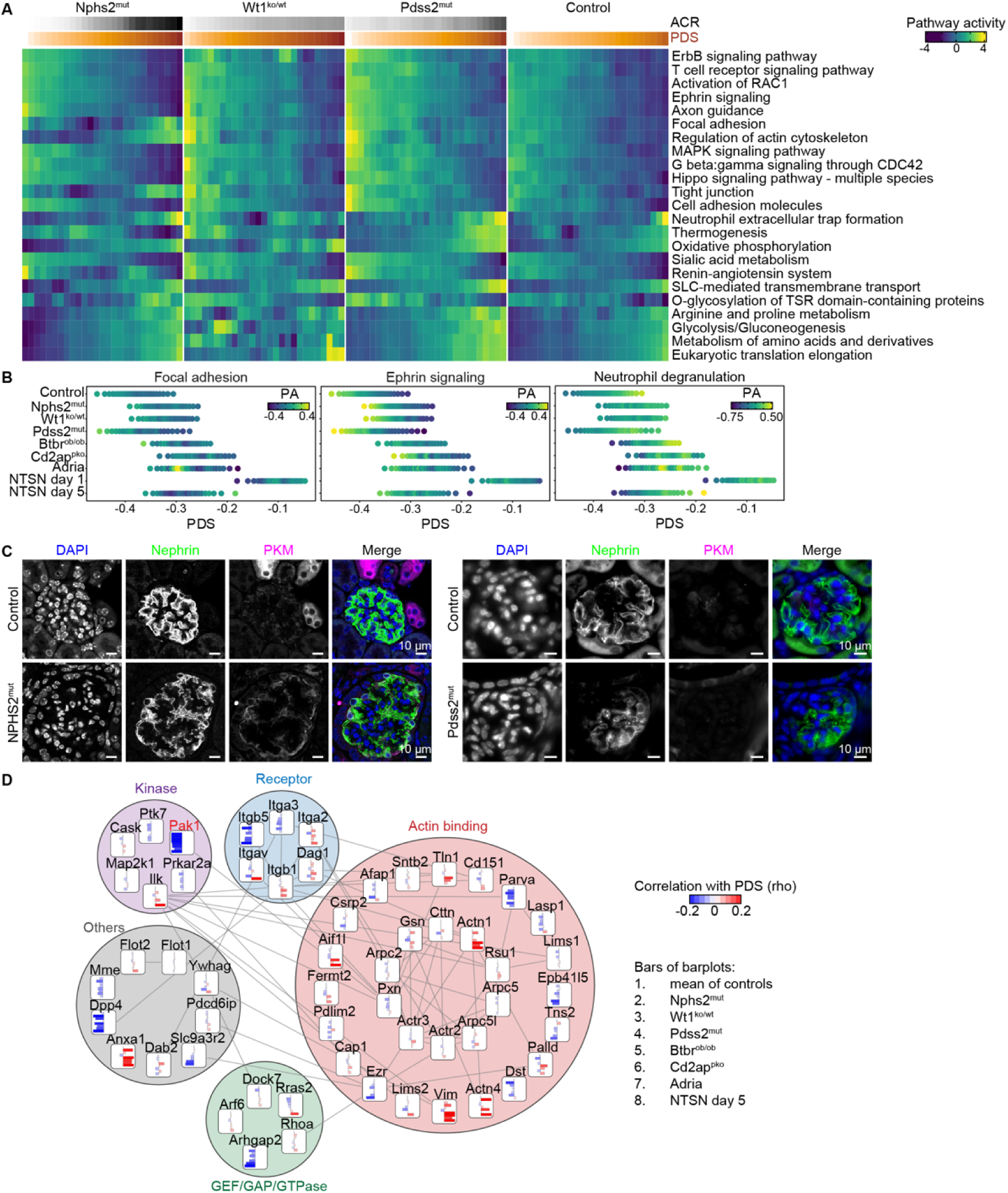
Application of PDS to FSGS datasets reveals model-specific pathway fingerprints. A. PDS pathway fingerprints for single-nucleus RNAseq data from Nphs2^mut^, Wt1^ko/wt^ and Pdss2^mut^ models of FSGS. ‘Control’ is the average of control samples from all three studies. Nuclei were sorted by increasing PDS (color bar on top). Pathway activities for KEGG and Reactome pathways were calculated using AUCell (see Methods) and smoothed using a sliding window of width twice bigger than step size, both adjusted to produce 27 bins in each study. Color shows pathway activities centered and scaled per pathway. Activity of pathways shown is significantly correlated with PDS in at least one of the three studies (Spearman rank correlation; adj. p-value < 0.05). Rows are clustered by pathway similarity. Color gradients on top show average albumin/creatinine ratio (ACR) and the average PDS per bin. B. Pathway activities for selected pathways calculated on sn/scRNA-seq data of cumulated control samples and various disease models. Cells/nuclei were grouped in evenly sized bins of 30 cells/nuclei with similar PDS (x-axis). Color indicates pathway activity (PA) centered and scaled per pathway. ‘Control’ corresponds to the average pathway activities across control cells of all models. Range of PDS per model reflects different damage levels between models and cumulated controls. C. Differential regulation of energy metabolism. Immunofluorescence staining of control and experimental kidney tissue from either Nphs2^mut^ 8-week-old (left) or Pdss2^mut^ 21-week-old (right) showing DAPI (blue), nephrin (green) and PKM (magenta). Mice of both models reflect similar damage levels as by ACR and SD length (see Figure 2A, Figure S3C). PKM expression is upregulated in podocytes in Nphs2^mut^ mice but not in Pdss2^mut^ mice. D. Integration of the *in vivo* podocyte adhesome protein-protein interaction network with PDS analysis. ^74^ Proteins are grouped by function, edges indicate protein-protein interaction. Bars inside each node show Spearman rank correlation (rho, see color scale) between PDS and the expression level of the gene per disease model (see legend on the right). Red and blue colors depict positive and negative correlations, respectively. Saturation of the color and length of the bar reflect strength of correlation. The marked upregulation of Actn1, Anxa1, and Vim exclusively in studies applying scRNA-seq with prior glomerular digestion is likely a technical artifact. Pak1 (marked in red) is a PDS marker gene.

For example, hippo signaling, which was initially found downstream of Wt1 transcriptional regulatory function in podocytes,^17^ was now shown to be unequivocally decreased upon podocyte damage in all models. Key pathways in podocyte biology related to the actin cytoskeleton, cell adhesions or the renin angiotensin system were also commonly modulated (Figure 4A). In addition to cell structural and regulatory changes, we also observed common metabolic changes of podocytes that were associated with increasing damage. Energy metabolism (glycolysis and oxidative phosphorylation) and protein production (translation) were consistently upregulated with increasing damage across models, suggesting that podocytes are majorly reorganizing their proteome while changing their shape and coping with increasing stress. In the clinical context, albuminuria is used as a measure to ‘align’ the disease status between patients. Here, we test if the PDS could serve a similar purpose for ‘aligning’ cellular damage states across disease models. Therefore, we integrated damage levels as indicated by the PDS with pathway activity changes across disease models (Figure 4B). This approach allows for aligning and directly comparing multiple disease models, samples or patient biopsies with respect to damage levels and associated signaling pathway behavior. Here, ephrin signaling, which was recently identified as a pathway relevant to the progression of kidney disease,^64^ was consistently downregulated upon podocyte damage. To experimentally test the relevance of ephrin signaling in the context of podocyte damage, we performed fluorescent labelling and STED imaging of Ephrin type-B receptor 1 (Ephb1), which is one of the PDS marker genes (Figure S3B). This analysis confirmed the localization of Ephb1 to the slit diaphragm and showed substantial depletion in damaged podocytes.

To confirm our approach and validate the alignment of independent datasets based on PDS experimentally, we confirmed the overactivation of the tight junction pathway in Nphs2^mut^ compared to Wt1^ko/wt^ mice. We carried out immunofluorescence staining followed by super-resolution microscopy for the SD marker nephrin and the tight junction protein claudin-5 in both disease models (Figure S9A). Claudin-5 has been shown to aggregate in the slit diaphragm upon podocyte damage,^65^ and plays a protective role in diabetic nephropathy.^24^ Our approach allowed for the parallel quantification of both, podocyte damage based on SD length and tight junction activation based on the degree of claudin-5 aggregation. Hence, we were able to obtain quantitative measures of pathway activation and podocyte damage independent of the PDS. By correlating SD length with claudin-5 aggregation, we indeed confirmed that in Nphs2^mut^ mice, tight junction activation was more pronounced at similar damage levels when compared to Wt1^ko/wt^ mice, as was evidenced by a stronger negative correlation in Nphs2^mut^ mice (Figure S9B).

In a second experimental confirmation of our pathway activity analysis, we addressed the podocyte energy metabolism represented by the pathways ‘thermogenesis’, ‘oxidative phosphorylation’ and ‘glycolysis / gluconeogenesis’ in our fingerprinting analysis (Figure 4A). While the activity of these pathways was generally upregulated with increasing podocyte damage in all models, we noticed that at high damage levels, a pronounced over-activation was evident in Pdss2^mut^ mice when compared to Nphs2^mut^ mice (Figure 4A-4B). For both models, consistent SD length and albuminuria measurements were available (Figure 2A, Figure S3C-S3D), allowing for gauging and aligning damage levels independent of the PDS. To evaluate key regulators of energy metabolism in podocytes, we stained kidneys for phosphorylated S6-kinase, which acts downstream of mTORC1 in the mTOR pathway, a major regulator of podocyte energy metabolism contributing to FSGS pathogenesis.^66,67^ Supporting our fingerprinting analysis, we saw differential activation of S6-kinase in podocytes between Nphs2^mut^ and Pdss2^mut^ mice at similar damage levels (Figure S9C). Podocytes heavily rely on anaerobic glycolysis as energy source.^66^ A key rate limiting enzyme controlling the balance between glycolysis, (anaerobic) lactate production and (aerobic) oxidative phosphorylation is pyruvate kinase (Pkm), which in turn is induced in podocytes by S6-kinase,^68^ and is a protective factor in diabetic nephropathy.^69,70^ Pkm function is controlled by two splice isoforms of the Pkm gene: the Pkm1 isoform dominates in tissues with high demand of oxidative phosphorylation (muscle, brain), while the Pkm2 isoform balances metabolism between anaerobic and aerobic pathways in most tissues, including podocytes (Figure S9D).^69,70^ First, we confirmed that Pkm2 is by far the predominant isoform in podocytes (Figure S9E), and showed that Pkm expression was significantly higher in Nphs2^mut^-derived compared to Pdss2^mut^–derived podocytes (Figure S9F). Immunofluorescent staining for Pkm in Nphs2^mut^ and Pdss2^mut^ glomeruli further confirmed the specific upregulation of this key energy metabolism regulatory factor in Nphs2^mut^ podocytes on the protein level (Figure 4C). In summary, corresponding to our pathway fingerprinting analysis and assaying two regulators of energy metabolism, we verified the differential activation of energy metabolism pathways as they depend on damage triggers and damage levels. This supports the applicability of the PDS in investigating differential molecular changes across podocyte damage models, aligned at similar damage levels.

In summary, pathway fingerprints and damage trajectories identified trigger-specific as well as commonly regulated pathways upon podocyte damage, rendering the PDS suitable as molecular diagnostic approach for FSGS to resolve its molecular etiology.

#### Identifying generic druggable targets with the PDS

A curious result of our pathway fingerprint analysis was the aberrant and model-specific behavior of the pathways “Focal adhesion” and “Regulation of actin cytoskeleton” in the Nphs2^mut^ model compared to all other models (Figure 4A). Focal adhesions attach podocytes to the glomerular basement membrane (GBM), making them a crucial organelle to the GBM detachment of podocytes during FSGS.^62,63^ Focal adhesions and the slit diaphragm are further tightly connected to the actin cytoskeleton, which is commonly modulated and rearranged upon podocyte damage.^71^ A key morphological feature associated with this rearrangement is foot process effacement. We therefore utilized the PDS to gain deeper insight into the reorganization of the ‘podocyte adhesome’ upon damage. We correlated the expression of members of the podocyte adhesome with the PDS in each of the disease models and control samples (Figure 4D).^72^ Selected members of this network were distinctly regulated in the Nphs2^mut^ model compared to all other models. For example, two genes encoding for actin binding proteins, dystonin (Dst) and palladin (Palld), exhibited strong negative correlations with the PDS in the Nphs2^mut^ model, whereas in other disease models they were uncorrelated or even oppositely correlated with the PDS (Figure 4D). These examples underline the importance of distinguishing model-specific changes that potentially act upstream of podocyte damage from alterations that are consistent across models. Therefore, we next identified adhesome components that were consistently downregulated with increasing damage irrespective of the damage trigger. These components included the actin binding molecules Parva and Epb41I5, the extracellular matrix receptors Itgb5 and Itga3, the cell surface peptidase Dpp4, and the kinase Pak1 (RAC1 activated kinase 1). All of these components had previously been identified as relevant to podocyte focal adhesion function.^72–75^ Parva and Epb41l5 in particular have been shown to prevent podocyte detachment from the GBM in FSGS.^73,74^ Pak1 is a key regulator of the focal adhesion and actin cytoskeleton networks and has been shown to be involved in the earliest events of FSGS in a CTCF deletion model.^76^ Further, our work surprisingly reveals that downregulation of the cell-surface peptidase Dpp4 (dipeptidyl peptidase-4) is a hallmark of increasing podocyte damage across disease models, including diabetes. Dpp4 has been reported as upregulated in diabetes in several tissues and cell types and contributes to decreased insulin sensitivity. In podocytes, Dpp4 physically interacts with the focal adhesion protein integrin beta-1 (Itgb1),^77^ and Dpp4 inhibitors decrease albuminuria in mouse models of podocyte damage.^78^ These drugs are a mainstay in the treatment of type 2 diabetes mellitus, where their use was associated with beneficial albuminuria outcomes in several clinical trials.^79–82^ Taken together, the surprising downregulation of Dpp4 in seven independent disease models can be interpreted as a repair response amenable to preventative treatment. Thus, our integrative adhesome and PDS network analysis revealed promising targets for the prevention of podocyte GBM detachment in podocytopathies.

### Transcriptional networks driving podocyte damage

#### Regulators of podocyte gene expression drive cross-model correlates of the PDS

To better understand the regulatory programs driving the expression changes of genes relevant to the podocyte damage response, we constructed a podocyte-specific transcription regulatory network (TRN) (Figure 5A). To derive this TRN, we acquired podocyte-specific ATAC-seq data by FACS-sorting podocyte nuclei from a nuclear GFP reporter mouse (Methods, Figure S10A). Next, we performed foot-printing and transcription factor (TF) motif scanning within accessible chromatin regions, using DNA binding motifs of 110 TFs that were expressed in podocytes. Finally, we grouped TFs with very similar DNA binding motifs into 14 TF families, as their binding signatures were difficult to distinguish without further information (Figure S10B). We combined predicted target genes of all motifs within each cluster, forming a TF regulon, and named each regulon after a representative transcription factor (Figure S10C). This established a network between 14 TF families and 6,360 target genes expressed in podocytes that will serve as a critical resource also for future research into podocyte biology.

**Figure 5:**
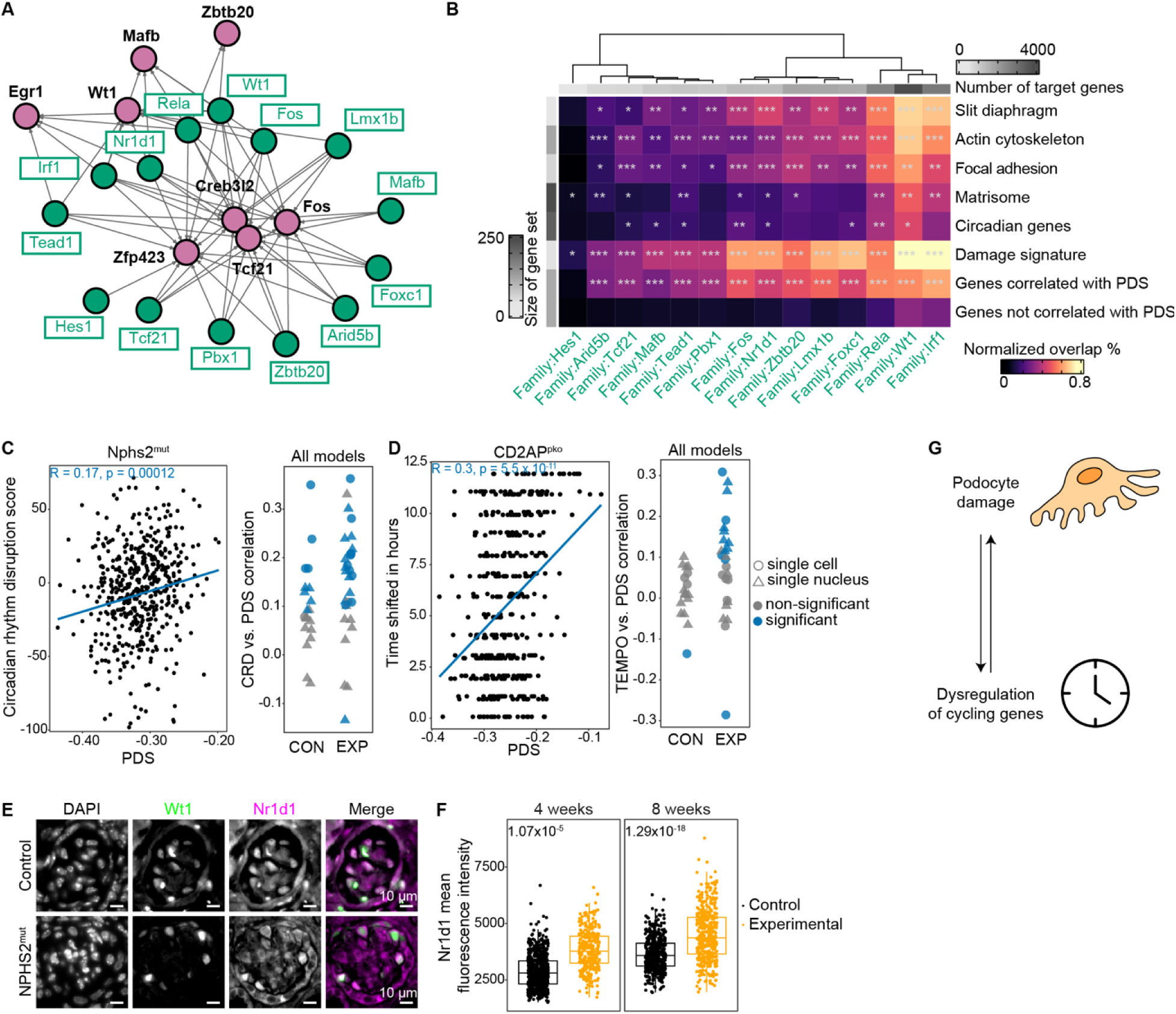
PDS reveals circadian dysregulation as a central mechanism in FSGS A. Podocyte transcriptional regulatory network (TRN), limited to transcription factor (TF) genes correlating with PDS in multiple studies (magenta nodes) and TF family regulons (green nodes). Edges indicate regulation of TF gene by respective TF family. B. TRN analysis reveals the circadian core TF Nr1d1 (Rev-Erb alpha) as a key regulator of podocyte genes. Heatmap of overlap (color scale) of TF target genes with gene sets as shown on the right. Color intensity shows the fraction of genes in gene sets (rows) regulated by the respective TF families (columns). Significance of the overlap was calculated with Fisher’s exact test, FDR-adjusted, and depicted on the heatmap with asterisks: *** q < 0.001, ** q < 0.01, * q < 0.1. Row and column annotation bars (gray scales) reflect size of gene sets or target genes, respectively. Bottom row: TF target enrichment for 100 randomly selected genes that are expressed in podocytes, but do not correlate with the PDS as a negative control. C. Increased podocyte damage correlates with increased circadian disruption. Left: Relationship of circadian rhythm disruption (CRD) score^91^ and PDS. Each dot represents one nucleus from snRNA-seq data of podocytes isolated from one individual Nphs2^mut^ mouse. Regression line and parameters for Spearman rank correlation are shown. Right: Spearman rank correlation coefficients (calculation as exemplified for one individual mouse on the left) for all individual control (CON) and experimental (EXP) samples of seven different mouse models of podocyte damage (Nphs2^mut^, Pdss2^mut^, Wt1^ko/wt^, Btbr^ob/ob^, NTN day 5, Cd2ap^pko^, Adria). Color distinguishes if the CRD to PDS correlation is significant (p < 0.05). See legend in panel D for meaning of symbols. D. Left: Relationship of absolute deviation of estimated circadian time from the reference sample time and the PDS. Circadian time of each cell was estimated by the TEMPO algorithm, using *Clock* as the reference gene. Each dot represents one cell from scRNA-seq data of podocytes isolated from one individual Cd2ap^pko^ mouse. Regression line and parameters for Spearman rank correlation are shown. Right: Spearman rank correlation coefficients (calculation as exemplified for one individual mouse on the left) for all individual control (CON) and experimental (EXP) samples of seven different mouse models of podocyte damage (Nphs2^mut^, Pdss2^mut^, Wt1^ko/wt^, Btbr^ob/ob^, NTN day 5, Cd2ap^pko^, Adria). Color distinguishes if the TEMPO to PDS correlation is significant (p < 0.05). E. Representative immunofluorescence staining of kidney tissue from 8-week-old Nphs2^mut^ control and experimental mice for DAPI, Wt1 (green) and Nr1d1 (magenta). Nr1d1 nuclear localization is disrupted in FSGS glomeruli. F. Quantification of Nr1d1 mean fluorescence intensity in podocyte nuclei for control and experimental Nphs2^mut^ mice aged 4 weeks (n = 20 technical replicates from 4 biological control animals, n = 20 technical replicates from 2 biological experimental animals), and 8 weeks (n = 20 technical replicates from 3 biological control animals, n = 20 technical replicates from 3 biological experimental animals). Podocyte nuclei were identified by segmenting first by DAPI-then by Wt1-positive fluorescent signals prior to the quantification of Nr1d1 fluorescence intensity. Significant larger variations in the Nr1d1 nuclear signals were observed in experimental mice for both 4-week-old and 8-week-old mice. P-values for F-test are shown. G. Schematic of regulation between damage and dysregulation of circadian genes in podocytes.

To systematically identify target genes of these regulons that were involved in FSGS progression across models, we computed how often a gene was correlated with the PDS across snRNA-seq studies of FSGS, in both experimental and control conditions. Note that this detection of commonly perturbed genes in experimental and control conditions cannot be achieved with traditional differential expression analysis but necessarily requires a within-sample trajectory analysis.^83^ Based on these results we selected 152 genes that were significantly correlated (Table S5, t-test, adjusted p-value < 0.1) with the PDS in seven or more tests. Forty of those genes were already members of the PDS marker gene set.

This set of 152 genes contained eight podocyte-expressed TFs, i.e. out of the 110 TFs that were used to create the TRN. Thus, we used the TRN to inspect the regulatory relationships among these eight TFs that were tightly associated with podocyte damage (Figure 5A). These TF genes included the core podocyte TFs Wt1, Mafb, and Tcf21, TFs known to respond to external damage cues (Egr1, Fos), and TFs with yet uncharacterized functions in podocytes (Zbtb20, Zfp423, Creb3l2). The regulators established two groups: one group of regulators controlling all eight PDS-associated TFs and another set of regulators only controlling four of them (Egr1, Wt1, Mafb, Zbtb20). Confirming previous studies, we identified the regulons for the Wt1-, Lmx1b-, and Tead1-families as controlling the majority of TF genes correlating with the PDS.^17^ Interestingly, in addition to these established podocyte regulons, the Irf1, Nr1d1, and Rela families were similarly prominent, indicating that inflammatory signals and processes dependent on nuclear receptor signaling such as the circadian clock may be relevant to podocyte damage (Figure 5A).^83^

#### Podocyte damage disrupts the circadian clock

Next, we utilized the podocyte TRN to identify potential regulators of genes that were likely involved in podocyte damage either because their own expression correlated with the PDS (i.e. the 152 genes from above) or because they are involved in cellular functions that are known to be relevant for podocytopathies, like the podocyte adhesome,^72^ matrisome,^84^ actin cytoskeleton,^85^ and slit diaphragm.^63^ Consistent with earlier work we found that Wt1 was predicted to bind regulatory elements of the majority of genes involved in these key podocyte structures as well as of genes correlating with the PDS (Figure 5B). Next to Wt1, the TF families containing Rela and Irf1 (both of which are reacting to inflammatory stress) were prominent regulators of these damage-associated gene sets. Further, all gene sets were co-regulated by a combination of cell type-specific podocyte TFs (e.g. Wt1, Lmx1b, Foxc1, Mafb, Tcf21) and generic TFs (e.g. Tead1, AP-1 complex / Fos, Figure S10D). The fact that some of the generic regulators were themselves correlated with the PDS (e.g. Fos) is suggesting that the transcriptional response to podocyte damage is coordinated via an intricate network of generic and cell type-specific TFs (Figure S10C. A striking finding was that nuclear receptor subfamily 1 group D (Nr1d1, Rev-Erb alpha) was predicted to regulate a wide range of genes that we found to be correlated with the PDS (Figure 5B). Rev-Erb alpha is one of the core TFs regulating feedback loops within the intracellular circadian clock,^86^ and is instrumental in establishing circadian gene expression. Recent studies have established that many key podocyte genes are expressed in a circadian fashion and that the disruption of the podocyte circadian clock results in podocyte dysfunction.^87–89^ Based on the tight link between Nr1d1 and many of the PDS-associated processes across disease models, we speculated that the reverse may also be true, i.e. that podocyte damage interferes with the circadian clock. To experimentally validate this link, we first generated circadian gene expression data from whole murine kidneys by performing transcriptomic profiling in 3-hour time intervals over a 48-hour period. Using this data, we determined genes with circadian expression oscillation and intersected this gene list with podocyte expression data to obtain a set of genes with circadian expression in podocytes (Methods, Figure S11A-S11B). This gene set was indeed significantly enriched in podocyte Nr1d1 target genes as determined in our TRN (Figure 5B).

We then asked if increasing podocyte damage as measured by the PDS was associated with increased disruption of circadian gene expression. To this end, we employed two different methods to determine disruption of circadian gene expression in single cells – TEMPO,^90^ which determines average time shifts between the expression of core clock genes and other circadian genes, and CRD,^91^ which measures the variance among circadian genes. Here, we fully exploited the single-cell capacity of our score as damage-dependent circadian expression heterogeneity was captured within one sample, with the organism being normalized for all common factors potentially affecting circadian rhythms such as cortisol levels, organism redox state, time of day, blood pressure etc. Both approaches revealed that increased podocyte damage correlated with increased disruption of circadian gene expression within sample, an effect that was shown across podocytopathy models (Figure 5C-5D). We further confirmed this result by repeating the analysis using a recently published, podocyte-specific set of circadian genes, yielding an even more pronounced difference between controls and experimental samples (Figure S11C).^87^

We hypothesized that upon desynchronization of circadian gene expression, even within single glomeruli, the core circadian clock TF Nr1d1 (Rev-Erb alpha) would show an increased variance of nuclear protein levels and a shift in nuclear vs. cytoplasmic localization, respectively. Indeed, IF co-staining for Nr1d1 and Wt1 revealed an increased variance of nuclear to cytoplasmic localization of Nr1d1 in podocytes in Nphs2^mut^ mice compared to controls (Figure 5E). Densitometry for Nr1d1 further showed that the variance of nuclear Nr1d1 levels increased with age and hence with progressive podocyte damage (Figure 5F). Of note, the Nphs2^mut^ model does not interfere with circadian clock components directly, supporting an effect of podocyte damage on the circadian clock. Similar to our PDS-based transcriptomic approach, this immunohistochemical approach was normalized and controlled for external effects on zeitgeber time, as nuclei within single glomeruli from the same animal were assessed.

In conclusion, disruption of circadian gene expression appears to be a common consequence of podocyte damage in response to multiple damage triggers. Given the large number of essential podocyte transcripts exhibiting circadian oscillation,^87^ this finding bears significant relevance to understanding podocyte damage. Further, given that disruption of the clock and hence circadian gene expression results in podocyte damage,^88^ and podocyte damage results in disruption of circadian gene expression, positive feedback loops contributing to podocyte damage by means of this mechanism appear likely (Figure 5G). In summary, our integrative analysis of the PDS together with the TRN have yielded important insights into mechanisms of podocyte damage.

### HDS reveals metabolically distinct cell state trajectories of damaged hepatocytes

Accumulation of damage can drive hepatocytes into highly distinct cellular states, including neoplastic transformation (i.e. establishing hepatocellular carcinoma, HCC), cell death, and senescence^92,93^. The accumulation of hepatocyte senescence in NAFLD has been shown to result in worse outcomes.^94^ However, since the pathophysiological role of senescent hepatocytes in chronic liver disease has not been widely investigated nor fully understood, we focused specifically on the transition of damaged hepatocytes into this cell state.^92^ To explore this cell state transition, we used hepatocyte snRNA-seq data from mice either fed a cholin-deficient and ethionine-supplemented (CDE) diet, or treated with thioacetamide (TAA), as well as controls. While both liver disease models eventually evoke hepatocellular carcinoma, CDE elicits severe steatosis with little fibrosis at early stages, whereas TAA features only mild steatosis but severe inflammation, necrosis and fibrosis.^95^ Samples used for snRNA-seq were from mice sacrificed after 3 weeks of treatment, in a precancerous state.^96^ Upon HDS analysis, greater average hepatocyte damage was seen in TAA hepatocytes compared to CDE or controls (Figure 6A).^56^ To assess the senescence status of hepatocytes, we quantified the activity of the Senescent Hepatocyte Gene Set (SHGS) per single cell.^94^ First, we evaluated the senescence status of cells in relation to the HDS and observed an apparent disease model-specific increase of senescent cells predominant in TAA-exposed hepatocytes (Figure 6A, Figure S12A). Interestingly, with increasing cellular damage this analysis revealed a ‘tipping point’: until an HDS of about -0.05 we did not observe any noticeable increase in the activity of the senescence gene signature (Figure 6B). Beyond this threshold value, cells increasingly adopted a senescent-like state (Figure 6B). This tipping point pattern was absent in cells from control animals, as hardly any cells reached the HDS threshold value (Figure S12B). However, the pattern was visible in both disease models (TAA and CDE). These findings indicated that the tipping point mostly depends on damage accumulation in individual hepatocytes (Figure S12B) but is independent of the specific disease model and the average damage level. Further, cells started to adopt divergent cell states with increasing HDS. Although a large fraction of cells became increasingly more senescent-like, many other cells were able to maintain a low senescence score despite being substantially damaged according to their HDS. To formally quantify this notion, we computed the inter quartile range (IQR) of the range of senescence score as a function of the HDS, which revealed a substantial increase of the IQR with increasing damage (Figure 6B). Thus, these findings support the notion that damaged hepatocytes develop along divergent trajectories towards distinct cellular states, one of them being cellular senescence. While cellular damage beyond a certain threshold seems to be a necessary requirement for the entry into the senescent state, that cell state transition does not seem to be fully deterministic, as a large fraction of highly damaged hepatocytes escapes entry into senescence. We confirmed these results in another independent NASH snRNA-seq murine dataset (Figure S12C-S12E).^56^ Furthermore, this tipping point was also preserved in human liver snRNA-seq data from patients suffering from NASH (Figure S12F-S12G). Of note, the value of the HDS tipping point remained consistently in the range of -0.05 between all three datasets (i.e. even across species), corroborating our previous analysis on the ability of the HDS to align independent studies (Figure S8B).

**Figure 6:**
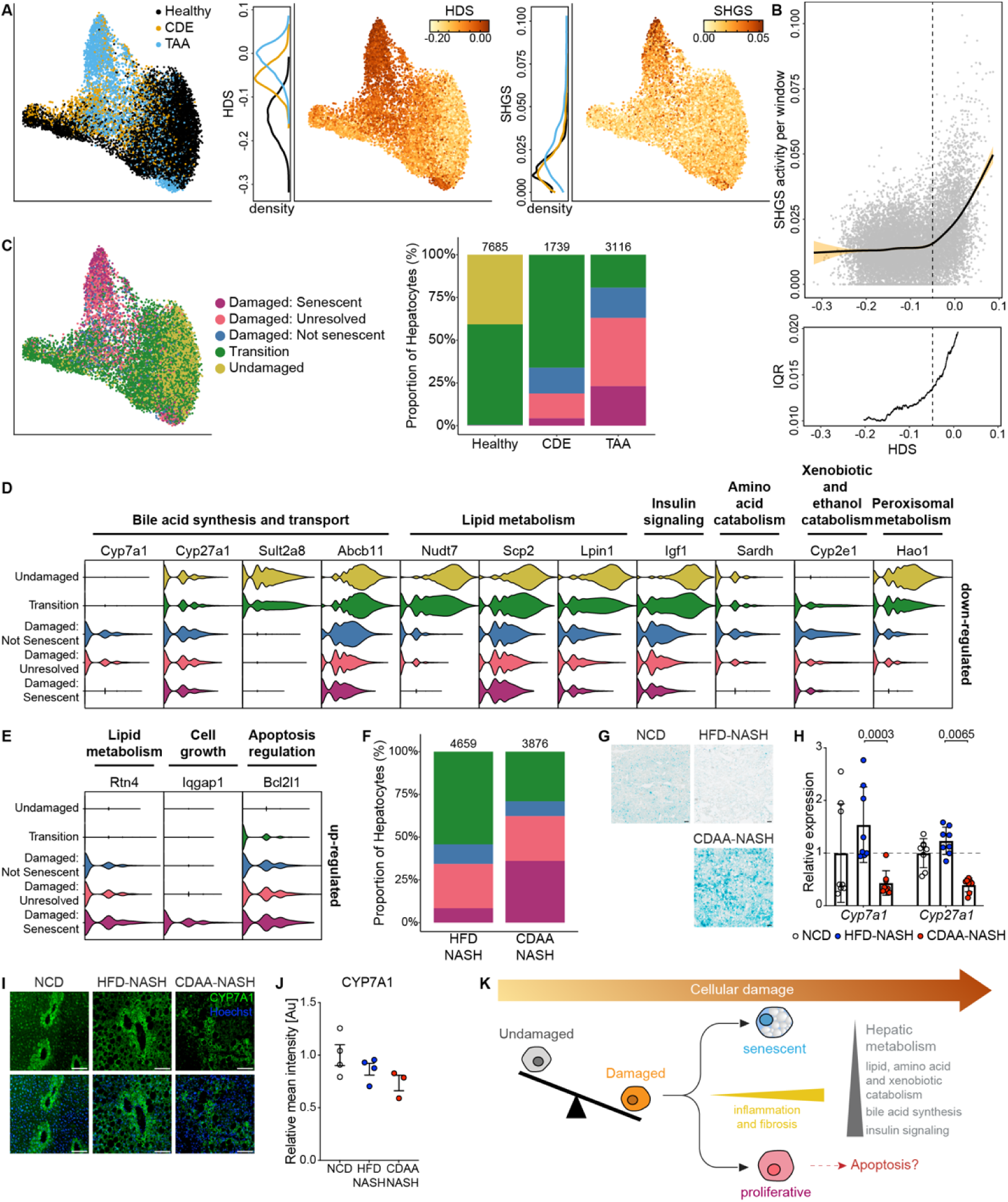
HDS reveals damage-driven cell state transition of hepatocytes. A. UMAP of merged snRNA-seq data from mice subjected to three different treatments. Healthy: normal chow; CDE: choline-deficient, ethionine-supplemented (CDE) diet; TAA: thioacetamide (TAA) in the drinking water (GSE200366^96^, n = 3 biological replicates per treatment). Left: UMAP of hepatocytes colored by condition. Middle: hepatocytes colored by HDS (color scale: floor and ceiling values corresponding to 1%- and 99%-quantiles of the HDS distribution across all samples). Right: Hepatocytes colored by Senescent Hepatocyte Gene Signature (SHGS). Density plots next to UMAP show HDS and SHGS distribution per condition, respectively (see left UMAP plot for legend). HDS is significantly higher in both CDE and TAA compared to controls (one-sided Wilcoxon rank sum test, for both comparisons p-value < 2.2 x 10^-16^). B. SHGS activity score versus HDS (same data as in A). Top: Solid line: Generalized linear model line shows behavior of SHGS activity with increasing HDS. Orange: confidence interval of regression line. The data is heteroscedastic (Breusch-Pagan test, p < 2.2 x 10^-16^). Bottom: IQR per sliding window of 2000 hepatocytes (y-axis) plotted against mean HDS per sliding window. SHGS increase shows a tipping point pattern beyond an HDS damage threshold of ∼ -0.05. C. Left: UMAP as shown in a, colored by cell state classification. Right: relative abundance of hepatocytes of each cell state per condition. Numbers above stacked bars show the total number of hepatocytes analyzed per condition (n = 3 per model). D. Gene expression of select genes from a group of 475 genes significantly (Bonferroni adjusted p-value < 0.05 and LFC < -0.5, Table S8) down-regulated in damaged, senescent hepatocytes compared to damaged, not-senescent hepatocytes. Bile acid metabolism is specifically downregulated in damaged, senescent cells. E. Gene expression of selected genes of a total of 422 significantly (Bonferroni adjusted p-value < 0.05 and LFC > 0.5, Table S7) differentially up-regulated genes in damaged, senescent hepatocytes compared to damaged, not-senescent hepatocytes. F. Damaged hepatocyte cell state assignment similar to panel C, based on scRNA-seq data from two murine NASH models (n = 1, biological replicates per model): choline-deficient, L-amino acid-defined (CDAA) high-fat diet, and high-fat, high-fructose diet supplemented with trans-fat and cholesterol (HFD) (GSE146049^109^). Cell state classification was carried out based on HDS and SHGS as in panel C. Numbers above stacked bars show the total number of hepatocytes. Color legend is the same as in C. G. Senescence-associated β-galactosidase staining of liver tissue derived from 26-week-old mice fed a CD, HFD-NASH, or CDAA-NASH diet (same experimental conditions as in F). Representative images of n = 3 mice per condition are shown. Scale bars = 100 µm. H. RT-qPCR for the bile acid metabolism enzymes *Cyp7a1* and *Cyp27a1* expression in liver tissues of 26-week-old mice fed a CD, HFD-NASH, or CDAA-NASH diet. Gene expression is shown relative to CD (two-way ANOVA with Tukey’s multiple comparisons test, n = 8). Whiskers represent -/+ SD. I. Representative immunofluorescence staining of liver tissue from NCD, HFD-NASH, and CDAA-NASH animals stained for CYP7A1 (green) and counterstained with Hoechst (blue). Livers were collected from 12-16-week-old NCD-fed mice or 26-week-old NASH-diet-fed mice (n = 3-4 biological replicates per condition). Scale bars = 100 µm. J. Quantification of CYP7A1 mean intensity across IF staining of NASH models as shown in I. Data are represented as mean of CYP7A1 intensity per mouse and relative to NCD (n = 3-4 mice per condition). Whiskers represent -/+ SEM. K. Schematic model of hepatocyte cell state transitions. Accumulating damage leads to loss of cellular homeostasis, increasing profibrotic and proinflammatory signals. After a critical HDS is reached, hepatocytes begin transitioning into different pathological cell state trajectories. When transiting in the direction of senescence, hepatocytes strongly downregulate genes involved in defining hepatic metabolic functions such as amino acid and xenobiotic catabolism, insulin signaling, and bile acid synthesis. In parallel genes involved in lipid catabolism are downregulated, thereby further exacerbating lipid accumulation in senescent cells.

To characterize the transcriptional alterations associated with these cell state transitions, we arbitrarily divided cells on the continuous HDS and SHGS trajectories in three classes per score, respectively: For the HDS, cells were grouped into undamaged hepatocytes (low HDS), damaged hepatocytes (high HDS), and cells in a ‘transition’ state with intermediate HDS. Damaged hepatocytes were further stratified on the SHGS scale as ‘senescent’, ‘non-senescent’ or ‘unresolved’ depending on their senescence score (see Methods). This grouping allowed us to quantify expression differences between the extremes on both scales. The relative abundances of hepatocytes in each cell state category varied greatly between the CDE and TAA disease models (Figure 6C). Confirming our UMAP and density plots (Figure 6A), we observed a sharp increase of the fraction of damaged hepatocytes in TAA mice when compared to CDE mice: whereas ∼30 % of CDE hepatocytes were categorized as ‘damaged’, this fraction of hepatocytes more than doubled in TAA-treated mice (∼75%; Figure 6C). Interestingly, despite almost identical HDS tipping point values between CDE and TAA hepatocytes (Figure S12B), the relative contribution of senescent hepatocytes to all damaged hepatocytes was markedly elevated in TAA mice as compared to CDE mice (Figure 6C).

To better characterize these cell states, we determined genes that were differentially expressed between the senescent and not-senescent damaged hepatocytes (Figure 6D-6E, Tables S7-S8). We found 422 genes with significantly increased expression and 475 genes with significantly decreased expression in senescent damaged hepatocytes (adjusted p < 0.05, absolute log_2_(Fold Change) > 0.5, and expressed in at least 10% of senescent or not-senescent damaged hepatocytes). Of the 422 genes significantly upregulated among the senescent cells, 30 were part of the SHGS.

Genes upregulated in senescent damaged hepatocytes were involved in cell-adhesion, actin cytoskeleton remodeling, responses to wound healing, as well as apoptosis signaling, and cell growth (Figure S12H, Tables S6, S8). In hepatocytes the actin cytoskeleton is of great importance for maintaining the polarized cell structure and ensuring proper function of bile canaliculi.^97^ A key molecule in this context is Iqgap1, an actin-binding scaffold protein and signaling integration factor, which we found as specifically upregulated in senescent damaged hepatocytes (Figure 6E). Another example is Reticulon 4 (Rtn4 also known as Nogo-B), whose expression was also highly upregulated in senescent damaged hepatocytes, and which has been linked to diet-induced fatty liver disease.^98^ Further, we detected upregulation of Bcl2l1 with increasing damage, reaching its highest levels in the senescent damaged hepatocytes (Figure 6E). Bcl2l1 encodes for an apoptosis inhibitor and exerts protective functions in hepatocytes, providing insight into functional consequences of the senescent cell state.^99^ Genes down-regulated in the senescent hepatocytes were enriched for hepatic metabolic functions such as lipid metabolism (e.g. *Nudt7*,^100^ *Scp2*,^101^ *Lpin1*^102^), bile acid synthesis (e.g., *Cyp7a1*, *Cyp27a1*, *Abcb11*,^103^ and *Sult2a8*^104^), amino acid catabolism (e.g., *Sardh*^105^) and xenobiotic breakdown (e.g., *Cyp2e1*^106^), as well as insulin signaling (e.g., *Igf1*^107^). (Figure 6D, Figure S12I, Tables S6, S8). For example, the expression of *Cyp7a1* and *Cyp27a1,* that encode central enzymes of the canonical and alternative bile acid biosynthetic pathways, were first upregulated with damage, but then downregulated with entry into a senescent cell state (Figure 6D). Additional effectors of the bile acid synthesis pathway were first downregulated upon damage and displayed an exacerbated downregulation in senescent hepatocytes. This pattern was evident for the sulfotransferase Sult2a8, an enzyme responsible for the detoxification of bile acids through sulfonation and important to the maintenance of a homeostatic bile acid pool,^104,108^ and for *Abcb11,* which codes for the canalicular bile salt pump, the primary transporter for bile acid salt (Figure 6D).^103^ In summary, liver snRNA-seq of murine NASH models indicated a drastic downregulation of bile acid metabolism in senescent damaged hepatocytes, which differentiated senescent cells from non-senescent hepatocytes at similar damage levels. To confirm this dysregulation of the bile acid metabolism in senescent hepatocytes, we carried out additional experiments and analyses in HFD-NASH and CDAA-NASH mouse models. These models display phenotypes comparable to the CDE and TAA models analyzed previously.^109^ First, we re-analyzed scRNA-seq data from HFD-NASH and CDAA-NASH mice by calculating HDS and SHGS activity in hepatocytes.^109^ On average, hepatocytes in the CDAA-NASH model had greater SHGS activity and HDS levels compared to HFD-NASH cells (Figure S12J-S12K). Further, this analysis confirmed a relative cell composition between senescence and damage categories similar to CDE and TAA animals (Figure 6F). Importantly, in CDAA-NASH more than 25% of all hepatocytes realized a senescent cell state (Figure 6F). Next, we assessed the prevalence of senescent cells in CDAA-NASH, HFD-NASH and control livers using a senescence-associated beta-galactosidase (SABG) assay. Indeed, SAGB activity was visibly increased in CDAA-NASH samples when compared to HFD-NASH or control livers (Figure 6G), confirming increased levels of hepatocyte senescent cell states in CDAA-NASH mice. Subsequently, we confirmed that enzymes involved in bile acid metabolism (Cyp7a1 and Cyp27a1) were downregulated in CDAA-NASH mice along with increased senescence in this model. RT-qPCR showed that *Cyp7a1* and *Cyp27a1* mRNA levels were significantly downregulated in CDAA-NASH when compared to HFD-NASH mice (Figure 6H). Finally, we stained murine livers for CYP7A1 as the key and rate-limiting enzyme of canonical bile acid synthesis by immunofluorescence and assessed protein levels by densitometry (Figure 6I-6J). Again, downregulation of CYP7A1 in CDAA-NASH when compared to HFD-NASH or control livers, respectively, was evident. Taken together, we confirmed a metabolic shift towards decreased bile acid metabolism in senescent hepatocytes.

In summary, our observations support a model in which the emergence of a senescent phenotype in hepatocytes is in part damage-dependent, as a critical level of damage must be exceeded before an appreciable number of cells embark on a trajectory towards senescence. Yet damage alone does not fully explain why some cells become senescent and others do not (Figure 6K). Further, senescent, and not-senescent damaged hepatocytes constitute two distinct cellular states that are metabolically highly divergent. According to our data, senescent hepatocytes may be key contributors to the development of steatosis due to their dysregulated lipid metabolism and repression of bile acid production in particular.

## Discussion

Understanding how cellular damage accumulates and drives degenerative diseases remains one of the most pressing challenges in biomedical research. Here, we introduce a novel strategy that quantifies cellular damage at single-cell resolution, providing a powerful tool to dissect disease progression across diverse biological systems. This integrative experimental and computational strategy leverages cell-type-specific molecular markers for the systematic stratification of cells along a damage continuum, revealing fundamental mechanisms that underlie cellular dysfunction in chronic degenerative diseases. Unlike conventional transcriptomic analyses that rely on discrete cell-state classifications (e.g. ‘diseased’ *versus* ‘healthy’), our method captures progressive molecular deterioration, offering a new paradigm to study the molecular pathogenesis of degenerative disorders. Further, our approach capitalizes on previous studies when analyzing new data, because the PDS and HDS essentially ‘crystallize’ information from earlier transcriptomic studies. Our damage scores allow for generating cellular damage trajectories that carry a defined interpretation and meaning. Damage trajectories stand in contrast to the established pseudo-time trajectories, which have failed to detect the damage signal and are subject to speculative interpretations as to what molecular process they might represent. In addition, the possibility to align damage scores across studies enables the direct, quantitative comparison of damage trajectories between new data of a particular study with any earlier (single cell) transcriptomic study of the same disease. Thereby our approach also substantially advances mechanistic insight, because it facilitates the distinction of trigger-specific pathway changes from generic, damage-related changes.

The scores provide a robust quantitative readout of cellular damage, because marker genes were derived from a great diversity of studies and disease models and had to be reasonably expressed in the target cell type to ensure detectability in single-cell data. Importantly, the scoring is rank-based, which makes it resistant to technical variability between cells (e.g. library size). As dozens of genes are integrated, the impact of individual ‘’outlier’ genes is reduced. Robustness of the scores has been confirmed *via* (1) varying the number of marker genes, which did not have any relevant effect on the results, (2) leave-one-model-out cross validation, (3) validation using spatial transcriptomic data with matching histological information, (4) correlation of damage scores with clinical and morphological endpoints, such as ACR and SD length, (5) application in a different species (i.e. human), and (6) confirmation of findings with follow-up experiments.

Especially in case of single-cell data, the damage scores reflect a composite measure of damage occurring *in vivo* and during sample processing. Cell lysis and single cell processing in microfluidic devices might additionally damage cells, which potentially contributes to measured damage scores. Indeed, we noted that PDS obtained from single-cell data were on average slightly greater than PDS obtained from single-nucleus data. This observation is consistent with the notion that the handling of live cells can lead to extra damage along with respective transcriptional ‘traces’. Yet, as long as all cells are treated equally and in parallel, such a bias would not affect relative differences between cells. Importantly, we found that the small differences between scRNA-seq and snRNA-seq data did not compromise the comparability of scores between studies (Figure S8A).

Even though our strategy for selecting marker genes did not require that markers causally contribute to the disease, the composition of marker gene sets revealed exciting insight into the nature of cellular damage. In both cases (PDS and HDS) marker genes were enriched for highly cell-type specific genes that were consistently repressed under disease conditions (Table S2). This observation is consistent with the notion that the loss of cell type identity is a hallmark of cellular damage.^42,43,110,111^ On the other hand, very few markers were related to generic stress response, such as the Integrated Stress Response pathway. While this may come as a surprise, generic stress markers are not necessarily good damage markers. Stress responses may be protective, adaptive, or transient. If a cell responds to stress in a physiologically appropriate way, this may not indicate cellular damage. Rather, the opposite: a physiological stress response may be an indicator of high cellular fidelity. Thereby our work highlights an important distinction between cellular stress and cellular damage.

Applying this concept to podocytes and hepatocytes, we identified conserved, damage-associated molecular programs that operated across animal models and human disease. In podocytes, we uncovered a metabolic shift toward oxidative phosphorylation, likely reflecting the energy demands of cytoskeletal remodeling in response to damage. By mapping the damage continuum, we could differentiate model-specific behavior of energy metabolism and tight junction pathways. In addition, our data established that focal adhesion components exhibit both universal and trigger-specific responses to injury, offering new insight into the structural adaptions of podocytes under stress. While it was already established that changes in the circadian rhythm impact podocytes^87–89^, we now provide the first direct evidence that podocyte damage actively disrupts circadian gene regulation, revealing a bidirectional interplay between circadian control and podocyte health - a critical finding that underscores the role of cell-intrinsic circadian dysregulation in disease progression.

In hepatocytes, we showed that progressive damage determines cell state transitions, driving cells toward divergent cell states. Notably, we identified a tipping point in hepatocyte damage beyond which cells may enter a senescent state, leading to the repression of lipid and bile acid metabolism. These findings suggest that senescent hepatocytes may serve as drivers of tissue dysfunction, potentially fueling chronic inflammation and hepatocellular carcinoma (HCC) development.^94^ The ability to pinpoint this tipping point in two species represents a critical step toward defining therapeutic windows for intervention in liver disease progression.^112^

Beyond its biological insights, our damage-scoring framework has direct clinical implications. The fact that the damage scores were readily applicable to human single-cell and diverse spatial omics platforms underlines the clinical applicability of our concept. As such, the pathway fingerprints derived from our approach can serve as diagnostic tools for the molecular characterization of patient biopsies, enabling subtype-specific stratification of disease states. Furthermore, by performing cross-model analysis, we identified decreased Dpp4 expression levels as a conserved hallmark of podocyte damage, suggesting that Dpp4 inhibitors - currently used in type 2 diabetes - could be repurposed for podocyte disease therapy. We demonstrate the feasibility and utility of this concept for two very distinct diseases; one affecting proliferating, metabolically highly active cells and the other one affecting post-mitotic cells that primarily serve a physical function in their organ. Hence, this study establishes a universal blueprint for investigating degenerative diseases at a systems level, offering a scalable approach to quantify cellular damage across species and disease contexts.

The ability to apply our single-cell damage scores across *in vivo* omics modalities and cross-species datasets opens exciting new opportunities for decoding the mechanisms of chronic disease progression beyond the kidney and liver, with potential applications in neurodegeneration, fibrosis, and aging-related disorders. By shifting the focus from static disease endpoints to progressive, single-cell damage trajectories, our approach lays the foundation for a new generation of precision diagnostics and targeted therapies aimed at intercepting degenerative diseases at their earliest stages.

## Limitations of the study

The damage scores are RNA-based, which makes them readily applicable to a great diversity of platforms and assays, including single-cell and spatial transcriptomics data. However, this approach might be limited in capturing cellular state transitions that do not leave a clear transcriptomic signature, such as the terminal phases during apoptosis.

While this study clearly demonstrated how molecular damage scores can be utilized to obtain insight into disease mechanisms, future work needs to explore clinical applications more deeply. For example, our study did not assess the prognostic value of the damage scores or pathway signatures derived from them. Further, interpretation of the human data is constrained by data availability. For example, cellular damage in humans is expected to result from a combination of the disease (including genetic pre-disposition), age and lifestyle. While our study clearly shows the contribution of disease and age, we lacked information on lifestyle factors to also demonstrate their correlation with the PDS or HDS.

Finally, while the PDS and HDS represent a significant step forward, they are likely not yet perfect. As larger training datasets become available, incorporating machine learning-based approaches will be essential. The application of pre-trained foundational models tailored to specific diseases and cell types seems like a powerful and exciting extension of our framework.

## Resource availability

### Lead contact

Requests for materials should be addressed to the lead contact, Andreas Beyer

## Material availability

Unique reagents or materials generated in this study are available from the lead contact.

## Data availability

All sequencing data generated within the study has been deposited at Gene Expression Omnibus (GEO) under the accession number GSE289878, GSE289795, GSE289877 and are publicly available as of the date of publication.

Xenium spatial transcriptomic data and de-identified clinical metadata have been deposited in the Zenodo database and can be accessed at https://doi.org/10.5281/zenodo.15746500.

## Code availability

All code has been made available on GitHub at: https://github.com/beyergroup/PodocyteDamageScore https://github.com/beyergroup/HepatocyteDamageScore

## Acknowledgements

We thank Christoph Schell for valuable discussions and providing protein interaction networks and Martin Höhne for support with image quantification macros. The Wt1 knock-out mouse was a gift from Jordan A. Kreidberg. We thank the FACS & Imaging Core Facility at the Max Planck Institute for Biology of Ageing, Cologne and the CECAD Imaging Facility, Cologne for their support in FACS and imaging acquisition, respectively. We thank Simon Tröder and Branko Zevnik from the in-vivo Research Facility at CECAD Cologne for technical support in mouse breeding and transgenics. We thank S. Lück, P. Thaben, and A. Kramer for assistance with circadian bulk RNA-seq mouse experiments. We thank Vanessa Künstler for expert technical assistance in acquiring Xenium spatial transcriptomic data. H.C. and P.U.A. received support from the Cologne Graduate School of Ageing Research. This study was conducted within CRU329 at the University of Cologne and was supported by grants from the Deutsche Forschungsgemeinschaft (DFG, German Research Foundation) (CRC 1678 Project-ID 520471345 to R.U.M & A.B.; BE2603/10-1, BE2603/10-2 to A.B.; KA3217/5-1, KA3217/5-2, KA3217/6-1 to M.K.; MU3629/4-1 to R.U.M; BR6668/2-1 to F.B., BR2955/8-1 to P.T.B., SCHE1562/7-1, SCHE1562/7-2 to B.S.; BE2212/23-1, BE2212/23-2 to T.B.; CRC TRR 422 to T.B., A.B., F.B., P.T.B., L.B., M.K., R.U.M., B.S.). The work was further supported by the German Ministry for Science and Education by means of Research grants 0315899 to P.O.W. and 01EO2106 to F.B. as well as within the STOP-FSGS consortium 01GM1901E to P.T.B. and T.B..

## Author Contributions

T. P., P. U. A., and H. C. contributed equally to this work. M. K. and A. B. jointly supervised the project. T. P., P. U. A., H. C., M. K., and A. B. conceptualized the study. T. P. and P. U. A. curated data and performed computational analyses. H. C., M.-K. K., C. V., P. W., L. B., D. U.-J., C. K. carried out experiments and analyzed data. F. B., P. O. W., and R. U. M. generated data for circadian rhythm analyses. C. Ö., P. T. B., and B. S. generated mouse models. H. G. provided expertise in renal pathology, tissue handling resources, and prepared human samples for spatial transcriptomics. C. K. provided expertise in spatial transcriptomics, and spatial transcriptomics resources. T. P., P. U. A., H. C., M. K., and A. B. visualized the data and generated figures. F. T. W. provided expertise on liver biology and validated results. T. B. provided expertise on podocyte biology. T. P., P. U. A., H. C., T. B., M. K., and A. B. wrote the original manuscript draft and revised and edited the manuscript. All authors approved of the manuscript.

## Supplementary Figures and Tables

**Figure S1:**
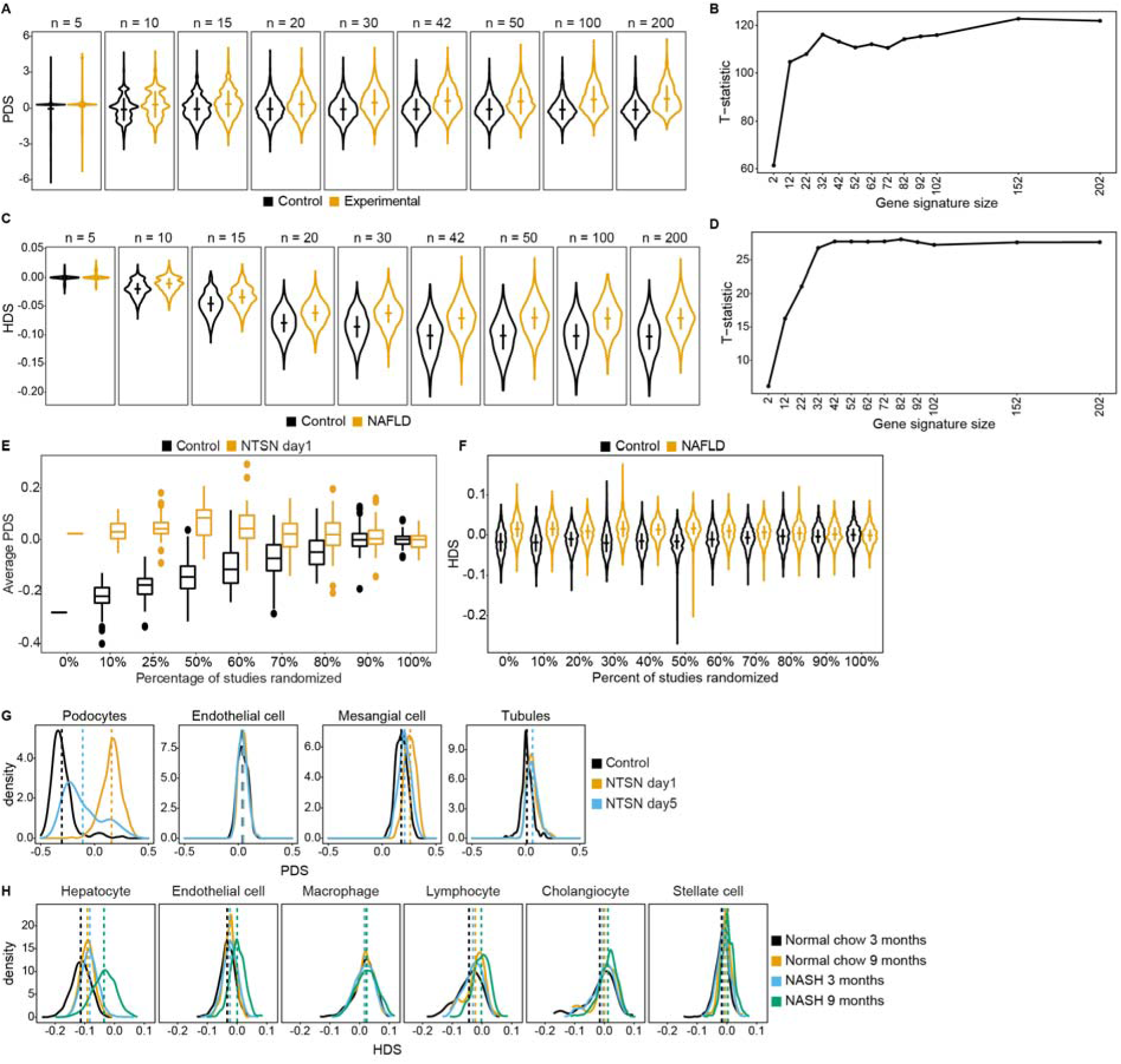
Robustness and specificity of PDS and HDS. A. Signature size test for PDS. Three novel snRNA-seq (Wt1^ko/wt^, Nphs2^mut^, Pdss2^mut^) and four public scRNA-seq datasets^61^ were used to test for the number of genes in the PDS needed to robustly differentiate control (black) and experimental (orange) conditions. PDS were scaled per study (Y-axis). Number of top genes (*n*) used per signature is indicated above. Insets in violin plots show the mean plus/minus 2 standard deviations. B. Signature size test shows absolute t-statistics of PDS comparisons between control and experimental samples in sc/snRNA-seq studies as a function of damage signature size (top *N* genes used; X-axis). Larger absolute t-statistics indicates stronger separation of experimental and control samples. C. Same as in A for HDS, except HDS is not scaled. Median HDS per distribution are indicated; whiskers represent the inter quartile range (IQR). HDS calculated for snRNA-seq of hepatocytes from 24-week-old mice with and without NAFLD-inducing diet (data from Liver Cell Atlas, GSE192742^9,48^). D. Same as in B for HDS. Signature size test shows absolute t-statistics of HDS comparisons between control and experimental samples in snRNA-seq data as a function of damage signature size (top *N* genes used; X-axis). Larger absolute t-statistics indicates stronger separation of experimental and control samples. E. PDS robustness test against noise in training data. Varying fractions of the training data were randomized to test how this would reduce the ability of the PDS to successfully distinguish experimental from control conditions in independent test data (see Methods). ScRNA-seq data from nephrotoxic serum nephritis (NTSN) model of podocyte damage^61^ was used for the test. 50 rounds of randomization were conducted for each ‘percentage of studies randomized’. Box plots show first (Q1), second, and third quartiles (Q) of the distribution of average PDS per cell across 50 randomization rounds per ‘percentage randomized’. Whiskers represent Q1 - 1.5 x interquartile range (IQR) and Q3 + 1.5 x IQR. Outlier cells are shown as dots. F. Same as in E, but for the HDS. Test was conducted on hepatocyte snRNA-seq data from 24-week-old mice fed with standard (control) or Western (NAFLD) diet from the Liver Cell Atlas (GSE192742^9,48^, same data as in C). HDS centered per run. Median HDS per distribution is indicated; whiskers represent IQR. G. Cell-type specificity test for the PDS. Panels show distribution of PDS in four major glomerular cell types (podocytes, endothelium, mesangium and tubules, color of lines reflect experimental condition (control and two damage time points). scRNA-seq data from the nephrotoxic nephritis model^61^. Vertical dashed lines represent the mean. H. Cell-type specificity test for the HDS. Panels show distribution of HDS in hepatocytes and five major liver cell types in a snRNA-seq data set^56^. Cell type annotation was adopted from the authors. Colour of density lines represent the condition of the mouse (n = 1). Vertical dashed lines represent the mean.

**Figure S2:**
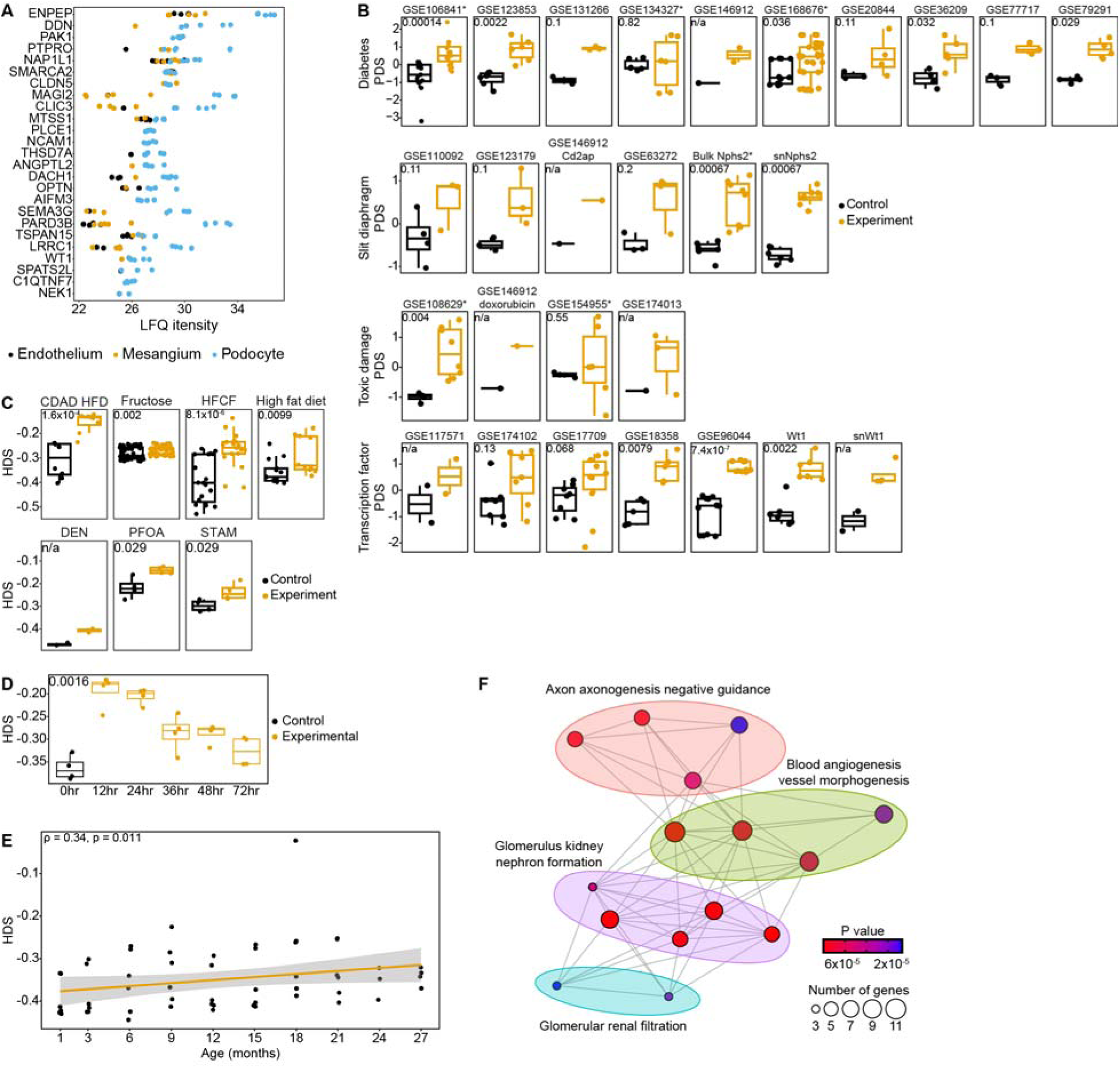
Cross validation of PDS and HDS. A. Protein expression of PDS marker genes in three types of mouse glomerular cells: podocytes (blue), endothelial cells (black) and mesangial cells (orange). Data is from glomerular proteomics as reported in PXD016238^13^. Non-scaled protein expression values (LFQ) are shown. B. Leave-one-dataset-out cross validation of the PDS. Each dataset of the indicated group of disease models were left out during the training phase of the PDS and used for testing the indicated disease model afterwards. Each dot represents the mean PDS across all cells of a sample. PDS were scaled and centered per study. Boxplots show first (Q1), second (Q2), and third quartiles (Q3) of the distribution. Whiskers represent Q1 - 1.5 x interquartile range (IQR) and Q3 + 1.5 x IQR. Outlier cells are shown as dots. P-value of Wilcoxon test is shown for all studies with at least 3 samples in both conditions. For studies GSE106841 (ageing series in diabetes), GSE134327, GSE168676, GSE108629, Bulk Nphs2, GSE108629, and GSE154955 diverse experimental animals (different age, treatments etc.) were summarized as ‘experiment’. C. Leave-one-model-out cross validation of the HDS. Procedure is the same as in B. Each dot represents the average HDS of one sample. CDAD HFD: choline-deficient, L-amino acid-defined, high-fat diet, HFCF: high fat, cholesterol, and fructose diet, DEN: Diethylnitrosamine (DEN)-induced carcinogenic liver injury, PFOA: perfluorooctanoic acid, STAM: mouse model that recapitulates human NASH progression to HCC and uses a combination of streptozotocin and high fat diet. Boxplots show the first (Q1), second (Q2), and third (Q3) quartiles of distribution. Whiskers show Q1 - 1.5 x IQR and Q3 + 1.5 x IQR. P-values of Wilcoxon rank test (experimental versus control conditions) are shown. D. Cross validation of the HDS on a time course dataset. Procedure is the same as in B Each dot represents a sample in time series murine liver samples post hepatic injury induced by acetaminophen administration (GSE111828^113^). Labels on x-axis represent the time in hours (hr) after acetaminophen administration. Untreated control samples are labeled with 0hr. P-value of Kruskal Wallis test shown. E. Leave-one-model-out cross validation of HDS. Procedure is same as in B. Each dot represents the HDS of a murine liver sample of an age group (in months on x-axis) from the Tabula Muris Senis data set (GSE132040^114,115^). Linear regression of the HDS as a function of age is shown in orange with confidence interval as gray shading. Correlation coefficient and p-value of Spearman correlation are indicated. F. Functional annotation of the podocyte damage marker set confirms its relations to the key podocyte functions. GO terms were grouped by semantic similarity. Nodes are colored by p-value of Fisher’s exact test and sized by the number of genes tested.

**Figure S3:**
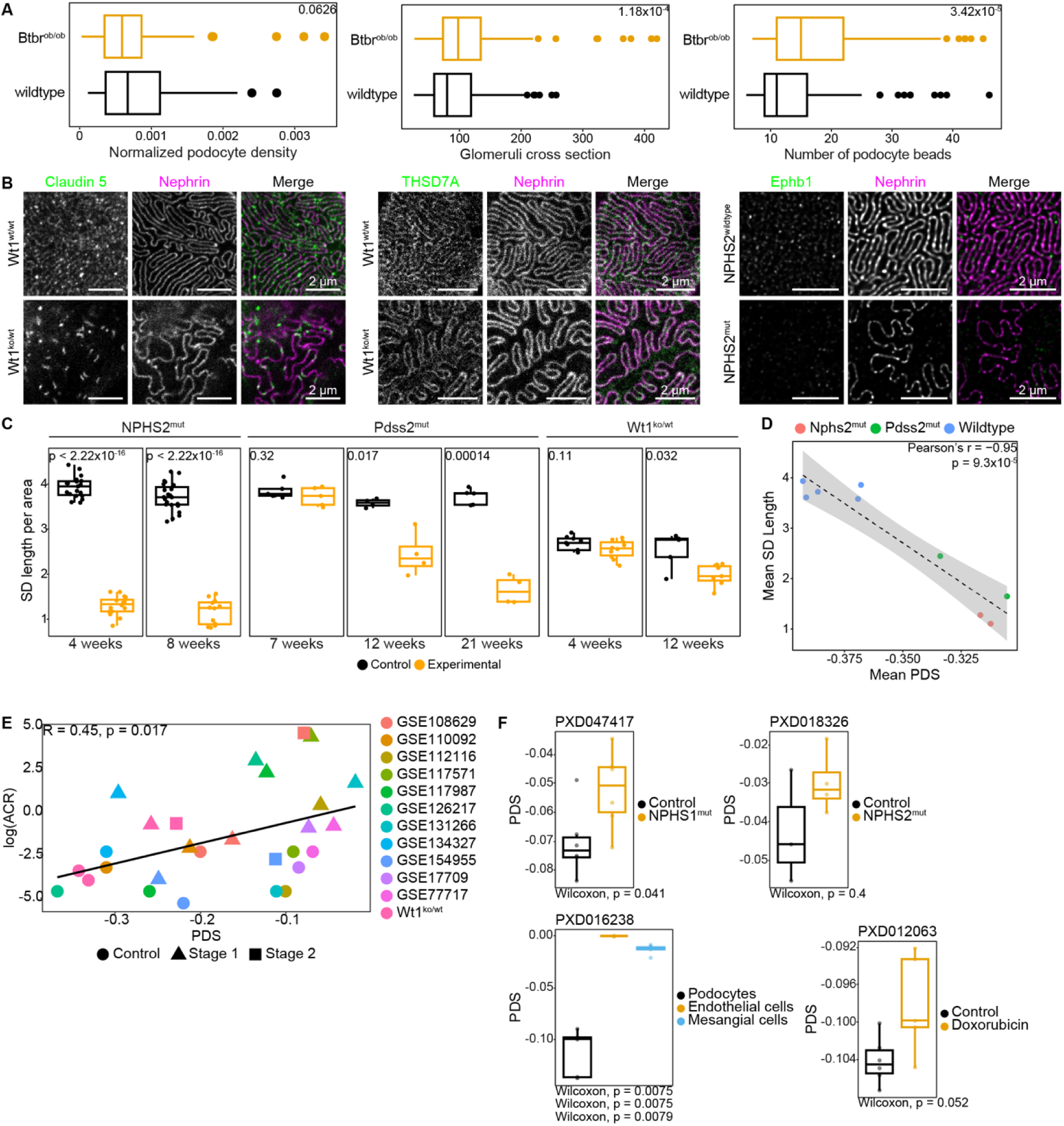
Functional validation of PDS. A. Boxplots showing glomerular size estimates, podocyte bead number, and normalized podocyte density for diabetic Btbr^ob/ob^ (orange) and control mice (black) as calculated on Slide-seqV2 spatial transcriptomics data provided in GSE190094^45^. Podocyte KNN filtration, glomerular identification and size estimation was performed using the programming code of the original publication^45^. Boxplots show quartiles (Q), with whiskers representing Q1 - 1.5 x interquartile range (IQR) and Q3 + 1.5 x IQR. Outlier cells are represented by dots. Values in top right are p-values of Wilcoxon rank sum test of the difference between experimental and control groups. B. STED of 12-week-old Wt1^ko/wt^ or control mice co-stained for nephrin to visualize the slit diaphragm and PDS marker proteins including Claudin 5 and THSD7A. STED of 4-week-old Nphs2^mut^ or control mice was co-stained for nephrin and the Eph B1 receptor. All PDS marker proteins localize to the slit diaphragms. In Wt1^ko/wt^ mice, Claudin 5 forms aggregates at the slit diaphragms. In Nphs2^mut^ mice, the expression of Eph B1 is downregulated upon podocyte damage. C. Quantification of SD length per area for Nphs2^mut^ (n = 19 for 4-week control, n = 16 for 4-week experimental, n = 23 for 8-week control, n = 12 for 8-week experimental), Pdss2^mut^ (n = 10 for 7-week, n = 8 for 12-week, n = 9 for 21-week) and Wt1^ko/wt^ (n = 9 for 4-week control, n = 10 for 4-week experimental, n = 5 for 12-week control, n = 7 for 12-week experimental) control and experimental animals at the indicated age. P-values of t-test are shown between control and experimental groups. D. Correlation plot of average PDS and average SD length per area for wildtype and experimental Nphs2^mut^ (Grouped control: n = 2 for 4-week, n = 1 for 6-week, n = 1 for 8-week, n = 2 for 12-week; Grouped experimental: n = 2 for each of 4-week, 6-week, 8-week and 12-week) and Pdss2^mut^ (Grouped control: n = 2 for each of 6-week, 12-week and 21-week; Groupe experimental: n = 2 for 6-week, n = 2 for 12-week and n = 1 for 21-week) mouse groups. Linear regression of PDS correlated with SD length shown in dashed line with 95% confidence interval colored in grey. Coefficient and p-value for Pearson’s correlation are shown. E. Correlation of urinary albumin/creatinine ratio (ACR) and PDS in bulk microarray and bulk RNA-seq data. Wt1^ko/wt^ represents bulk RNA-seq data published in ^116^. Stage 1 and 2 indicate datasets, in which longitudinal ACR data was available. F. Boxplots show PDS calculated for various conditions and genotypes in 4 proteomics datasets. Individual samples are shown as dots. P-value of Willcoxon test of condition/genotype effect are provided below each panel. ProteomeXchange identifiers (PXDs) of the corresponding datasets are indicated^13,46,50,51^.

**Figure S4:**
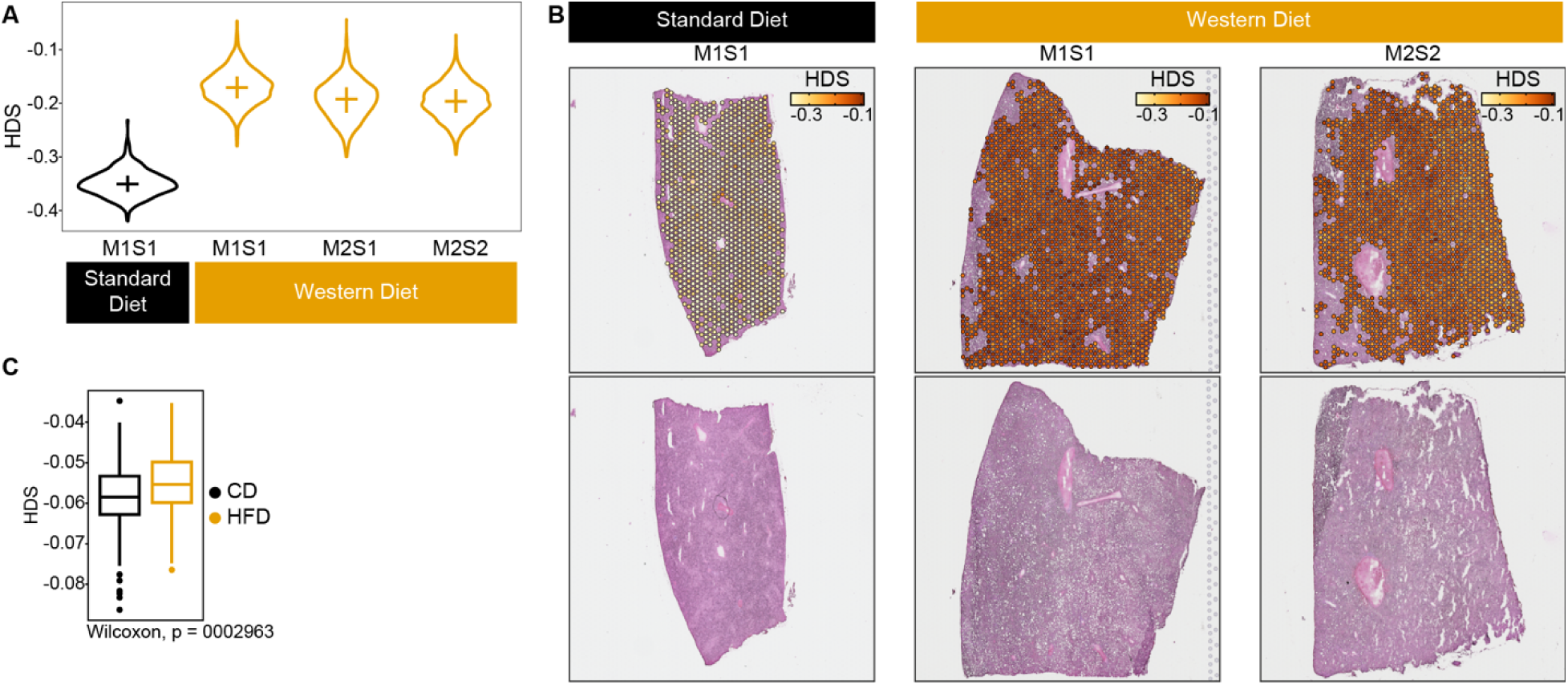
Functional validation of the HDS. A. Violin plots of HDS distribution in all murine spatial transcriptomic 10X Visium samples from the Liver Cell Atlas (GSE192742^9,48^) comparing western (n = 3) to standard diet (n = 1). M1S1 = Mouse 1 Sample 1, M2S1 = Mouse 2 Sample 1, M2S2 = Mouse 2 Sample 2. Horizontal lines show the median and vertical lines the IQR of the distributions. Statistical comparison not appropriate due to n = 1 per “Standard Diet” group. B. 10X Visium spatial transcriptomics of murine liver from the Liver Cell Atlas (GSE192742^9,48^). Spatial plots of HDS distribution in spatial transcriptomic not previously shown in Fig. 1. Color range: floor = -0.37, ceiling = -0.09. P-value for Wilcoxon test shown. C. Confirmation of HDS on murine liver proteomics data^52^. Mice fed a high fat diet (HFD) compared to mice fed a control diet (CD), (n = 315 across conditions, different strains, age range between 157 and 785 days). P-value for one-sided Wilcoxon test is shown.

**Figure S5:**
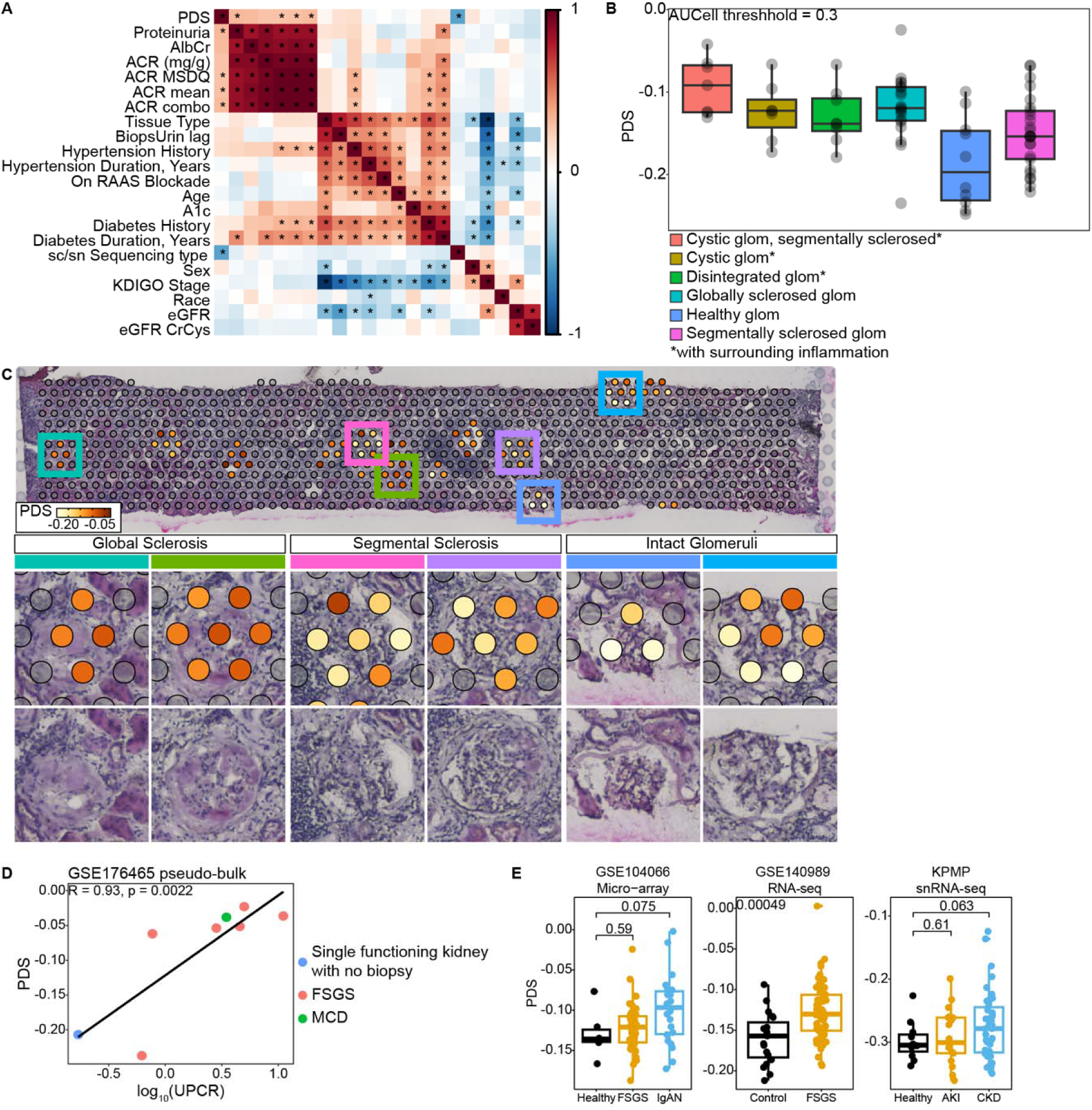
Validation of the PDS on human data. A. Heatmap of spearman correlation between PDS and detailed clinical traits from patients with acute kidney injury (AKI) and CKD. Anonymized patient data, sc/snRNA-seq data used in the analysis were retrieved from the KPMP database. Stars denote cells with p-value of Spearman correlation < 0.05. B. Boxplot shows PDS in further glomeruli as annotated manually in the histology image shown in the next panel, PDS measurements based on KPMP spatial transcriptomics dataset,^3^ patient ID 29-10282, accessed at https://atlas.kpmp.org/. C. 10x Visium spatial transcriptomic data on a human kidney biopsy from the KPMP registry (patient ID 29-10282)^3^ showing an FSGS phenotype. PDS values were computed only for Visium spots with podocyte signals as determined by Wt1 and Nphs2 expression. Selected glomeruli are marked in the upper panel with respective colors. High magnification images show various degrees of glomerular sclerosis with PDS gradients corresponding to degree of sclerosis. See boxplots for PDS distribution as by glomerular damage in the previous panel. D. Relationship between PDS calculated on human urinary podocyte snRNA-seq data and urinary protein creatinine ratio (UPCR) in patients suffering from either minimal change disease (MCD) or focal and segmental glomerulosclerosis (FSGS) as reported in GSE176465.^54^ Data from 8 patients with at least three measured podocytes each was used. E. Validation of PDS on human transcriptomic data obtained from healthy patients (CON) and from patients suffering from FSGS, IgA nephropathy (IgAN), acute kidney injury (AKI), and chronic kidney disease (CKD). Samples were acquired and reported in transcriptomic modalities and studies as indicated on top of the plots. Podocytes from snRNA-seq datasets were aggregated in pseudo-bulk samples. P-values of Wilcoxon rank sum test comparing experimental and control samples are shown.

**Figure S6:**
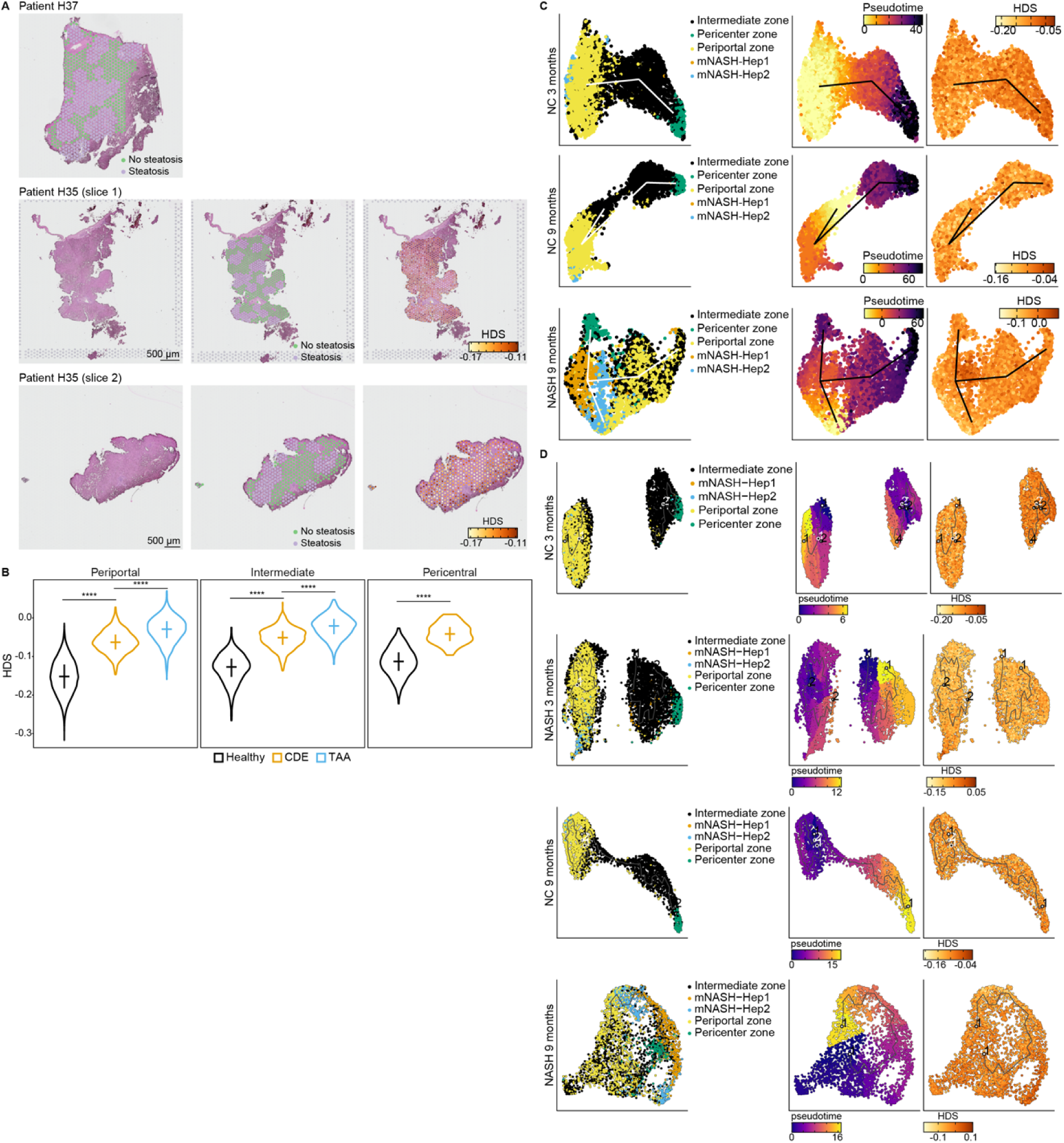
Validation of HDS using single nucleus and spatial transcriptomics. A. HDS calculation on 10x Visium human spatial transcriptomic data from the Liver Cell Atlas (GSE192742^9,48^). Only subjects with liver steatosis were analyzed. Top panel: manual assignment of steatotic vs non-steatotic areas as used for Figure 3D. Bottom two rows: Left panel: histology, Middle panel: assignment of steatotic vs non-steatotic areas, right panel: HDS per Visium spot overlay. Spots representing hepatocytes were manually annotated. Color scale represents the 10%- and 90%-quantiles of the HDS distribution across all samples. Spots in steatotic regions: light purple, spots in non-steatotic regions: green. B. HDS distributions by liver zonation in snRNA-seq data from mice subjected to three different treatments. Healthy: normal chow; CDE: choline-deficient, ethionine-supplemented (CDE) diet; TAA: thioacetamide (TAA) in the drinking water (GSE200366^96^) (n = 3 biological replicates per treatment). Same data is in Figure 6A. Hepatocyte zone annotation as provided by authors of data set. Pericentral hepatocytes were not detected in TAA-treated animals. Horizontal lines show median of the distribution and vertical lines represent the IQR. ****p < 2 x 10^-16^ in one sided Wilcoxon rank-sum test. C. Analysis of HDS versus pseudo-time trajectories with TSCAN. UMAPs for further samples from the same study as shown in Figure 3G. Each dot represents a hepatocyte colored either by liver zone (right), pseudo-time (PT) (middle) and HDS (left). Hepatocyte zone annotation from original publication^56^. MNASH-Hep1/2: classification from original publication corresponding to damaged hepatocytes whose zone of origin could not be determined according to the authors. PT trajectory inference is based on clusters identified using Seurat after quality control cell filtering, normalization with Seurat’s SCTransform and dimensionality reduction. PT is dominated by liver zonation, while the HDS is independent of it. Top: Normal Chow 3-month-old, middle: NASH-inducing diet 9-month-old, bottom: Normal Chow 9-month-old mouse. D. Analysis of HDS versus pseudo-time trajectories with Monocle3. Same data and legends as in C. Trajectory inference is based on clusters identified using Monocle3 default processing and clustering pipeline. PT trajectories and clustering are dominated by liver zonation also in Monocle3 analysis. HDS is not reflected by PT trajectories.

**Figure S7:**
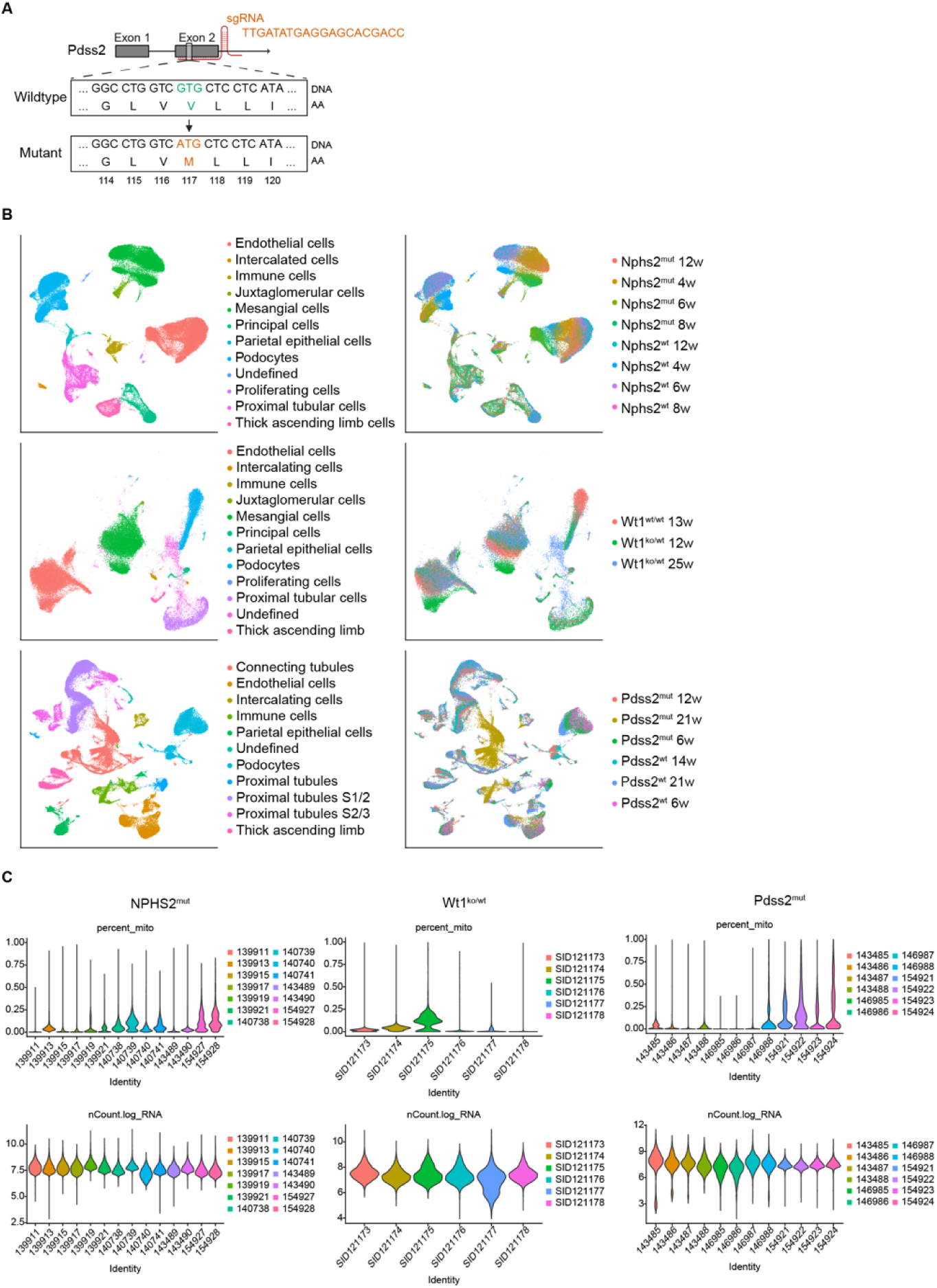
Glomerular snRNA-seq of Nphs2^mut^, Wt1^ko/wt^ and Pdss2^mut^ mice. A. CRISPR strategy for the generation of the Pdss2^mut^ mice. Exon 2 of Pdss2 is targeted using single guide RNA (sgRNA) to generate a point mutation which exchanges valine (V) at amino acid (AA) position 117 to methionine (M). B. UMAP of preprocessed snRNA-seq data colored by cell types (left) or condition (right) of NPHS2^mut^ (top), Wt1^ko/wt^ (middle) or Pdss2^mut^ (bottom) datasets. UMI counts corrected for ambient RNA, filtered and normalized (see Methods), are used for this and the next panels. C. Percent mitochondria (top) and log transformed number of RNA counts (bottom) for individual snRNA-seq samples of NPHS2^mut^ (left), Wt1^ko/wt^ (middle) or Pdss2^mut^ (right) datasets.

**Figure S8:**
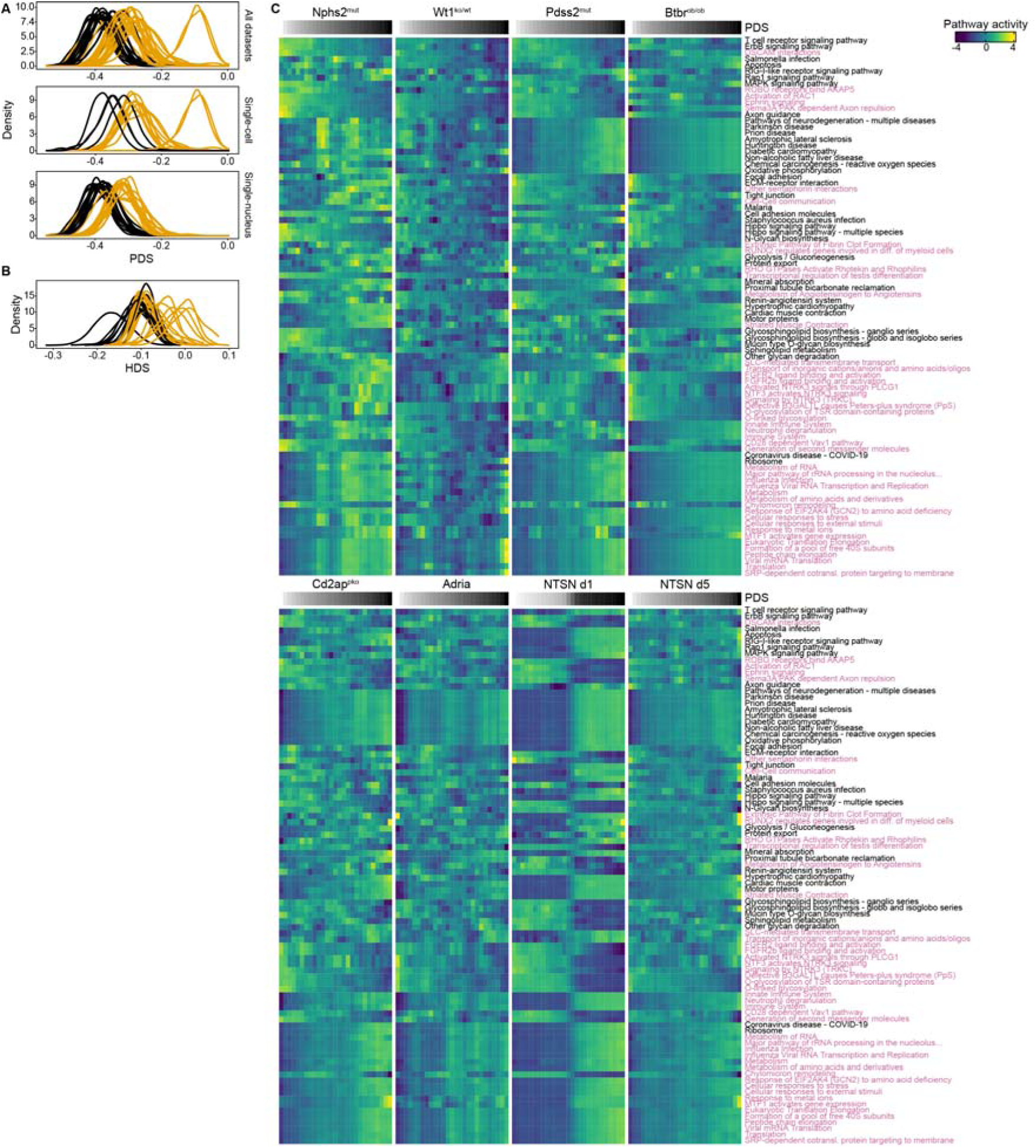
Pathway fingerprints of all FSGS sc/sn-RNA-seq models. A. Density plots show greater variation of PDS in experimental compared to control samples. PDS densities of individual samples are aligned by x-axis. Black: controls, orange: experimental conditions. Top panel shows results for all sc/snRNA-seq samples, bottom two panels show results for single-cell and single-nuclei datasets separately. Wilcoxon Rank Sum test confirms (p-value = 2.5x10^-11^) that controls are significantly better aligned than experimental samples (see method section Damage score alignment test). B. Density plots show greater variation of HDS in experimental compared to control samples. HDS densities from three sn/scRNA-seq studies across different liver damage models (NAFLD, NASH, ageing, GSE192742^9,48^, GSE200366^96^, GSE189600^56^). Black: controls, orange: experimental samples. Wilcoxon rank sum test confirms that control samples are aligned significantly better than experimental samples (p-value = 2.7x10^-4^). C. Full PDS pathway fingerprints for sn/scRNA-seq data from all studies analyzed. Cells/nuclei were sorted by increasing PDS (left to right, gray color scale). Pathway activities for KEGG (black) and Reactome (magenta) pathways were calculated using AUCell (see Methods). A total of 85 pathways - a union of 10 pathways that most significantly (adjusted p-value < 0.05) corelate with PDS in each model - is used.

**Figure S9:**
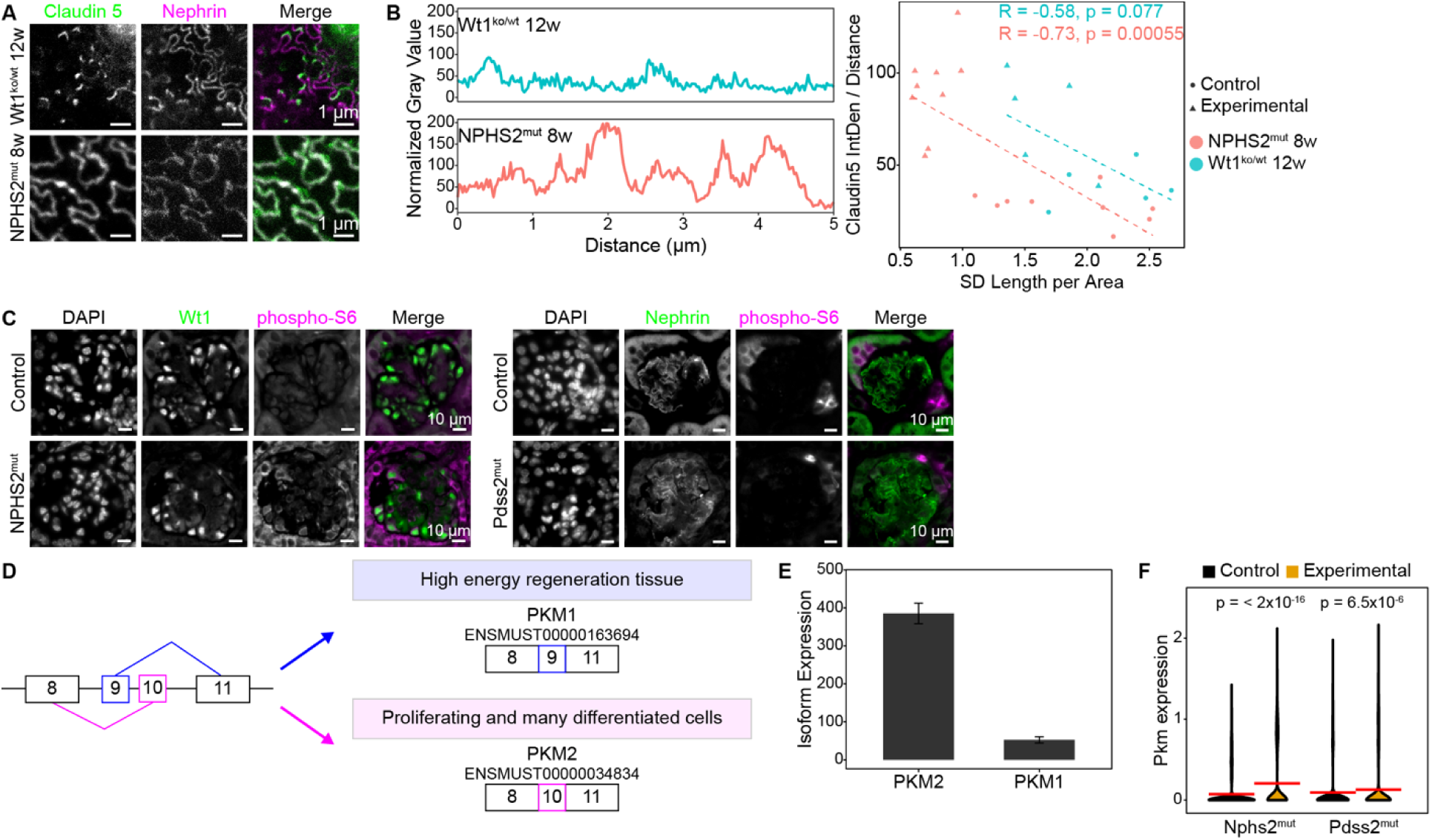
Validation of the PDS pathway fingerprints. A. Representative STED microscopy of 12-week-old Wt1^ko/wt^ or 8-week-old NPHS2^mut^ kidneys co-stained for claudin 5 (green) and nephrin (magenta) to assess the aggregation of claudin 5 at the podocyte slit diaphragm. B. Representative normalized line profiles of claudin-5 fluorescence intensity along manually traced slit diaphragms, illustrating differences in intensity and focal aggregation of claudin-5 along equal-length segments in NPHS2^mut^ (8-week-old) and Wt1^ko/wt^ (12-week-old) experimental mice. Right: Correlation plot of normalized claudin-5 fluorescence intensity to average SD length per area for 8-week-old NPHS2^mut^ (n = 9 for control, n = 9 for experimental) and 12-week-old Wt1^ko/wt^ (n = 5 for control, n = 5 for experimental) control and experimental animals. Correlation coefficient and p-values in Spearman are shown. C. Representative immunofluorescence staining of control and experimental kidney tissue from either NPHS2^mut^ 8-week-old (left) or Pdss2^mut^ 21-week-old (right) showing DAPI (blue), nephrin (green) and phospho-S6 (magenta). Phospho-S6 expression is upregulated in podocytes in NPHS2^mut^ mice. D. Cartoon of PKM1 and PKM2 isoforms and related dominant isoforms in respective cells or tissues. PKM2 is the dominant isoform in podocytes. E. Barplot of the expression of PKM1 and PKM2 isoforms in healthy mouse glomeruli, from bulk glomerular RNA-seq of Nphs2^mut^ mice. Error-bars indicate 95% confidence intervals F. Violin plot of expression of PKM in murine podocytes from Nphs2^mut^ and Pdss2^mut^ snRNA-seq experiments. Red horizontal lines indicate mean. P-value of Wilcoxon test of the genotype effect is displayed.

**Figure S10:**
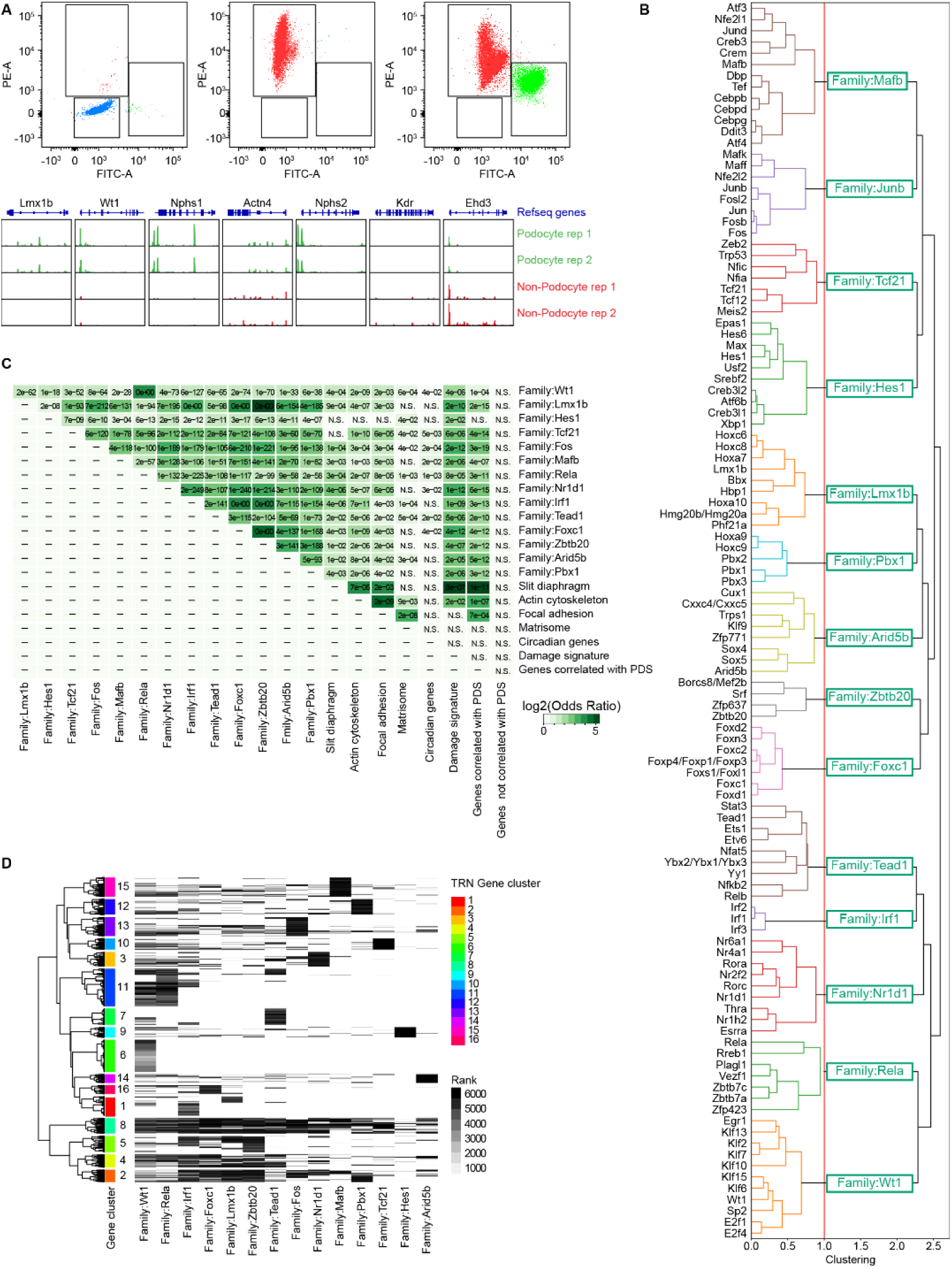
Development of the podocyte-specific transcription regulatory network and identification of podocyte damage pathways. A. Top: Fluorescence-activated cell sorting plot of glomerular nuclei extracted from nTnG;Pod:cre mice that express no nuclear fluorescence reporter (left), only tdTomato+ reporter (middle) or both GFP+/tdTomato+ reporter (right). Sorted nuclei were collected and used for the generation of cell type-specific ATAC-seq (see Methods). Bottom: genome browser tracks at podocyte-specific loci (Lmx1b, Wt1, Nphs1, Actn4, Nphs2) show accessible chromatin in podocytes, whereas those at other glomerular cell type-specific loci (Kdr, Ehd3) show accessible chromatin in non-podocytes. B. Dendrogram of hierarchical clustering of TF motifs (PWMs). The dendrogram is cut (red line) to produce 14 TF clusters (families), each one is named by the representative motif. C. Heatmap shows podocyte specific TRN: 14 regulatory TF-clusters (each representing a group of TFs with similar PWMs) are in columns and 6360 of their target genes expressed in podocytes are in rows. Shades of grey color shows ranks of predicted interaction score (from TFtargetCaller prediction), black and white color means the strongest and no predicted interaction, corresponding dendrogram on the left shows hierarchical clustering of genes. The dendrogram is cut to produce 16 clusters (annotated with rainbow colors), that represent common and unique targets of each of 14 regulatory modules D. Heatmap of overlaps of 14 TF regulons and podocyte gene sets used in Figure 5B. Color shows log2 odds ratio of the Fisher’s exact test, which represents the strength of association, cell labels show labeled with adjusted P-values of the test (if padj < 0.05, otherwise “NS”).

**Figure S11:**
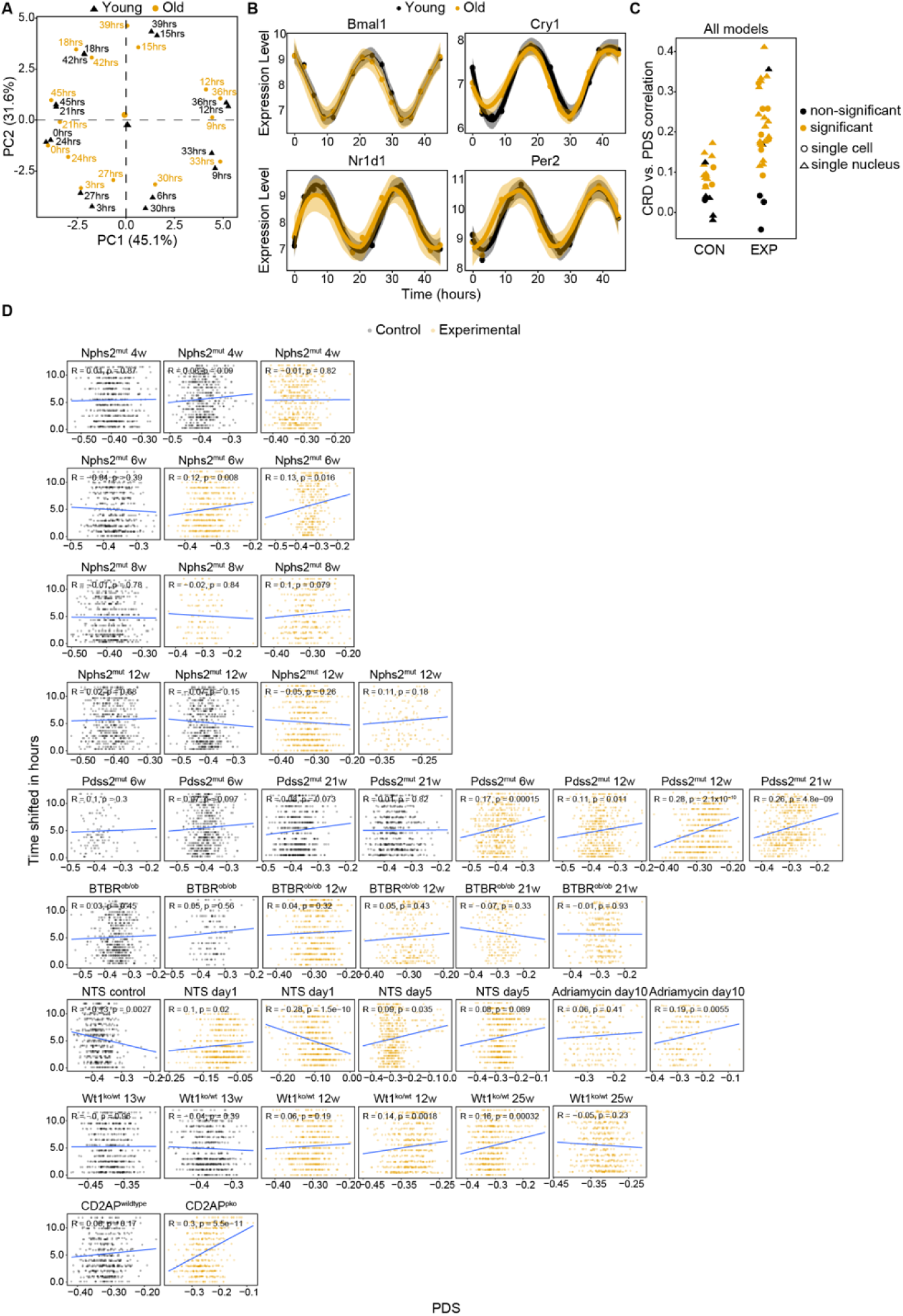
Transcriptome analysis of circadian gene expression. A. PCA of circadian bulk RNA-seq samples from whole kidneys. Each symbol represents a pooled replicate of 3 individual mice (yellow: age 78 weeks, black: age 8 weeks). Samples were taken at zeitgeber times 0, 3, … 21 hours during 12-h/12-h light–dark cycles. Replicates per group (n = 2) are presented in serial fashion resulting in a 48h period. B. Expression of circadian genes Bmal1, Cry1, Nr1d1, and Per2 in the same dataset as in panel a. Y-axis - gene expression value normalized by DEseq2 rlog function, X-axis – zeitgeber time, in hours. Replicates per group (n = 2) are represented in serial fashion resulting in a 48h period. Solid line indicates an estimate of the conditional mean function, shades indicate 95% confidence interval of estimates. Yellow: age 78 weeks, black: age 8 wks. C. Coefficients of Spearman rank correlation between PDS and CRD calculated based on podocyte-specific circadian genes^87^, for all individual control (CON) and experimental (EXP) samples of seven different mouse models of podocyte damage. Color distinguishes if the CRD to PDS correlation was significant (p < 0.05). D. Absolute deviation between circadian time of each cell from the reference sample time (Y-axis) of the respective sample as a function of the PDS (X-axis). Regression line and parameters for Spearman rank correlation are shown. Black and orange color of dots distinguish cells of control and experimental samples. Circadian time was calculated using a Tempo algorithm,^90^ Clock was used as a reference gene.

**Figure S12:**
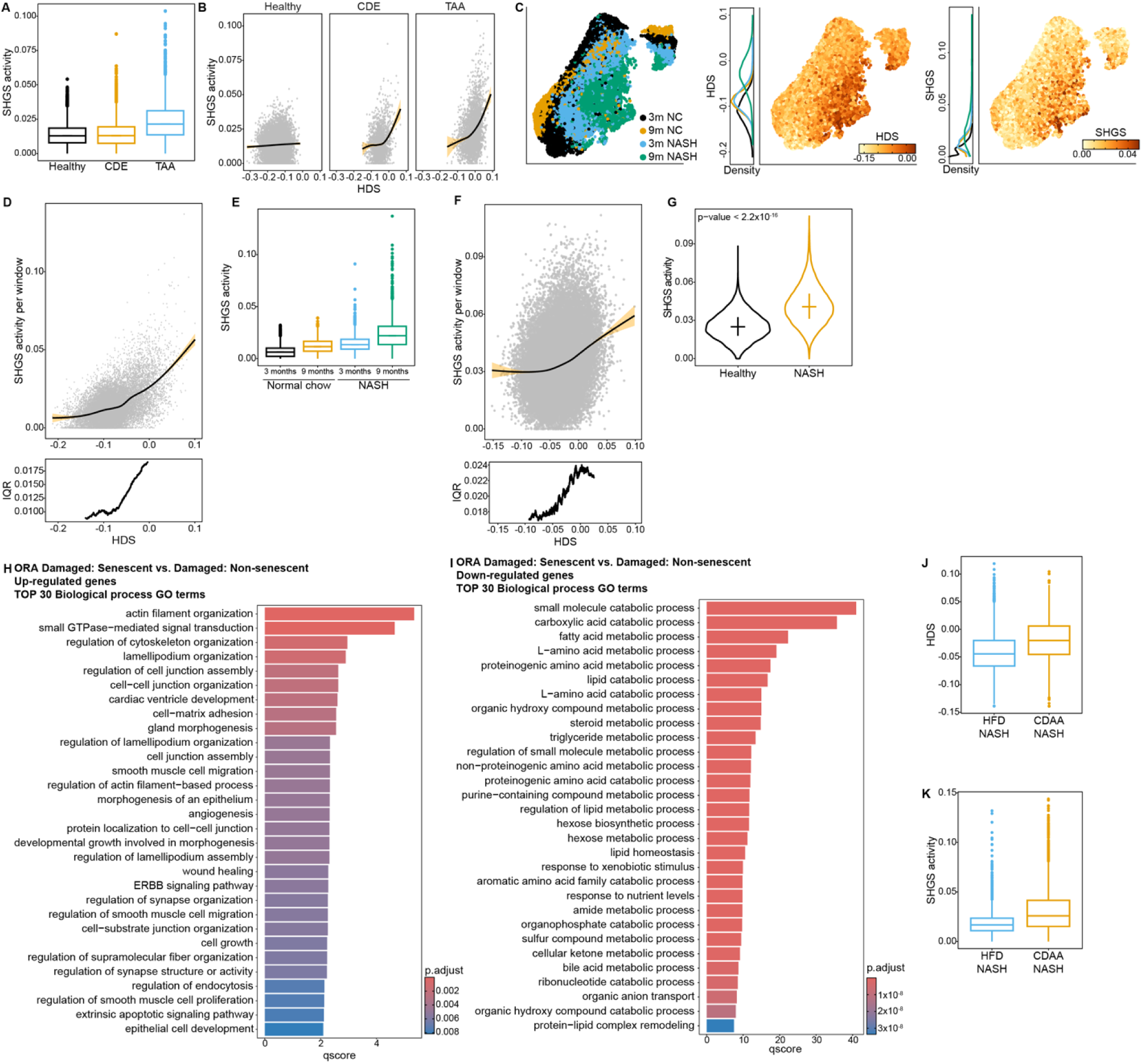
Analysis of hepatocyte cell state transitions. A. Boxplots showing activity distributions of the Senescent Hepatocyte Gene Signature (SHGS) in snRNA-seq data of mice subjected to three different treatments: Healthy: normal chow; CDE: choline-deficient, ethionine-supplemented (CDE) diet; TAA: thioacetamide (TAA) in the drinking water (GSE200366,^96^ n = 3 biological replicates per treatment, same data as in Figure 6A). Wilcoxon rank sum test for both, comparison of TAA to controls and TAA to CDE p < 2 x 10^-16^. Wilcoxon rank sum test for comparison of CDE to controls was not significant (p-value = 0.052). B. SHGS activity score versus HDS, separated by model. Same data as in Figure 6A-6B and panel A, but separated by condition). Each dot represents a hepatocyte. Solid line: Generalized linear model line showcases behavior of SHGS activity with increasing HDS. A tipping point pattern with consistent HDS threshold level is evident. Orange: confidence interval of regression line. X-axis represents the same range of HDS for all three plots. C. UMAP of merged snRNA-seq samples from mice held in four different conditions (n = 1): Black: 3-month-old fed normal chow (3m NC); Orange: 9-month-old fed normal chow (9m NC): Light blue: 3-month-old fed NASH inducing diet (3m NASH); Green: 9-month-old fed NASH inducing diet (9m NASH). Left: UMAP of hepatocytes colored by condition. Middle: hepatocytes colored by HDS. The color range for the HDS has floor and ceiling values corresponding to the 1%- and 99%-quantiles of the HDS distribution across all samples. Right: Hepatocytes colored by SHGS activity score.^94^ Density curves of HDS and SHGS activity score are shown per condition, color of line represents condition. Significance tests not appropriate because of sample size (n = 1). D. SHGS activity score versus HDS in NASH murine snRNA-seq (same data as in panel C). Top: SHGS activity score for each hepatocyte. Solid line: Generalized linear model line showcases behavior of SHGS activity with increasing HDS. Orange: confidence interval of regression line. Bottom: IQR of SHGS activity calculated per 2000-hepatocytes sliding window (y-axis) plotted against mean HDS per sliding window. E. Boxplots show distribution of SHGS activity per hepatocyte in each condition (n = 1). Same data and legend as in panel C. Statistical significance test not possible due to n =1. F. SHGS activity score versus HDS in human snRNA-seq data set of healthy and NASH patients (n = 3, same dataset as in Figure 3A). Top: SHGS activity score for each hepatocyte. Solid line: Generalized linear model line showcases behavior of SHGS activity with increasing HDS. Orange: confidence interval of regression line. Bottom: IQR of SHGS activity calculated per 2000-hepatocytes sliding window (y-axis) plotted against mean HDS per sliding window. G. Violin plots show distribution of SHGS activity for all hepatocytes per patient health status (black: healthy, orange: NASH, n = 3). Same dataset as in Figure 3A. Horizontal lines mark the median, and vertical lines the IQR. H. Biological Process (BP) Gene Ontology (GO) over-representation analysis (ORA) of genes significantly differentially up-regulated (p-value < 0.05 and LFC < -0.05) in damaged, senescent vs. Damaged, not-senescent hepatocytes. ORA results were summarized to avoid redundancy using the ClusterProfile simplify() function with default setttings. Barplot includes top 30 terms kept after filtering for adjusted p-value < 0.01. I. BP-GO-ORA of genes significantly differentially down-regulated (p-value < 0.05 and LFC > 0.05) in damaged-senescent vs. damaged-not-senescent hepatocytes. ORA results were summarized, and bar plot created, as describe in H. J. Boxplots of HDS distribution computed on scRNA-seq data from murine samples subject to either HFD-NASH or CDAA-NASH condition (n = 1).^109^ K. Boxplots of SHGS activity distribution computed on scRNA-seq data from murine samples subject to either HFD-NASH or CDAA-NASH condition (n = 1).^109^

**Table S1**: Disease models used for the kidney (PDS) and liver (HDS) analysis. https://docs.google.com/spreadsheets/d/10iI-gT9rHKqAnAGcTxrU9kCCoZ_bT4z-/edit?usp=drive_link&ouid=104332495390310243054&rtpof=true&sd=true

**Table S2**: Top 42 marker genes for PDS and HDS. https://docs.google.com/spreadsheets/d/1IJTCjhsYDoTGfaw_lx2jndaaR7Kjc9w8/edit?usp=drive_link&ouid=104332495390310243054&rtpof=true&sd=true

**Table S3:** Functional characterization of top 42 HDS markers genes by GO term overrepresentation analysis. https://docs.google.com/spreadsheets/d/1XWbiI0G_cGqRnD_IeYgK2rQDh83jcSa49al1OvxNrAk/edit?gid=0#gid=0

**Table S4**: Description of FSGS damage models and corresponding scRNA-seq datasets used in the manuscript. https://docs.google.com/document/d/1kR1WHLUGVFdD5UfkMLIYnqQCpISb61yg/edit?usp=drive_link&ouid=104332495390310243054&rtpof=true&sd=true

**Table S5:** List of 152 podocyte-specific TF target genes significantly correlated with the PDS. https://docs.google.com/spreadsheets/d/1mDy4L6pT4ndOS_yqxGM8iz_GN7Vpt-H6/edit?usp=drive_link&ouid=104332495390310243054&rtpof=true&sd=true

**Table S6:** GO term over-representation analysis of genes significantly (ajusted p-value < 0.01) down- or up-regulated in damaged senescent-like hepatocytes compared to damaged not-senescent-like hepatocytes. https://docs.google.com/spreadsheets/d/1Grq4roNyS3url6QX-Huq8OVMCUZkfxO2onkTY74ASYI/edit?usp=sharing

**Table S7:** Genes differentially upregulated in damaged-senescent hepatocytes compared to damaged-not-senescent hepatocytes. Only the genes passing filtering criteria are included. https://docs.google.com/spreadsheets/d/10hyJ8VkJfIKCuw5BKN-gYMHcX_vOLdjTjxO993vG4Ow/edit?usp=sharing

**Table S8:** Genes differentially downregulated in damaged-senescent hepatocytes compared to damaged-not-senescent hepatocytes. Only the genes passing filtering criteria are included. https://docs.google.com/spreadsheets/d/1a0e2Je8OywrumdjX6BT7mtS0e1LeGbJYGPmx0BsXUBY/edit?usp=drive_link

## STAR□Methods

### Experimental model and study participant details

#### Animals

All animals were housed and maintained in the CECAD *in vivo* Research Facility, University of Cologne, or in a barrier-protected facility of the MPI for Metabolism Research, Cologne. All mice were kept under a 12-hour light/dark cycle and had *ad libitum* access to food and water. All mice were maintained on C57BL6 (The Jackson Laboratory, Strain#000664) background except for Wt1^ko/wt^ mice, which were kept on FVB/N (The Jackson Laboratory, Strain#001800) background. NPHS2^mut^ mice have been published.^46^ Pdss2^mut^ mice were generated as previously published by the CECAD *in vivo* Research Facility (Figure S7A).^117^ The Wt1^ko/wt^ mice were previously described.^118^ Mice carrying the nuclear fluorescent reporter (nTnG, The Jackson Laboratory, Strain#:023035) were crossed to mice carrying NPHS2 Cre (Pod:cre, The Jackson Laboratory, Strain#: 008205),^119^ to generate podocyte-specific expression of nuclear GFP (nTnG;Pod:cre). To induce liver phenotypes, the following experimental diets were fed to *Stat3^fl/f^*^l^ (The Jackson Laboratory, Strain#:016923),^120^ or *NIK^fl/fl^* mice^121^ (in the context of this study considered as wildtype mice) mice starting at the age of 6 weeks: EF Control to NASH diet (CD: E15767-040, ssniff Spezialdiäten GmbH) containing 65% calories from carbohydrates, 22% of calories from protein, and 13% of calories from fat; EF NASH diet (HFD-NASH: E15766-340, ssniff Spezialdiäten GmbH) containing 42% calories from carbohydrates, 18% of calories from protein, and 40% of calories from fat; Choline free, lard, 1% cholesterol diet (CDAA-NASH: S2159-E042, ssniff Spezialdiäten GmbH) containing 58% calories from carbohydrates, 11% of calories from protein, and 31% of calories from fat. Controls received CD or NCD (R/M-H; ssniff Spezialdiäten GmbH), as indicated.

Animal experiments used age-matched littermates at indicated weeks of age. All mouse experiments were performed with approval from the Animal Care Committee of the University of Cologne and LANUV NRW (Landesamt für Natur, Umwelt und Verbraucherschutz Nordrhein-Westfalen, State Agency for Nature, Environment and Consumer Protection North Rhine-Westphalia) for this study.

To analyze circadian gene expression, two groups of male C57BL/6J mice aged 8 weeks and 78 weeks were acclimated for 2 weeks under 12-h/12-h light–dark cycles before euthanasia at zeitgeber times (ZT): 0, 3, …, 21 h, in accordance with the regional animal ethics committee guidelines. Duplicates of pooled samples consisting of three biological replicates were acquired per zeitgeber time and group, respectively.

#### Human clinical sample acquisition

Patients included in this study were seen in the Department of Nephrology at the University of Cologne for their first manifestation of the nephrotic syndrome. Kidney biopsies were carried out as part of routine patient care. After diagnosis with either MCD or FSGS and after providing informed consent, patients were enrolled in the ForMe registry (ClinicalTrials.gov identifier NCT039499728)^53^ intended for the study of MCD and FSGS patients. Within this registry, we provide comprehensive genetic analysis, monitor the clinical outcome, and carry out in vitro and in vivo analysis of biosamples and novel mutations to predict and investigate MCD and FSGS outcomes. The registry itself and its proceedings were approved by the local ethics committee of the University Clinic Cologne, Germany.

Core patient clinical data at the time of kidney biopsy have been made available in de-identified fashion together with human spatial transcriptomics data (see Resource availability). Information on gender, ancestry, race, and socioeconomic status were not acquired within the registry and are hence not reported here.

### Method details

#### Isolation of glomeruli and nuclei

Mice were sacrificed by cervical dislocation and kidneys were extracted and perfused *ex vivo* through the renal arteries with magnetic M-450 Tosylactivated Dynabeads (14013, Thermo Fisher) suspension in HBSS (5.4 mM KCl, 0.3 mM Na_2_HPO_4_, 0.4 mM KH_2_PO_4_, 4.2 mM NaHCO_3_, 137 mM NaCl, 5.6 mM D-glucose, 1.3 mM CaCl_2_, 0.5 mM MgCl_2_, 0.6 mM MgSO_4_). Perfused kidneys were minced by scalpel and digested with digestion buffer (1 mg/ml Pronase E (P6911, Sigma), 1 mg/ml collagenase type II (LS004176, Worthington) and 50 U/ml Dnase I (A3778,0050, Applichem), in HBSS) for 15 minutes at 37 ℃. The kidney suspension was triturated until homogenous and filtered twice through 100 µm cell strainers (83.3945.100, Sarstedt). The kidney lysates were washed with HBSS and pelleted at 1500 rpm for 5 minutes at 4 ℃. The pellet was resuspended in HBSS and washed on the magnet until isolated glomeruli were obtained.

Isolated glomeruli were resuspended in 1 ml of lysis buffer (EZ lysis buffer (NUC101-1KT, Sigma) supplemented with EDTA-free protease inhibitors (5056489001, Roche)) and shaken at 1400 rpm for 10 minutes at 4 ℃. After magnetization, the supernatant containing loose nuclei was collected through a 30 µm cell strainer (04-004-2326, Sysmex). The leftover beads were resuspended in 1 ml lysis buffer further lysed 5 times through a 27-G needle. After the loose nuclei were collected, the beads were resuspended in 1 ml of lysis buffer and further lysed through a 30-G needle 5 or 10 times to collect the final aliquots of loose glomerular nuclei. The pooled nuclei were centrifuged at 500 g for 5 minutes at 4 ℃. The pellet was resuspended in 300 µl of lysis buffer, transferred to a 1.5 ml tube and placed on the magnet for 3 minutes to remove residual magnetic beads.

#### ATAC-seq

50,000 nuclei were isolated from glomeruli of nTnG;Pod:cre mice, stained with DAPI (D9542, Sigma) and sorted using the FACSAria III (BD) equipped with the 70 µm nozzle with chillers. Singlet and DAPI+ nuclei were gated by forward scatter and side scatter and the UV1 laser. DAPI+ nuclei were then sorted by fluorescent signal and collected into lysis buffer. Nuclei were centrifuged at 1000 g for 10 minutes at 4 ℃ and resuspended in 50 µl of transposition reaction mix (25 µl of transposition buffer, 2.5 µl of TDE1 (20034197, Illumina) and 22.5 µl of nuclease-free water). The reaction was carried out at 37 ℃ for 30 minutes. Immediately following transposition, DNA was purified using the DNA clean concentrator-5 kit (D4014, Zymo Research) per manufacturer’s instructions. Library preparation and sequencing were performed by the Cologne Centre of Genomics.

#### Single nuclei RNA-seq

Glomeruli were isolated from murine kidneys and subjected to isolation of nuclei using lysis buffer supplemented with 0.1% murine RNase inhibitor (M0314L, New England Biolabs GmbH). Isolated nuclei were counted with a hemocytometer. After centrifugation at 1000 g for 10 minutes, nuclei were resuspended in PBS supplemented with 2% BSA (3737.2, Carl Roth) to reach an approximate concentration of 1000 nuclei per µl. Prior to immediate processing, the samples were further filtered through a 10 µm strainer (04-004-2324, Sysmex). For Wt1^ko/wt^ and Nphs2^mut^ mice, glomeruli nuclei were used for snRNA-seq. For Pdss2^mut^ mice, glomeruli nuclei were combined with nuclei extracted from whole kidney (1:1 volume) to additionally capture tubules. Libraries and sequencings were prepared per the manufacturer’s instructions supported by 10X Genomics by the Cologne Centre of Genomics.

#### Bulk RNA-seq

To analyze circadian gene expression in kidneys, whole kidneys were dissected, snap-frozen and RNA was prepared using the miRNeasy mini Kit (217004, Qiagen) to isolate total RNA. After removal of ribosomal RNA using biotinylated target-specific oligos combined with Ribo-Zero ribosomal RNA removal beads, RNA was fragmented into small pieces and copied into first-strand complementary DNA (cDNA) followed by second-strand cDNA synthesis. Products were purified and enriched with PCR to create the final cDNA library. After library validation and quantification using the 2100 Bioanalyzer (Agilent), equimolar amounts of five to six pooled libraries were quantified by using the KAPA Library Quantification Kit (KK4856, Peqlab) and the Applied Biosystems 7900HT Sequence Detection System and sequenced on a Hiseq2000 sequencer.

#### Xenium spatial transcriptomics

Xenium Prime spatial transcriptomics were carried out using the 5K human pan tissues and pathways panel supplemented with a custom 50 gene panel covering PDS genes. For generation of the custom panel, the top 70 ranking genes were provided to the 10X custom panel design tool. Genes predicted to result in optical crowding due to high expression levels and genes predicted to cause technical problems in probe design as by 10X recommendations were excluded from the panel. The remaining top 50 PDS marker genes were included in the final custom panel. In case of genes overlapping between the 5K human panel and the PDS marker gene set, the number of probes per gene was boosted to improve the PDS signal. Full information on the custom panel is available with the Xenium spatial transcriptomics raw data (see section on Resource availability).

Kidney biopsy cylinders of four patient biopsies were combined in one FFPE tissue block, sectioned at 5 µm thickness and mounted according to 10X instructions and processed for Xenium spatial transcriptomics according to the manufacturer’s instructions. Standard H&E staining was carried out upon completion of spatial transcriptomic data acquisition. Biopsy cylinder attribution to individual patients is ensured by metadata and H&E images uploaded together with Xenium raw data (see section on Resource availability). Serial sections were obtained in the same manner for confocal microscopy (see “Immunofluorescence and quantification”).

Standard built-in segmentation analysis of the Xenium Prime platform was used for single podocyte segmentation. Images were generated using the Xenium Explorer 3 software (version 3.2.0).

#### STED microscopy and quantification

Murine kidney tissues were fixed in formalin overnight at 4 ℃ and stored in PBS. Kidney cortex was cut into 1-2 mm pieces and immersed in hydrogel solution (4 % acrylamide (Carl Roth), 0.25 % VA-044 initiator (Sigma) in PBS) overnight at 4 ℃. The gel was polymerized by incubation at 50 ℃ for 3 hours. Excess polymer was removed, and tissues were washed in PBS three times and embedded in 3% agarose (Carl Roth) prior to cutting into 200 µm sections with a vibratome (Leica). Samples were transferred to a clearing solution (200 mM boric acid, 4 % SDS, pH 8.5) and incubated at 50 ℃ until cleared. Samples were washed briefly in a wash buffer (10 mM HEPES, 200 mM NaCl, 3 % Triton X-100) before incubation with primary antibodies (nephrin: 20R-NP002, Fitzgerald; THSD7A: HPA000923, Sigma; Claudin-5: 35-2500, Thermo Fisher Scientific; EphB1: PAB3018, Abnova) at 1:100 at 37 ℃ overnight. Samples were washed with wash buffer before incubation with secondary antibodies (goat anti-guinea pig: 106-005-003, Jackson ImmunoResearch; donkey anti-rabbit: 711-005-152, Jackson ImmunoResearch; goat anti-mouse: 115-005-003, Jackson ImmunoResearch; each conjugated to fluorophores in-house) at 1:100 at 37 ℃ overnight and embedded in saturated fructose solution (80.2 % w/w fructose (USBIF8000, VWR), 0.5 % 1-thioglycerol (M6145, Sigma)) before mounting in a glass-bottom dish (Ibidi) for imaging with the SP8 STED microscope (Leica).

To quantify the distribution of Claudin 5 at the podocyte slit diaphragm, the fluorescent intensity of claudin 5 along a manually traced slit was quantified in ImageJ (version 1.54),^122^ the area of under the curve was calculated and normalized to slit diaphragm length.

#### Albumin to creatinine ratio quantification

Urines from mice were analyzed to determine albumin and creatinine ratio (ACR). Urinary albumin was determined by ELISA. Briefly, wells were coated for 1 hour with 100 µl of anti-mouse albumin coating antibody (A90-134A, Bethyl Laboratories) at 1:10,000 in coating buffer (0.05 M carbonate-bicarbonate, pH 9.6), then washed 5 times with 200 µl of wash solution (50 mM Tris, 0.14 M NaCl, 0.05 % Tween-20, pH 8.0) before incubating for 30 minutes with 200 µl of blocking solution (50 mM Tris, 0.14 M NaCl, 1 % BSA, pH 8.0). The wells were washed again 5 times with 200 µl of wash solution before incubation for 1 hour with 100 µl of standards or samples diluted in diluent buffer (50 mM Tris, 0.14 M NaCl, 0.05 % w/v Tween-20, 1 % w/v BSA, pH 8.0). Next, the wells were washed 5 times with 200 µl of wash solution before incubation with 100 µl of HRP detection antibody (A90-134P, Bethyl Laboratories) diluted 1:25,000 in the diluent buffer for 1 hour. Finally, the wells were washed 5 times with 200 µl of wash solution and developed with 100 µl of substrate solution (100 ug/ml 3,3’,5,5’-Tetramethylbenzidine (TMB), 48 mM sodium acetate, 0.01 % v/v hydrogen peroxide, pH 5.2) for 15 minutes in the dark. The reaction was stopped by adding 100 µl of stop solution (0.18M sulfuric acid) and the absorbance was measured at 450 nm. Urinary creatinine was measured using a commercial colorimetric kit (500701, Cayman Chemical). Briefly, urines were diluted 1:20 in water and assayed according to the manufacturer’s protocol. All samples were measured in triplicates. To obtain the urinary ACR, albumin concentrations were divided by creatinine concentrations.

#### Immunofluorescence and quantification

Human kidney biopsies were sectioned into 5 µm thickness and mounted on slides for confocal imaging as described in Unnersjö-Jess et al.^123^ Briefly, after deparaffinization antigen retrieval was carried out with heated Tris-EDTA at pH 9. Slides were washed in PBS supplemented with Triton-X 0.5 % and incubated with primary antibody (nephrin: AF4269, R&D Systems) in PBS supplemented with 5 % bovine serum albumin and 5 % normal donkey serum overnight. Slides were washed in PBS and incubated with Alexa 488-coupled donkey anti-goat secondary antibody (A11055, Thermo Fisher Scientific), washed in PBS supplemented with DAPI and mounted with 80% fructose solution with 0.5 % 1-thioglycerol. Slides were imaged on a SP8 3x confocal microscope (Leica) using a 100x lens. Acquired images were deconvoluted using Huygen’s software (version 25.04). The slit diaphragm length per area was measured using the nephrin staining using ImageJ (version 1.54)^122^ and published macro as described in Butt et al.^46^

Formalin-fixed paraffin-embedded (FFPE) murine kidney samples were sectioned to 5 µm using a microtome and treated by heat-mediated antigen retrieval in Tris-EDTA pH 8. The sections were permeabilized using PBS-T (0.1 % Triton X-100) and blocked for 1 hour at room temperature using normal donkey serum (017-000-121, Jackson ImmunoResearch) before overnight incubation with primary antibodies (Nr1d1: 13418, Cell signaling Technology, Wt1: AF5729, R&D Systems, PKM: 25659-1-AP, Proteintech, phosphorylated-S6: 4858, Cell Signaling Technology). Samples were washed and stained with secondary antibodies and Hoechst dye before mounting in ProLong Gold Antifade Mountant (P36934, Invitrogen). Images were acquired using the Zeiss Apotome 3 microscope. Podocyte nuclei were segmented and the expression of Nr1d1 was quantified using QuPath (version 0.6.0),^124^ using the InstanSeg extension (version 0.1.0-rc1) and the fluorescence_nuclei_and_cells model.^125^ Nphs2^mut^ mice aged 8 weeks and Pdds2^mut^ mice aged 21 weeks were used for phospho-S6-kinase and Pkm staining, as these represented similar damage levels as by SD length and albuminuria. For Pkm staining kidneys were sectioned to 2 µm and heat-mediated antigen retrieval was carried out using citrate buffer.

FFPE murine livers were sectioned to 7 µm using a microtome, deparaffinized, and rehydrated prior to heat-mediated antigen retrieval in citrate buffer pH 6. Sections were treated for 10 min with 0.3 % glycine in PBS, washed with PBS, and permeabilized using 0.03 % SDS in PBS. Sections were blocked for 1 hour with 3 % normal donkey serum in PBS-T (0.25 % Triton X-100). Primary antibody (CYP7A1: PA5-100892, Thermo Fisher Scientific, 1:100) was applied overnight at 4 °C. Slides were washed with PBS-T (0.1 % Triton X-100) and stained with secondary antibodies and Hoechst 33342 (H1399, Invitrogen, 1:1000) diluted in PBS-T (0.25 % Triton X-100) at RT. After a final wash, slides were mounted using Vectashield Antifade Mounting Medium (Vector Laboratories). Images were acquired with an Olympus Slideview VS200 system (Olympus). For CYP7A1 quantification, regions of interest (ROIs) around bile ducts were preselected in 16-bit .vsi images using QuPath (version 0.6.0). ROIs were further processed using Fiji (version 2.16.0) by measuring the mean intensity of CYP7A1-positive areas. The average CYP7A1 mean intensity per mouse liver is represented.

#### qRT-PCR

Frozen liver tissue was homogenized in QIAzol (79306, Qiagen), followed by total RNA isolation using the RNeasy Mini Kit (Qiagen, 74106) with DNase treatment using the RNase-free DNase Set (79254, Qiagen) according to the manufacturer’s instructions. Reverse transcription was performed using the High-Capacity cDNA Reverse Transcription Kit (4368813, Applied Biosystems) according to the manufacturer’s protocol. Subsequently, qPCR was conducted using the QuantStudio 7 Flex Real-Time PCR System with Taqman Real-Time PCR Assays (*Cyp7a1*: Mm00484150_m1, *Cyp27a1*: Mm00470430_m1, *Tbp*: Mm00446973_m1, Thermo Fisher Scientific) and Takyon Low ROX Probe 2x MasterMix dTTP Blue (UF-LPMT-B0701, Eurogentec). Relative target mRNA expression was normalized to expression of *TATA-binding protein* (*Tbp*) and analyzed using the 2^-ΔΔCt^ method. Expression levels are presented relative to the control condition.

#### Senescence-associated β-gal staining

Livers were removed from mice and cryopreserved in Tissue freezing medium (14020108926, Leica). 15 µm cryosections were generated using the Leica CM3050 S cryostat. Sections were fixed in 0.2% glutaraldehyde (50-262-23, Electron Microscopy Sciences) in PBS for 10 min at room temperature. Sections were washed for 3 x 1 min using PBS at RT and incubated in freshly prepared X-gal staining solution (40 mM citrate sodium dihydrogen phosphate buffer pH 5.5, 5 mM potassium hexacyanoferrate(II) trihydrate, 5 mM potassium ferricyanide(III), 150 mM sodium chloride, 2 mM magnesium chloride hexahydrate, and 1 mg/ml 5-bromo-4-chloro-3-indolyl-b-D-galactopyranoside (X-gal)) for 24 h at 37 °C. Finally, slides were washed 3 x 1 min with PBS, dehydrated, and embedded using Cytoseal XYL medium (Thermo Fisher Scientific). Images were acquired with an Olympus Slideview VS200 system (Olympus).

#### Transcriptomic data retrieval and processing

CEL files were downloaded for microarray data. UMI count tables were downloaded for single cell RNA-seq experiments. Unnormalized gene expression count matrices were used, when available, in case of bulk RNA sequencing experiments. If the data was not available in this format, unnormalized count matrices were generated from raw reads, as described below.

Gene IDs were mapped to Mouse Genome Informatics (MGI) gene symbols as a common reference using the R package biomaRt (version 2.50.2).^126,127^ On a few occasions, if the expression data set included counts for different splice variants of the same gene, the average expression across the splice variants was taken as the expression count of that gene before converting to MGI nomenclature, since MGI nomenclature does not account for alternative splicing.

##### RNA-seq data processing

Raw sequencing files of published datasets available in the ‘sra’ format were extracted and converted to ‘FASTQ’ format using the SRA Toolkit (version 2.11.3). Quality control of FASTQ files was done with FastQC (version v0.11.9).^128^ MultiQC (version v1.1) was used to summarize the quality statistics of multiple samples from one study into a comprehensive overview.^129^ No samples were excluded due to lack of quality.

An index of the *Mus musculus GRCm38 (mm10) and* GRCm39 reference genomes and its annotations were generated using STAR (version 2.7.3) aligner to be used for alignment of raw reads from liver and kidney studies,^130^ correspondingly. When STAR was run to align reads, the option -quantMode’ was set to the command ‘GeneCounts’ to generate count matrices as an output. Gene abundances were quantified using both exonic and intronic reads with pre-mRNA full gene models.

##### Sn- and sc-RNA-seq data processing

STARsolo function of the STAR aligner was used to map reads of single nuclei RNA-seq experiments. Sequence files from 2 sequencing lanes, if present, were pulled in one sample. Ambient RNA was removed from UMI count tables using decontX (version 1.2.0) R package.^131^

The read count (UMI) matrices of all single cell RNA-seq samples were processed in Seurat (version 4.0.1) R package.^132^ As individual datasets stem from multiple sources data was handled individually with respect to quality control, filtering, dimensionality reduction, and normalization. The respective parameters have been included in out GitHub repository (see Section on Code availability). Doublets were removed in snRNA-seq datasets with scDblFinder (version 1.13.6) using cluster-based approach.^133^

#### ATAC-seq data processing

Mapping to the mouse genome mm10 was done using bwa-mem (version 0.7.17) with default parameters.^134^ Data quality was assessed by calculating the insert sizes between the R1 and R2 read pairs. Peaks were called using Genrich (version 0.6.1) tool using default parameters.^135^

#### Spatial transcriptomics data analysis

##### Mouse Visium kidney spatial transcriptomics

Kidney Slide-seq V2 spatial transcriptomics data was retrieved from GSE190094.^45^ Podocyte annotation and KNN filtration, glomerular identification and size estimation were performed using the original code from the publication.^45^ Samples with low capture rates (below the average of 100 features per bead) and without visually detectable podocyte clusters were removed, which excluded 20 out of 49 original samples from the downstream analysis. For the remaining 29 samples, the number of beads annotated as podocytes was normalized to the glomerular area. Subsequently, the bead density per glomerulus was correlated with the average PDS of the respective glomerulus.

##### Mouse Visium liver spatial transcriptomics

HDS was applied on spatial transcriptomic data from the Liver Cell Atlas dataset.^9,48^ Seurat (version 4.0.1) was used to analyze the 10X Visium counts.^132^ Uninformative spots located in regions with no or minimal tissue were removed from analysis. Filtering for sequencing quality control was performed using the same parameters as described in the Liver Cell Atlas GitHub repository accessible at https://github.com/guilliottslab/scripts_GuilliamsEtAll_Cell2022. Spots were retained only if they contained more than 1000 detected features (nFeature), total number of counts (nCount) between 5000 and 60 000, and less than 15% mitochondrial RNA. The HDS was computed per spot.

##### Human kidney spatial transcriptomics

10X Visium spatial transcriptomic data from a human with FSGS, patient ID 29-10282, was retrieved from KPMP and analyzed.^3^ The count data and the spots coordinates were loaded and processed with Seurat (version 4.0.1).^132^ Glomeruli were annotated manually. The PDS was calculated per spot for all spots within annotated glomeruli using the PDS_calc function (see Github repository – section Code availability) with standard parameters.

Human Xenium data was analyzed with the Seurat package. Slide scans were split in individual samples with sf (version 1.0-20)^136^ R package by drawing a polygon around each sample and then cropping all layers of the Seurat object to the selected region. Default cellular segmentation was used for the analysis. Cell types were annotated with Azimuth (version 0.5.0)^132^ R package, using the default set of cell-type markers.

For automated glomerular size estimation, coordinates of podocytes, endothelial and mesangial cells were used as input to KNN filtration and clustering (see Method section Mouse kidney spatial transcriptomics), to identify clusters of glomerular cells and filter out scattered individual cells. Glomeruli were segmented by drawing circles around centers of clusters, using distance from the cluster center to the furthest cell of the cluster as glomerular radius. Results were inspected manually and nested glomeruli or clusters containing more than one glomerulus were excluded from downstream analysis. PDS was calculated for all individual podocytes within automatically segmented gloms as detailed for Visium analysis.

##### Human liver spatial transcriptomics

Human spatial transcriptomic data from patients “H35” and “H37” were retrieved from the Liver Cell Atlas^9,48^ and Seurat was used to analyze 10x Visium counts. Spots representing hepatocytes were manually annotated and classified a steatotic or non-steatotic based on histology. The count matrices were subset to include only protein coding genes and only the spots annotated as hepatocytes with more than 250 detected features were included in the analysis (nFeature_Spatial > 250). We used the Seurat function NormalizeData() to normalize the counts per spot in each sample and proceeded to calculate the HDS.

#### Proteomics data analysis

Proteomics datasets PXD016238,^13^ PXD018326,^46^ PXD047417,^50^ and PXD012063^51^ were retrieved from proteome exchange. The LFQ data was processed with the DEP (version 1.4.1).^137^ Only proteins expressed in more than 50% of the replicates in at least one condition were considered. To calculate PDS in proteomics data, 62 instead of 42 podocyte damage markers were used to ensure at least half of the markers are present in all the studies. For the application of the HDS on murine proteomic data, processed data from Williams et al. provided by the authors were used.^52^

#### Differential gene expression analysis

CEL files for microarray data were processed and differential expression analysis performed using the oligo (version 1.54.1)^138^ R package with the default parameters. Non-normalized bulk RNA-seq count tables were subjected to differential expression analysis with DEseq2 (version 1.34.0)^139^, with default parameters. For hepatocyte data, counts were transformed using the variance stabilizing transformation in DESeq2 with the parameter ‘blind’ set to ‘true’. Single cell RNAseq UMI count tables were processed and differential expression analysis performed with Seurat^108^ using the FindMarkers() function with the default parameters. For single cell data, only cells of the respective cell type of interest were considered (i.e. podocytes or hepatocytes). Gene set over-representation analysis was performed using ClusterProfiler (version 4.10.1).^140^ For details on specific parameters, see our GitHub repository.

#### Generation of PDS and HDS marker genes

Candidate lists for damage markers were generated to ensure that marker genes were expressed in the targeted cell type (here: podocyte for PDS, hepatocyte for HDS). Data from diseased and control samples were included. The expression of genes included in the damage scores was not required to be exclusive to the targeted cell type, allowing for the inclusion of marker genes that are generically expressed in various cell types.

To create a catalogue of genes expressed in podocytes, three single-nuclei RNA-seq kidney datasets generated within this study, and four published single-cell RNA-seq kidney datasets from GSE146912^55^ (Supplementary Table 1) were used, with each of the datasets representing a unique podocyte damage model. Cells annotated as podocytes were subset from all seven datasets and used for the selection.

To create a catalogue of genes expressed in hepatocytes, murine gene expression data from the Liver Cell Atlas^9,48^ were used and analyzed using Seurat^132^. Cells annotated as hepatocytes by the authors were subsetted from the two experiments available: Standard Diet and Non-Alcoholic-Fatty-Liver-Disease (NAFLD).

The selection procedure for the identification of genes expressed in podocytes and hepatocytes single-cell RNA-seq data was identical. A detection rate for each gene was calculated across cells that fulfilled the following criteria: ‘nFeature_RNA’ greater than 500 and lower or equal to 6,000, less or equal to 2.5 % mitochondrial RNA and ‘nCount_RNA’ smaller than 30,000. Genes with a detection rate of at least 0.03 (i.e. 3% of the cells) in the target cell type were kept for the subsequent analyses.

#### Damage score algorithm

To generate the damage scores, we selected and analyzed datasets that represent different disease models and that were partially obtained using different experimental platforms (Table S1), as well as novel data generated for the PDS (see above and Table S1). First, each dataset was individually processed and analyzed to obtain differentially expressed genes, always comparing experimental condition versus control. In case of the single cell data, only cells of the respective cell type were considered (i.e. podocytes or hepatocytes). Results of differential expression analysis were integrated across studies in the following way:

1. Gene must be identified as potentially expressed in podocytes or hepatocytes respectively.
2. Genes were selected if they were detected in >=75% of the studies.
3. Out of those, genes were kept if their fold changes were in the same direction (i.e. either positive or negative fold change) in >=75% of the studies.
4. Selected genes were ranked by their p-value of differential expression per study. P-values for differential expression may not be comparable between studies, due to variation in sample numbers and/or variable quality of the data. We therefore decided to base the selection process on ranks of p-values rather than defining a common p-value threshold across studies.
5. Ranks were combined across studies by averaging the rank of each gene.
6. Genes were sorted by their mean rank, and the top N (N = 42 for the final score) were selected as a universal damage signature. In the calculation of the score, damage signature genes were treated differently depending on whether they were up or downregulated under the disease condition (see below).

Damage scores were computed for each cell/spot/sample using the damage signature genes with the AUCell (version 1.12.0)^12^ R package. AUCell scores were calculated separately for up and down regulated genes and combined as follows to get the final podocyte damage score:

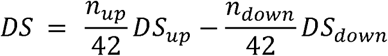

where *DS_up_* and *DS_down_*are the respective AUCell scores obtained with the up and downregulated genes. Note that the AUCell score is negative for low-ranking genes. These scores were weighted based on the number of up and down regulated genes (*n_up_*, *n_down_*). Since in both cases (PDS and HDS) the majority of marker genes were downregulated, *DS_down_* contributed more to the final damage score *DS*.

#### Damage score validation

##### Leave-one-model-out cross-validation

Cross validation was performed by excluding all data of one disease model group (Table S1) from the training procedure. Damage signature genes were determined as described above on the remaining training data. Subsequently, damage scores were computed as described above on the samples from disease model left out for testing.

##### Damage signature randomization test

Robustness of the PDS was tested by randomizing an increasing percentage of studies used to generate the PDS by means of shuffling p-values and LFCs of differential expression results. Signatures for partially randomized data were generated and used to calculate deviant damage scores in a test snRNA-seq dataset. As the choice of studies for randomization likely affects the resulting scores, 50 rounds of subsampling of studies were carried out for each level of randomness (i.e. percentage of randomized studies) and results were averaged separately for control and experimental conditions of the test dataset (nephrotoxic serum nephritis, GSE146912).^61^

##### Damage signature transfer to human data

Damage signature genes in humans were determined by manually mapping mouse genes to their human orthologs using the MGI gene annotation.^141^ Signature genes without known human orthologs were omitted. In cases of several human orthologs to a single mouse gene, we kept all human orthologs. In cases of several mouse disease signature genes with a single human ortholog, we included the human ortholog and reduced the number of genes contributing to score calculation.

##### Pseudo-time trajectory inference

After filtering hepatocytes based on annotation and quality control metrics using Seurat, we performed pseudo-time trajectory (PT) inference on normalized counts of each sample separately. We used TSCAN (version 1.13.0),^59^ and Monocle3 (version 1.3.7)^60^ with default parameters.

##### Damage score alignment test

We hypothesized that damage scores of cells from control samples were more similar to each other across studies, as opposed to damage scores from experimental conditions varying stronger between studies, as different interventions affect cellular damage to different degrees. To test our assumption statistically, we first calculated pairwise Kolmogorov-Smirnov (KS) distances between the damage score distributions of all control and experimental samples of all sn/scRNA-seq datasets and then used a one-sided two-sample Wilcoxon test to see if the average KS distance between damage score distributions of individual samples is lower in control than in experimental samples.

#### Disease association of damage signature genes

To find genomic variants associated with damage signature genes, we extracted the genomic coordinates of the RefSeq canonical isoforms for PDS and HDS marker gene loci and extended them to include 1) a 2 kb promoter; 2) the 5’ upstream DNA sequence spanning from the start of the promoter to the adjacent gene or 250 kb whichever is smaller; and 3) the 3’ downstream sequence spanning from the end of the 3’ UTR to the adjacent gene or 5 kb whichever is smaller. We then used the gwasrapidd (version 0.99.17)^142^ R package to search the NHGRI-EBI GWAS catalog for all variants associated with EFO traits related to kidney or liver disease, respectively. Search terms have been deposited in our GitHub repository (see Section on Code availability). Finally, we intersected selected variants with the extended genomic ranges of the podocyte and liver damage marker genes. To test the significance of enrichment of damage signatures in GWAS variants we used Fisher’s exact test, where all genes expressed in sc/snRNA-seq kidney and liver datasets were used as the corresponding backgrounds.

#### Pathway activity analysis

Pathway activity was defined as the AUCell (version 1.12.0)^12^ score obtained for the set of genes annotated as pathway members. Pathway membership (i.e. gene annotation) was taken from Reactome, KEGG and 50 hallmarks from MSigDB.^143^ This scheme was used to compute relative pathway activities per single cell and for bulk RNA-seq samples. To obtain pathway signatures of the damage models we computed Spearman correlation of single-cell pathway activities with the PDS for each damage model and for the cumulated controls. Per disease model, the top 10 significantly correlating pathways were included in further analyses (adjusted p-value < 0.05), resulting in a set of 85 pathways (Figure S8C). Of those 85 pathways, 26 pathways were manually selected for visualization in Figure 4A.

#### Transcription regulatory network construction

To identify transcription factors (TFs) expressed in podocytes, we utilized sc/snRNA-seq data from podocytes as follows: we selected TFs with a detection rate above 5% in at least one of the analyzed sc/snRNA-seq datasets, which resulted in 110 transcription factors contributing to the construction of the podocyte transcription regulatory network (TRN). A TRN is a directed graph representing interactions between transcription factors and their target genes. The TRN was constructed using the podocyte ATAC-seq data and known transcription factor binding motifs in following steps: First, ATAC-seq footprinting was performed on the ATAC-seq peaks with TOBIAS (version 0.14.0)^144^ to more precisely locate putative TF binding events. ^145^ Second, motif scanning was done on ATAC-seq footprints using FIMO from the MEME-suit (version 5.5.0).^146^ Motif hits with a p-value < 5 x 10^-4^ were kept for the analysis. Third, motif hits (i.e. potential TF binding sites) for each of the 110 TFs were associated with putative target genes using the ClosestGene approach implemented in the TFtargetCaller (version 0.7) R package.^147^ Genes with a q-value < 0.1 were assigned as targets of the corresponding TF.

TFs of a common family often have very similar or even identical position weight matrices (PWMs) resulting in virtually identical DNA binding motif predictions. To address this ambiguity, PWMs of 110 analyzed TFs were subjected to hierarchical clustering, resulting in 14 TF motif clusters with high similarity of motif sequences and predicted target genes within each TF family. We combined predicted targets of all TFs within each family into one regulon for the downstream analysis.

#### Circadian clock analysis

##### Kidney-specific circadian genes analysis

Bulk RNA-seq data from kidneys of young (10 weeks) and old (80 weeks) mice were used to identify kidney-specific circadian genes. Cycling genes were identified with the meta2d(cycMethod = ““JT””) function from MetaCycle (version 1.2.0).^148^ Overall, 317 genes with BH adjusted p-value < 0.01 in either young or old mice were called ‘circadian’. Estimation of acrophase and acrophase uncertainty for circadian genes was done by fitting a cosinor model with the cglmm(Y ∼ amp_acro(time_col=Time, period = 24)) function of GLMM cosinor (version 0.2.0.9)^149^ R package, where Y are the rlog-normalized expression values of a gene.

##### Circadian rhythm disruption analysis

Circadian rhythm disruption (CRD) in individual podocytes was calculated using cal_CRDscore() from the CRDscore (version 0.1.0) R package.^91^ The set of cycling genes identified from the kidney circadian bulk RNAseq was used to calculate CRD.

##### Tempo analysis

Circadian time in individual podocytes was estimated by the Tempo (version 0.0.1.dev)^90^ algorithm using expression of core clock genes. Estimates of gene acrophases, required by the algorithm, were taken from the analysis of bulk circadian RNA-seq data from the mouse kidney dataset. *Clock* was used as the reference gene.

#### Hepatocyte cell state analysis

AUCell was used to calculate the activity score of genes in the SHGS gene set.^94^ The HDS and SHGS activity scores were then used to classify hepatocytes in murine snRNA-seq data from Carlessi et al.^96^ into five cell state trajectories. The global HDS distribution across all sampleswas assessed to define HDS thresholds: undamaged - HDS smaller than the global 25%-quantile; transition - HDS between the global 25%- and 75%-quantile, damaged - HDS greater than the global 75% quantile. The distribution of the SHGS activity score across samples from the TAA condition was used to set senescence thresholds: senescent – greater than 75% quantile; unresolved – between 25 % and 75 % quantile, not senescent – lower than 25 % quantile.

To find genes differentially expressed in damaged senescent hepatocytes and not-senescent damaged hepatocytes, we used Seurat’s FindMarkers function on a SC-transformed merged Seurat object. By default, the function utilizes a Wilcoxon Rank Sum Test. FindMarkers performs p-value adjustment using Bonferroni correction based on the total number of genes expressed. We defined genes significantly differentially expressed if they had an adjusted p-value < 0.05, an absolute Log2 fold change > 0.5 and were expressed in at least 10 % of hepatocytes of either senescent-damaged or not-senescent damaged clusters.

Murine scRNA-seq hepatocyte data from Vesting et al.,^109^ was provided with quality control, normalization, and cell-type-annotation by the authors. HDS and SHGS scores, activities and distributions were calculated as described above. Damage and cell state classification was carried out in identical fashion.

### Quantification and statistical analysis

The statistical methods for each experiment were conducted using R scripts. Details regarding datasets were provided in the figure legends, while statistical information regarding experiments and analyses were available in both the figure legends and the main text.

## Key resources table

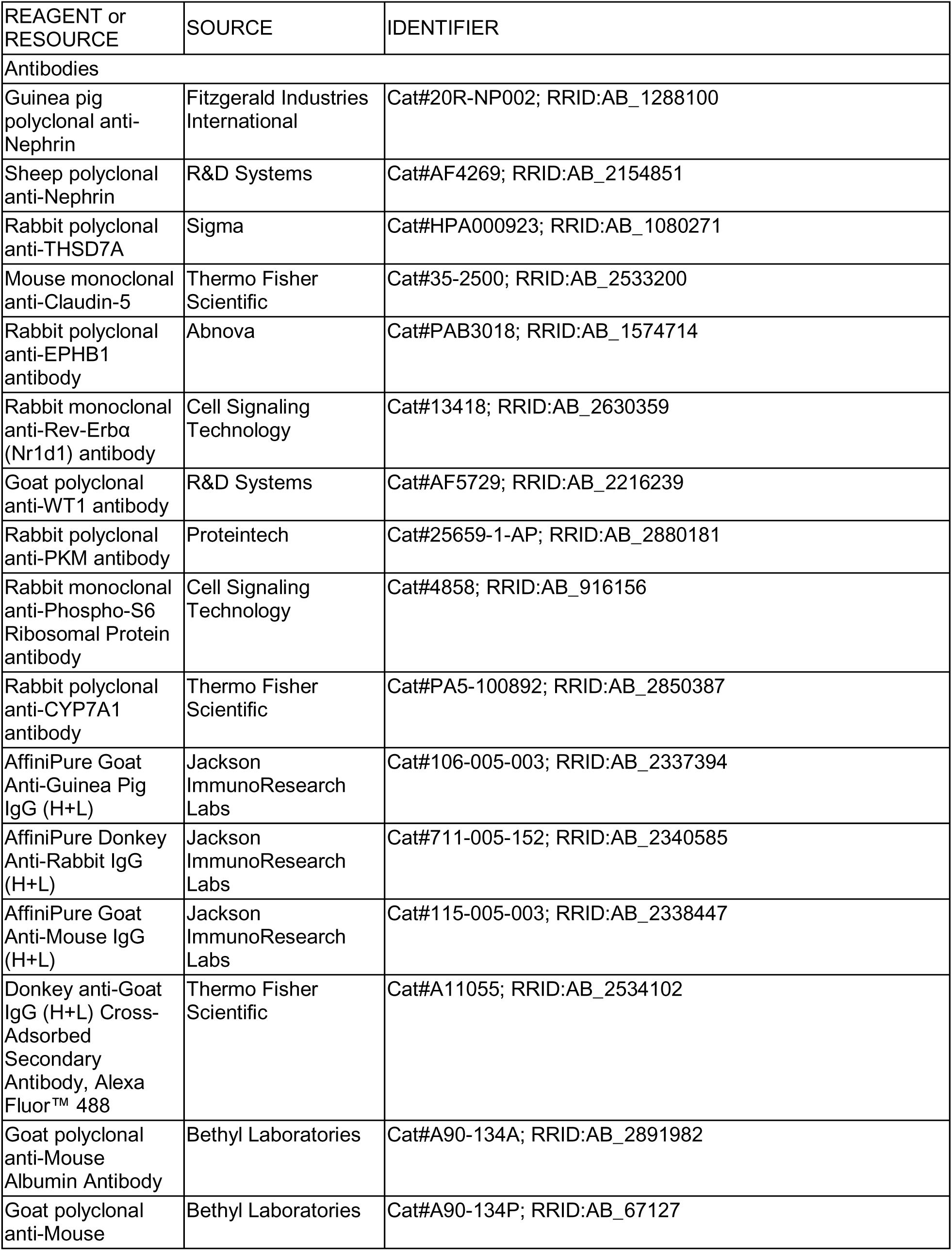
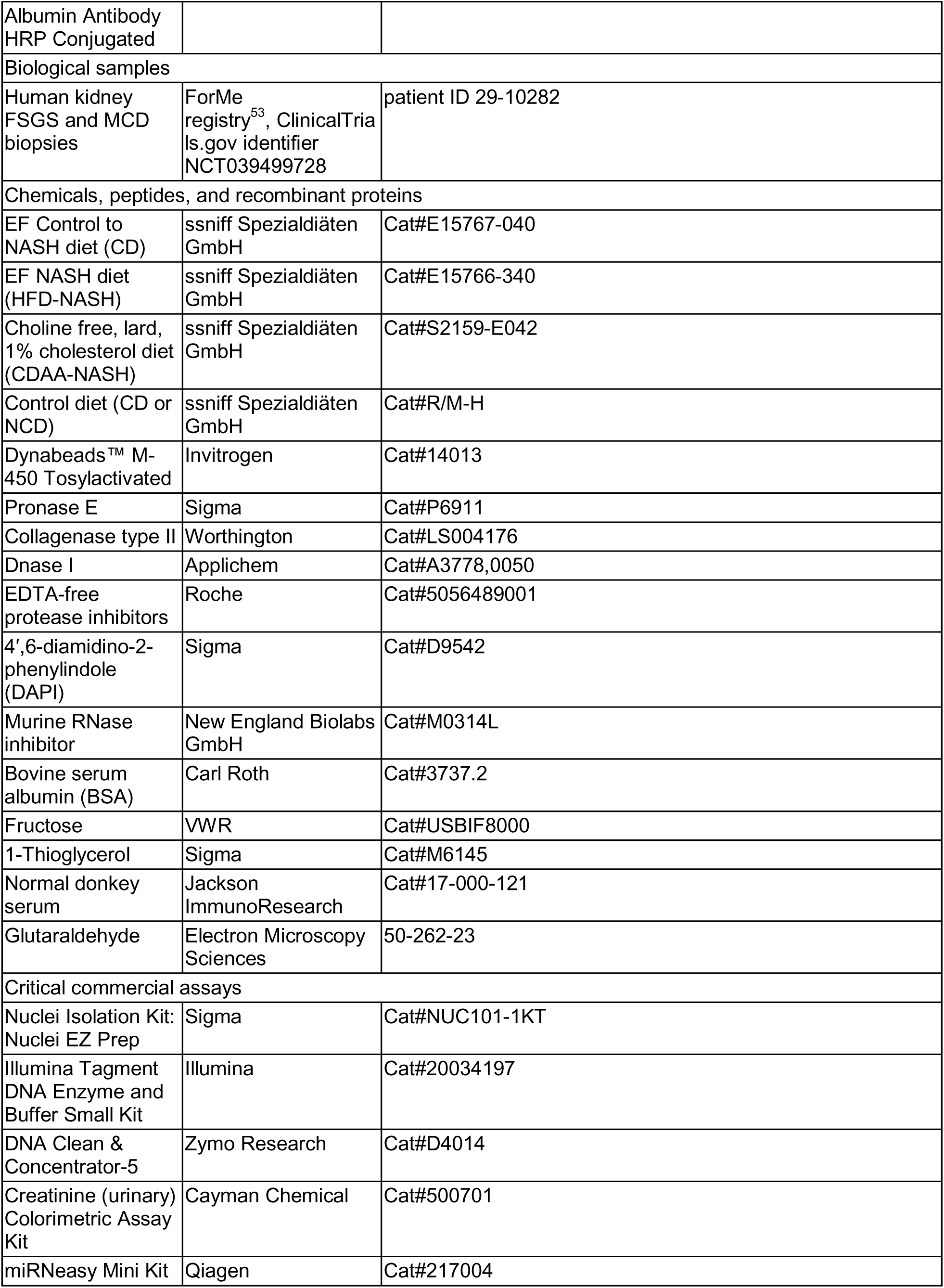
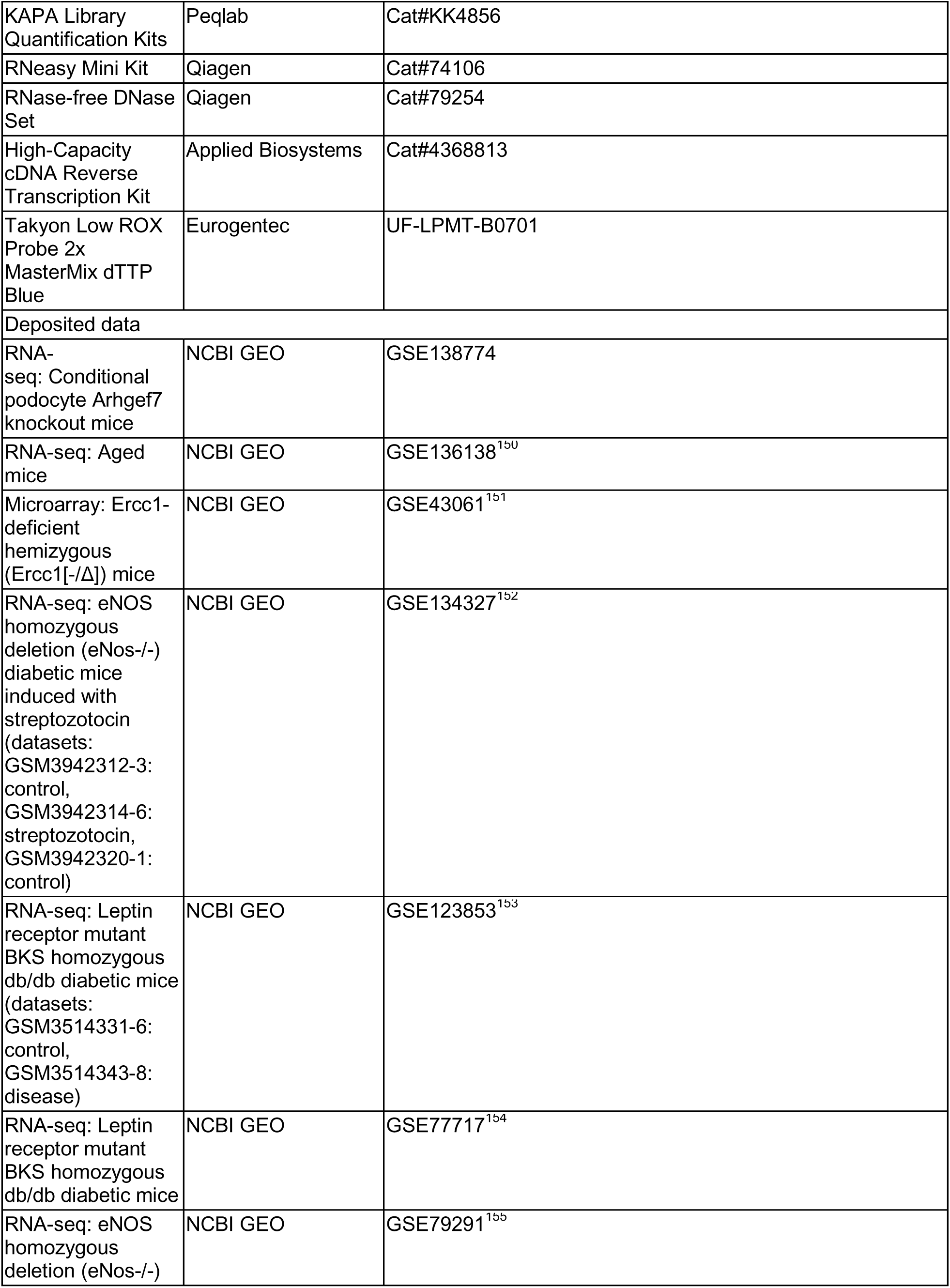
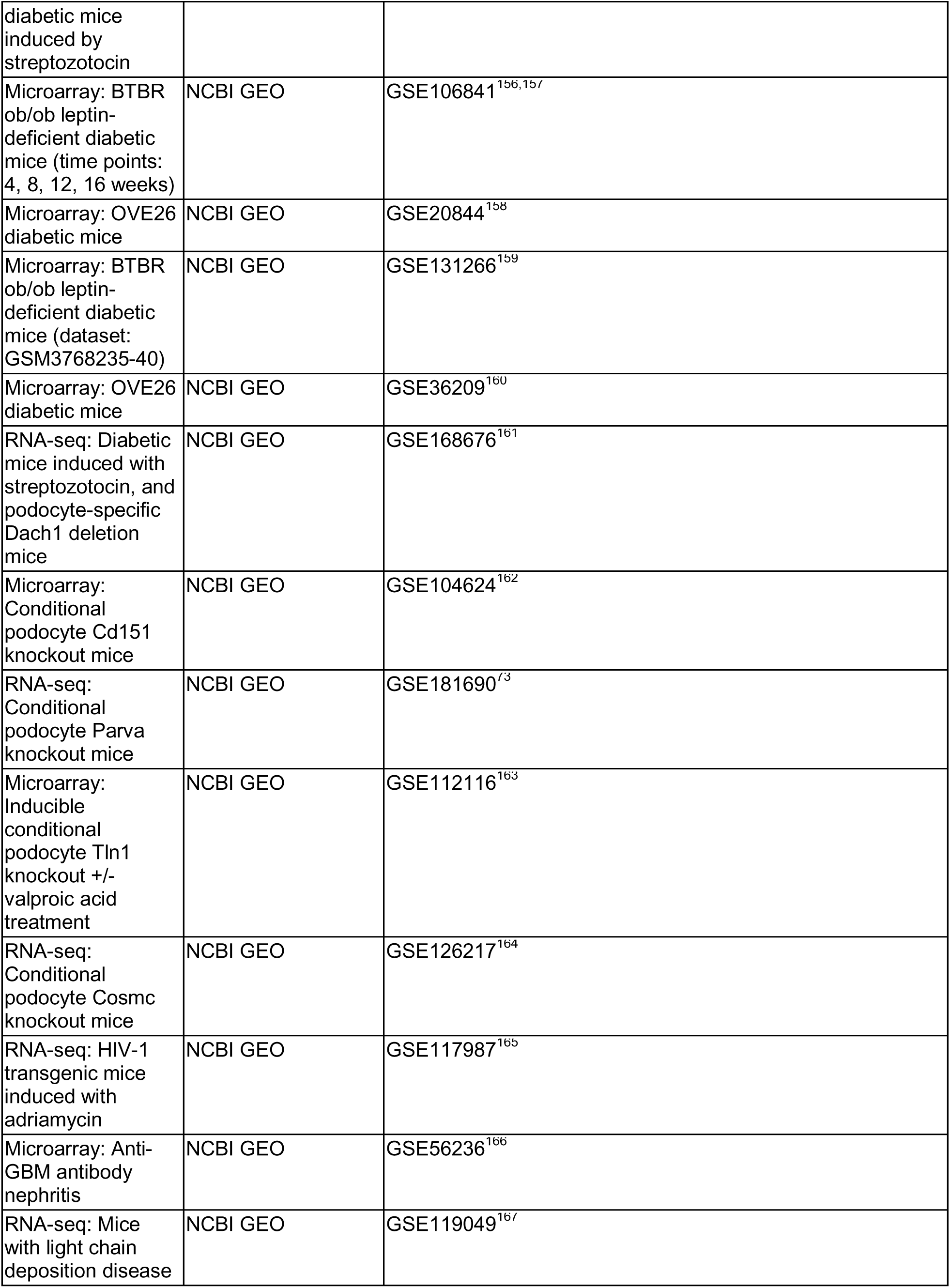
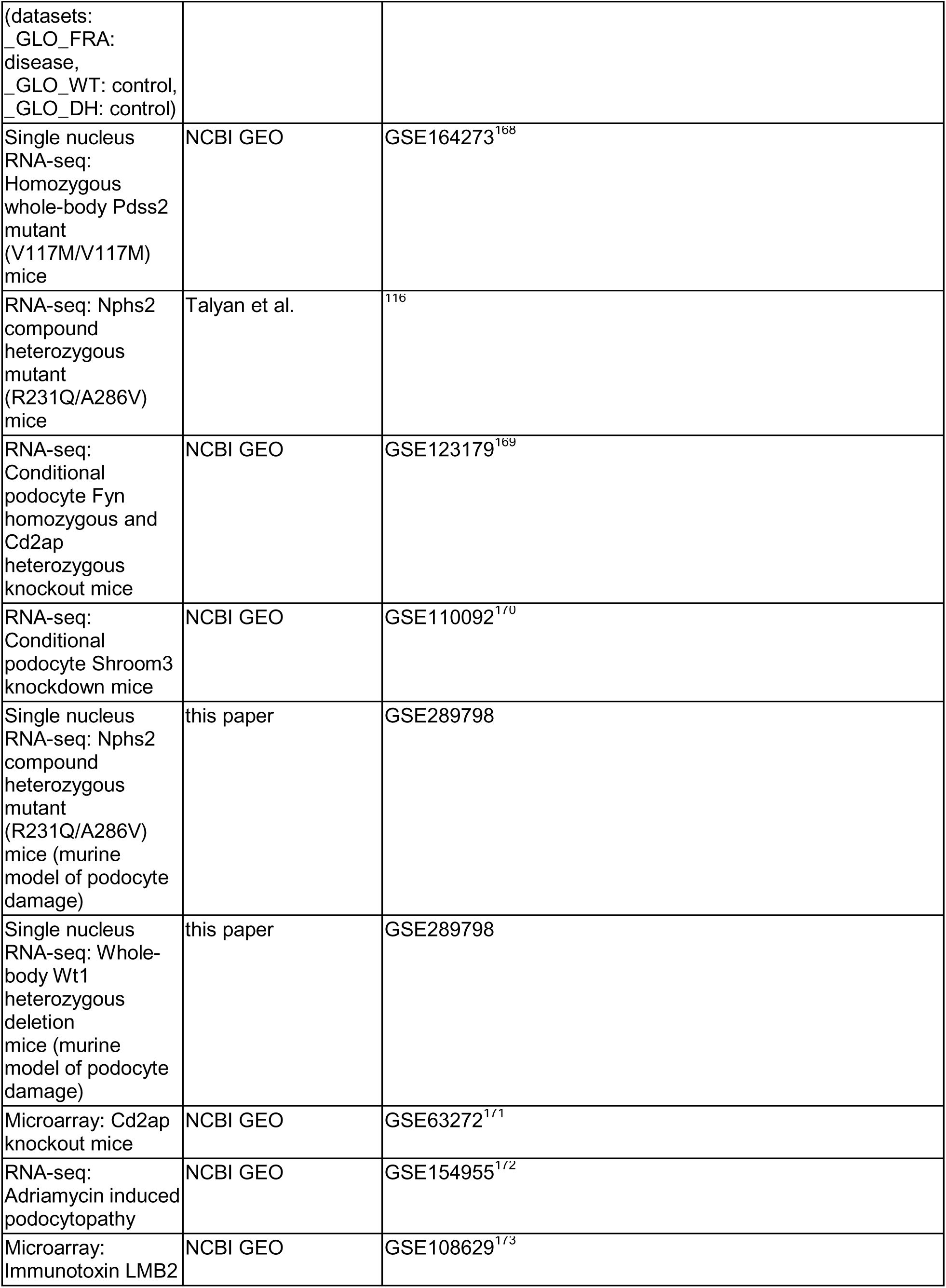
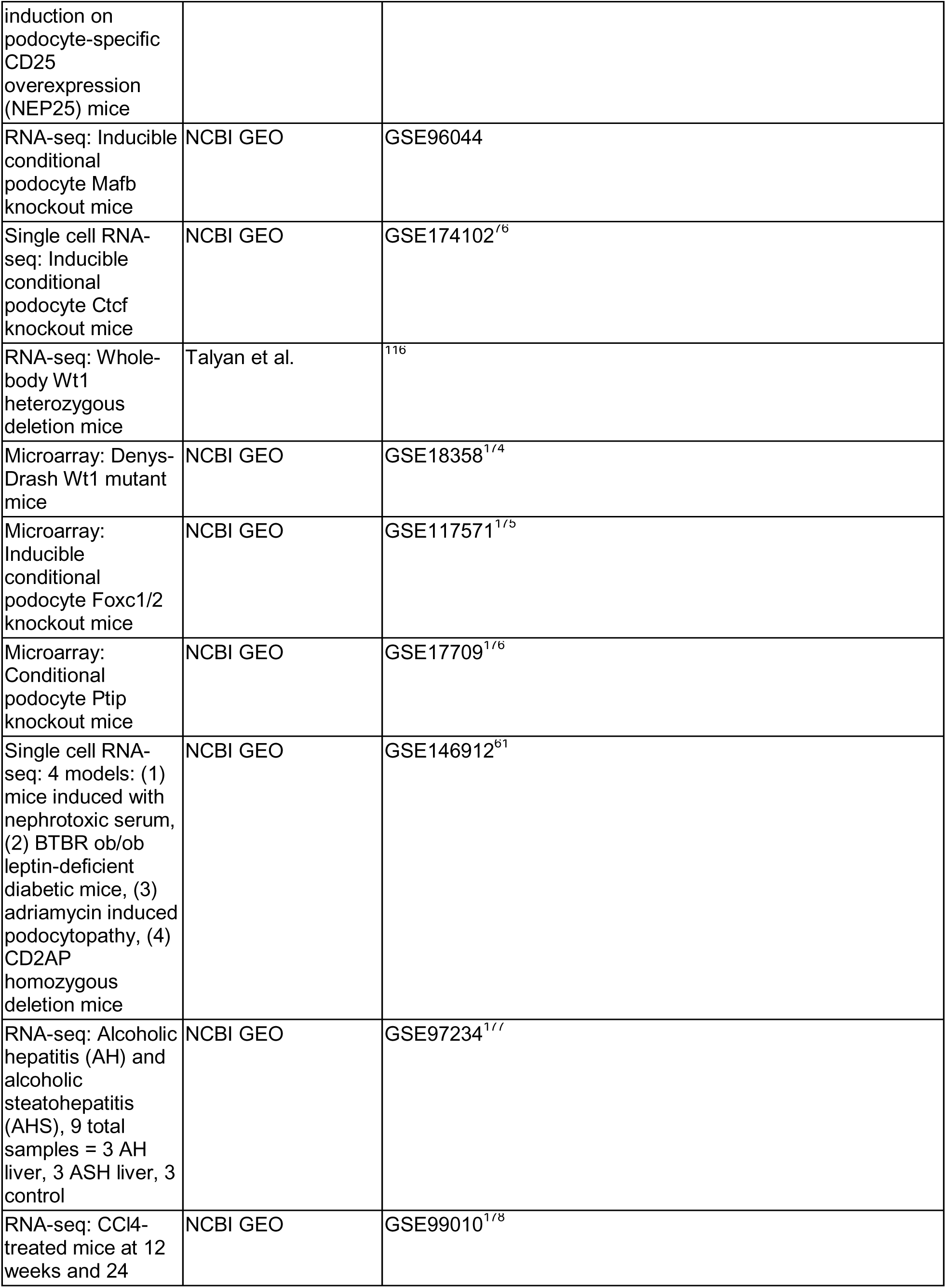
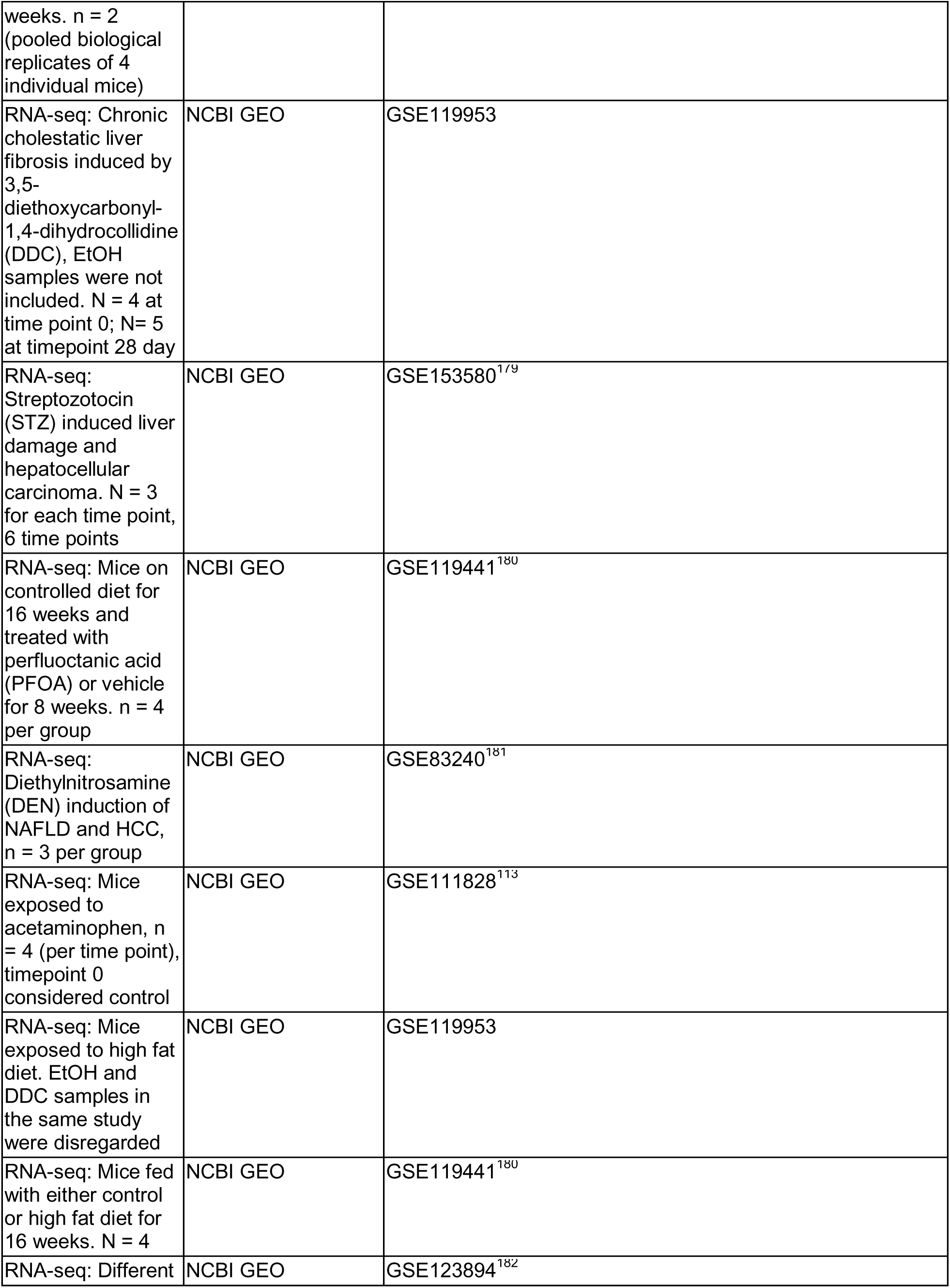
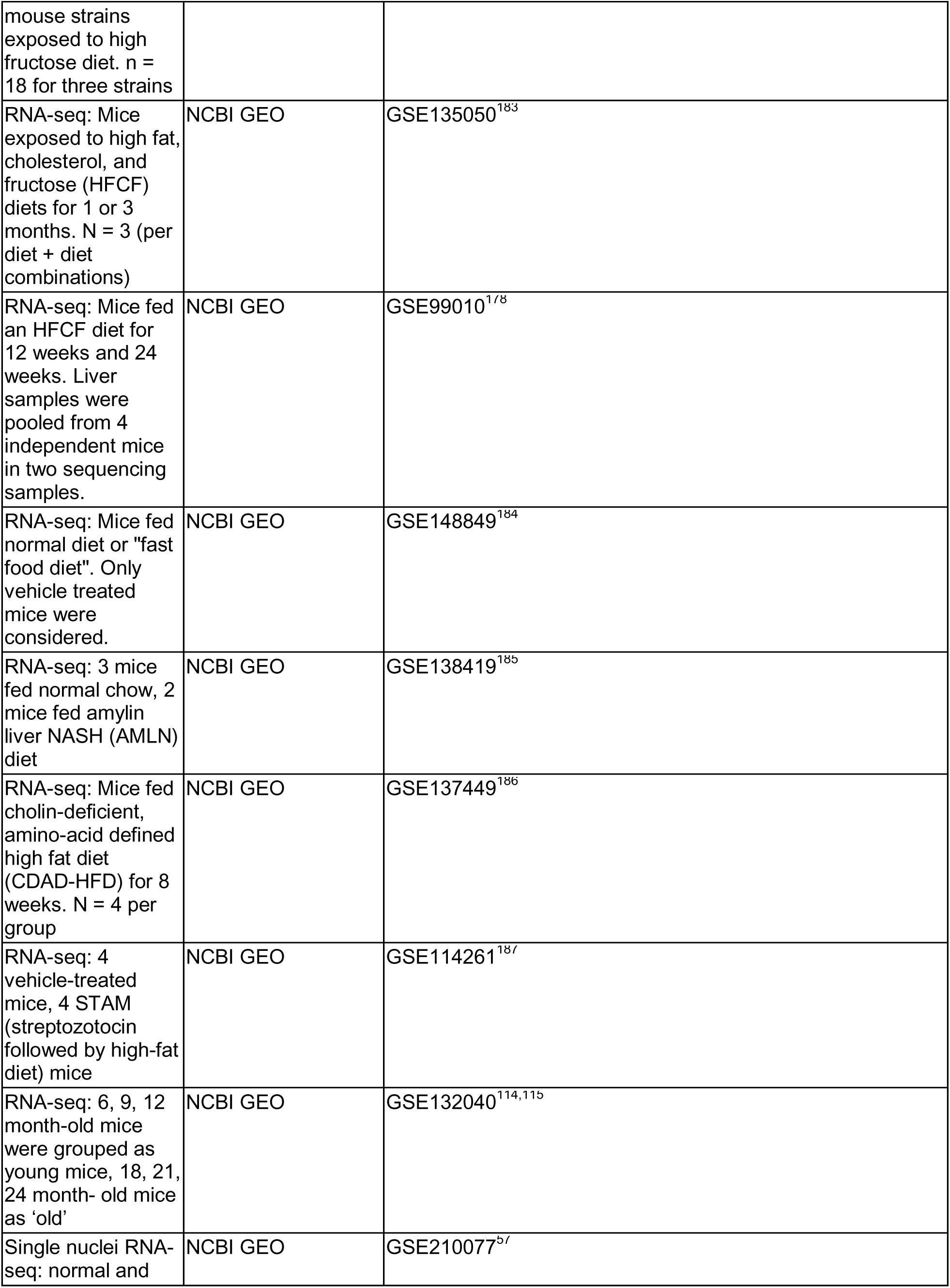
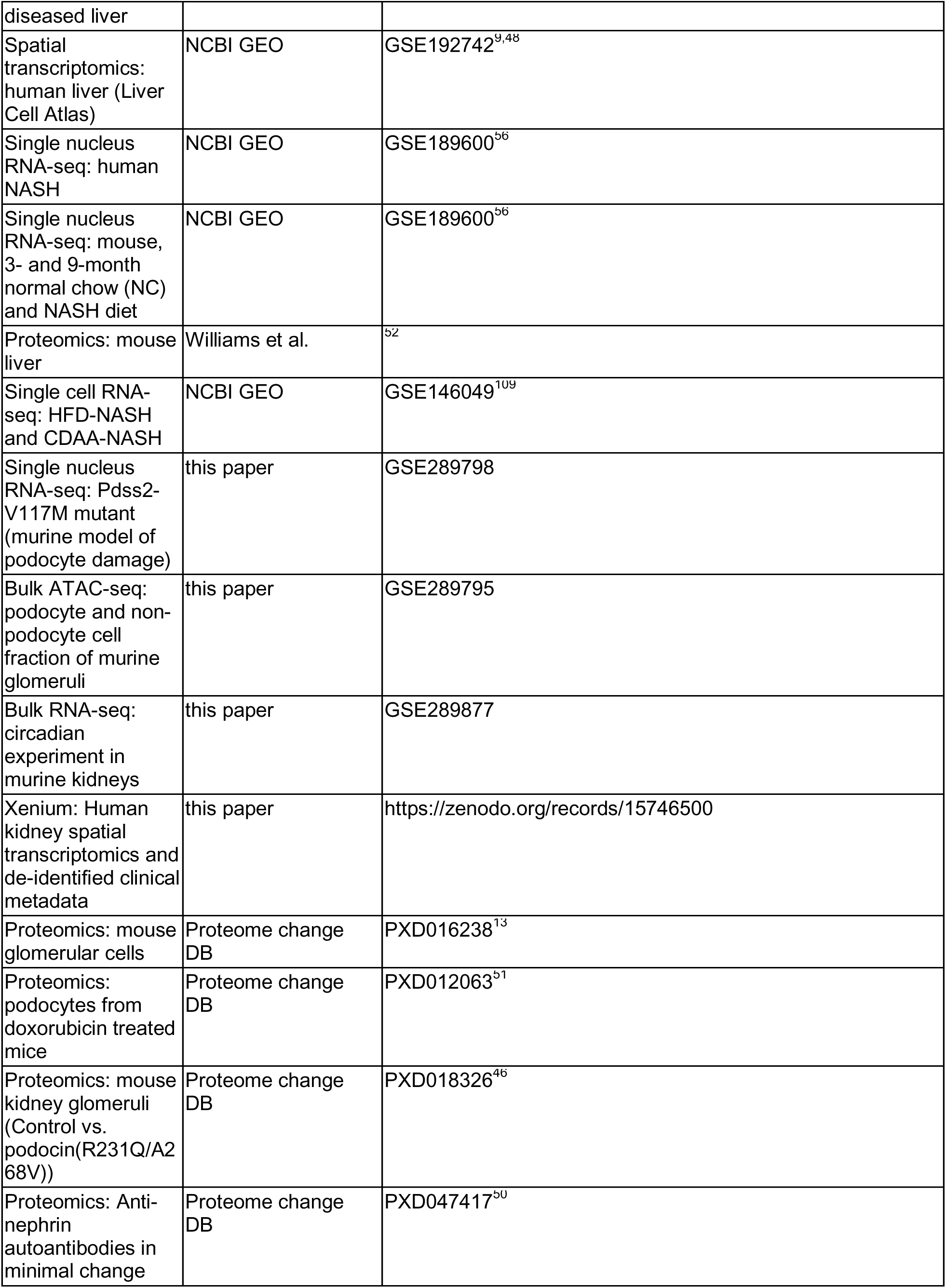
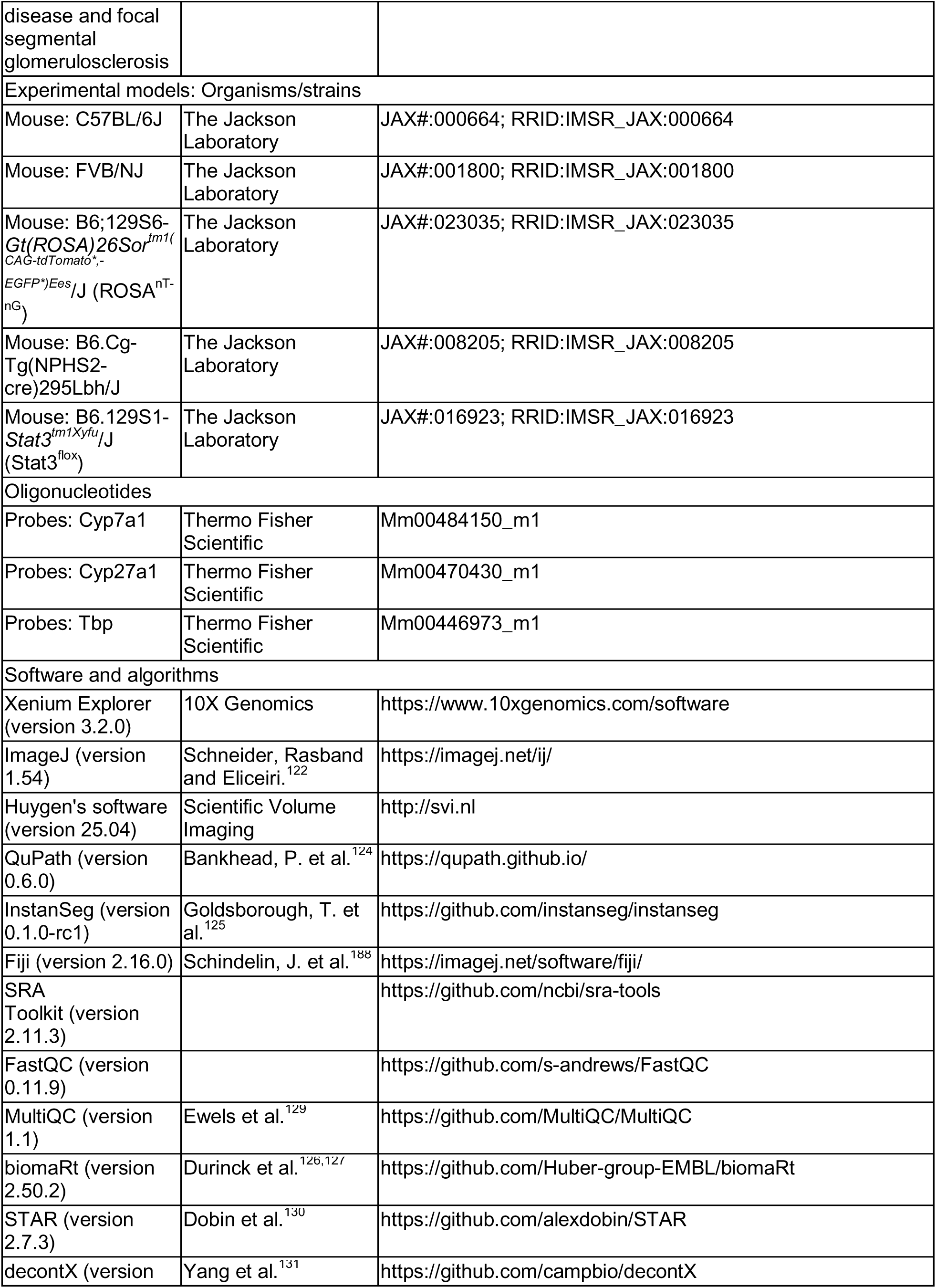
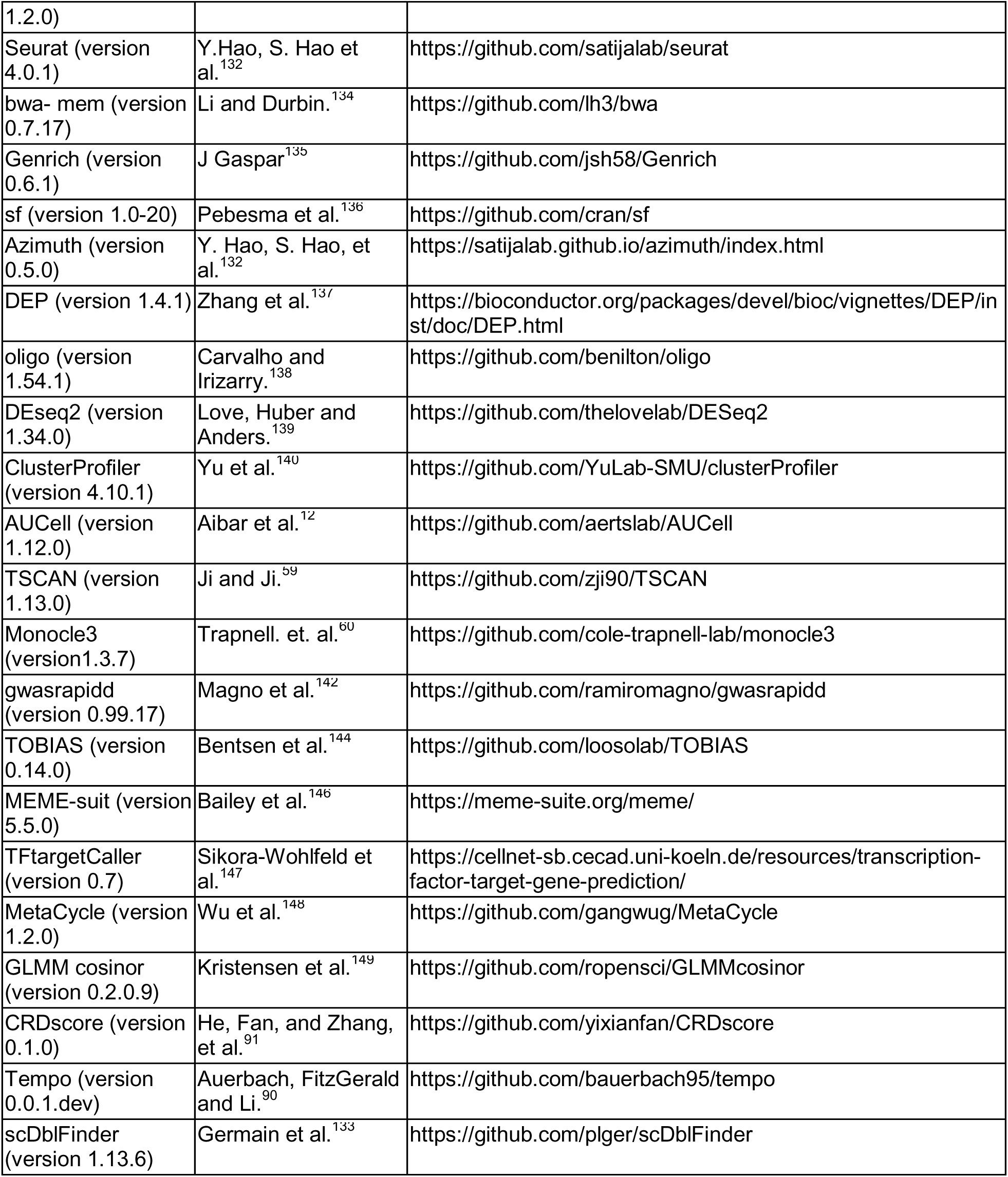

## Abbreviations

AFLD: alcoholic fatty liver disease
NAFLD: non-alcoholic fatty liver disease
NASH: non-alcoholic steatohepatitis
ASH: alcoholic steatohepatitis
CCl4: carbon tetrachloride
HCC: hepatocellular carcinoma
HFD: high fat diet
scRNA-seq: single cell RNA sequencing
snRNA-seq: single nucleus RNA sequencing
CKD: chronic kidney disease
FSGS: focal and segmental glomerulosclerosis
PDS: podocyte damage score
HDS: hepatocyte damage score

